# How *p53* stress memory could redirect JAK/STAT1 antiviral signalling: a model-based prediction

**DOI:** 10.64898/2026.07.09.737453

**Authors:** Francis Didier Tshianyi M.K., Rebecca Walo Omana, Apollinaire Ndondo Mboma, Didier Kumwimba Seya, Didier Gonze

## Abstract

Viral infection can co-activate interferon (IFN)–JAK/STAT1 signalling and the *p53* –Mdm2 stress-response pathway, two modules that jointly shape antiviral defence and cell-fate decisions. Here, we focus on viral infection contexts capable of inducing genotoxic stress associated with DNA double-strand breaks, thereby triggering oscillatory or sustained p53–Mdm2 dynamics. Whether *p53* acts merely as a parallel stress pathway, or actively reshapes how an activated JAK/STAT1 response is temporally decoded and functionally routed, remains unclear. We develop a coupled ordinary-differential-equation model linking an IFN-*γ*-centred JAK/STAT1 core, a *p53* –Mdm2 module, downstream antiviral and apoptotic effectors, and a coarse-grained viral-burden layer, with *p53* regulation placed downstream of STAT1 activation. We find that *p53* does not simply increase nuclear STAT1 availability; it redistributes the response towards DNA-bound STAT1 persistence, transcriptional memory and STAT1-driven feedback, producing a persistence–recovery trade-off in which prior *p53* stress prolongs the transcriptionally active STAT1 state but delays re-inducibility after repeated IFN stimulation. When IFN and *p53*-associated stress are both driven by viral burden, *p53* is not a uniform amplifier of host defence: *p53* preactivation strengthens the upstream memory layer, but downstream effectors buffer rather than mirror this priming. The model further separates antiviral-state engagement from realised viral control: strong effector activation does not guarantee suppression of poorly sensitive viral classes, whereas sensitive viral classes can be cleared before apoptosis. The origin of the stimulus also matters: exogenous IFN or *p53* stimulation allows us to assess the host’s intrinsic response capacity, whereas virus-induced IFN and *p53* stress remain coupled to viral persistence. Persistent viral burden thus emerges as the dynamical link between IFN induction, *p53* stress-memory, antiviral maintenance, viral control and the choice between JAK/STAT–IRF1-associated, *p53*-autonomous or dual apoptotic routing.

## 1. Introduction

Interferon signalling is a central component of antiviral and inflammatory host defence. Among interferon-induced pathways, the IFN-*γ*-centred JAK1/JAK2–STAT1 axis is of particular interest: it converts an extracellular cytokine signal into STAT1 phosphorylation, nuclear accumulation, DNA binding and transcriptional regulation. Unlike type-I-interferon signalling, which classically operates through STAT1–STAT2–IRF9 complexes, the IFN-*γ* response is organised around STAT1 activity and GAS/IRF1-associated programmes (Stark et al., 1998; Stark and Darnell, 2012; Schneider et al., 2014). This makes it a useful setting in which to ask how a cell interprets and propagates an activated STAT1 signal. Crucially, this functional output cannot be read from pathway activation alone: because STAT1 responses depend on nuclear trafficking, DNA binding, transcriptional propagation and feedback, their biological meaning is set not only by amplitude but also by timing, persistence, recovery after stimulation and transmission to downstream effectors.

At the network level, IFN-*γ* signalling begins when extracellular IFN-*γ* binds its receptor complex and activates the associated JAK1/JAK2 kinases. Activated receptors phosphorylate free cytosolic STAT1 monomers, allowing them to dimerise and translocate to the nucleus, where they accumulate and bind DNA at gamma-activated sequence (GAS)-associated regulatory elements. DNA-bound STAT1 then induces transcriptional programmes, including IRF1-dependent antiviral responses and SOCS1-mediated negative feedback. In the model, signal termination and attenuation are represented by three regulatory layers: SOCS1-mediated inhibition of receptor-proximal activation, cytosolic and nuclear PTP-mediated STAT1 de-phosphorylation, and PIAS1-associated regulation of nuclear STAT1 activity.

The tumour suppressor *p53* is best known as a regulator of cell-cycle arrest, DNA repair, senescence, metabolism and apoptosis (Michalak et al., 2005; Purvis et al., 2012; Lees et al., 2021), but it also operates in antiviral and inflammatory settings. In unstressed cells, *p53* activity is kept low largely through *Mdm2*-dependent degradation. Cellular stress can stabilise *p53*, producing transient, oscillatory or sustained responses whose temporal structure influences downstream cell-fate programmes (Purvis et al., 2012). Viral infection and interferon-associated stress can activate *p53*, and *p53* in turn can contribute to antiviral defence (Takaoka et al., 2003; Muñoz-Fontela et al., 2008; Rivas et al., 2010). More directly, *p53* can modulate IFN-*γ*-associated JAK/STAT signalling. In melanoma cells, IFN-*γ*-induced PD-L1 expression requires *p53*, which supports JAK2 expression and thereby sustains the JAK2/STAT1 response to IFN-*γ* (Thiem et al., 2019). Recent work further reports *p53*-dependent regulation of interferon signalling through SOCS1, STAT1 phosphorylation and gene-specific interferon-regulated outputs in contexts involving both type I and type II interferons (Będzińska et al., 2025). Together, these studies establish that *p53* can shape STAT1-associated signalling, yet the dynamical consequences of this influence remain poorly understood.

In particular, existing studies identify molecular and functional links between *p53* and interferon/STAT1 signalling but do not establish how a *p53*-dependent stress history reshapes the temporal decoding of a STAT1 response: whether *p53* acts merely as a parallel stress-response module, or whether it changes how an IFN-*γ*-centred JAK/STAT1 signal is stored, prolonged, recovered from, transmitted downstream and routed towards antiviral or apoptotic outcomes. Addressing this question requires a dynamical treatment in which signal duration, persistence, recovery, memory and downstream decoding are represented explicitly.

Previous models of JAK/STAT signalling have examined feedback regulation, desensitisation and signalling memory, while models of *p53* dynamics have analysed stress encoding and cell-fate control (Sarasin-Filipowicz et al., 2009; Kalliara et al., 2022; Purvis et al., 2012). To our knowledge, however, few if any models have addressed how a *p53*-dependent stress history reshapes the temporal organisation and functional interpretation of an IFN-*γ*-centred JAK/STAT1 antiviral response.

In earlier work, we studied the complementary direction of STAT1–*p53* coordination, asking how STAT1 modulates a *p53*-centred stress-response system and its downstream cell-fate programmes (Tshianyi et al., 2026). Here, we reverse this perspective and ask how *p53* - dependent stress signalling modifies a JAK/STAT1-centred response, and how that modulation propagates through antiviral effectors, viral burden and apoptotic competence.

To this end, we develop a coupled ordinary differential-equation model linking an IFN-*γ*-like JAK/STAT1 signalling core, a *p53* –*Mdm2* stress-response module, downstream antiviral and apoptotic effectors, and a coarse-grained viral-burden layer. The model is deliberately coarse-grained: rather than resolving pathogen sensing, interferon production or receptor-proximal regulation, we place the *p53*-dependent coupling downstream of STAT1 activation in order to isolate how *p53* stress history affects nuclear STAT1 processing, memory, effector transmission and apoptotic competence. This choice does not exclude receptor-proximal effects of *p53* reported experimentally, such as modulation of JAK2/STAT1 signalling or SOCS1-associated regulation (Thiem et al., 2019; Będzińska et al., 2025). Instead, it treats those mechanisms as complementary axes outside the present model, while focusing on the post-activation decoding of STAT1.

Viral classes enter as passive dynamical contexts rather than adaptive agents. Within the model, they deploy no active host-manipulation strategies and do not modify pathway parameters. They are characterised solely by fixed traits: their capacity to sustain an IFN-like STAT1-activating input, their capacity to induce *p53*-associated stress, and their sensitivity to antiviral effectors. This enables us to separate STAT1-pathway activation from downstream antiviral engagement, realised viral control and apoptotic routing. The model therefore provides a theoretical basis for determining when *p53* remains a parallel stress module and when it becomes part of the temporal decoding and functional routing of an IFN-*γ*-centred JAK/STAT1 antiviral response.

## 2. Model

In this section we present the assumptions used to build our model, the states variables used and its architecture.

### 2.1. Model overview

Our model formalises a JAK/STAT1-centred antiviral response coupled to a *p53* –Mdm2 stress-response module. It combines four functional layers: an IFN-driven JAK/STAT1 signalling module, a *p53* –Mdm2 module, a downstream antiviral-effector layer, and an intracellular viral-burden layer. The aim is not to describe each pathway at full molecular resolution, but to construct a mechanistic dynamical framework in which the effect of *p53* on the JAK/STAT1 response can be evaluated at the signalling, antiviral, viral-control, and apoptotic-decision levels. The complete equations, variables, parameters, and calibration choices are provided in Section S1 in the Supplementary Material and the state variables are provided in Table 1. Throughout the model, we use the term “gate” for a dimensionless regulatory function that makes a downstream process conditional on a specific dynamical context. A gate does not represent a single molecular reaction; it encodes whether a required condition, such as stress memory, co-activation, viral persistence or apoptotic competence, is sufficiently present to transmit or amplify a signal.

**Figure 1:**
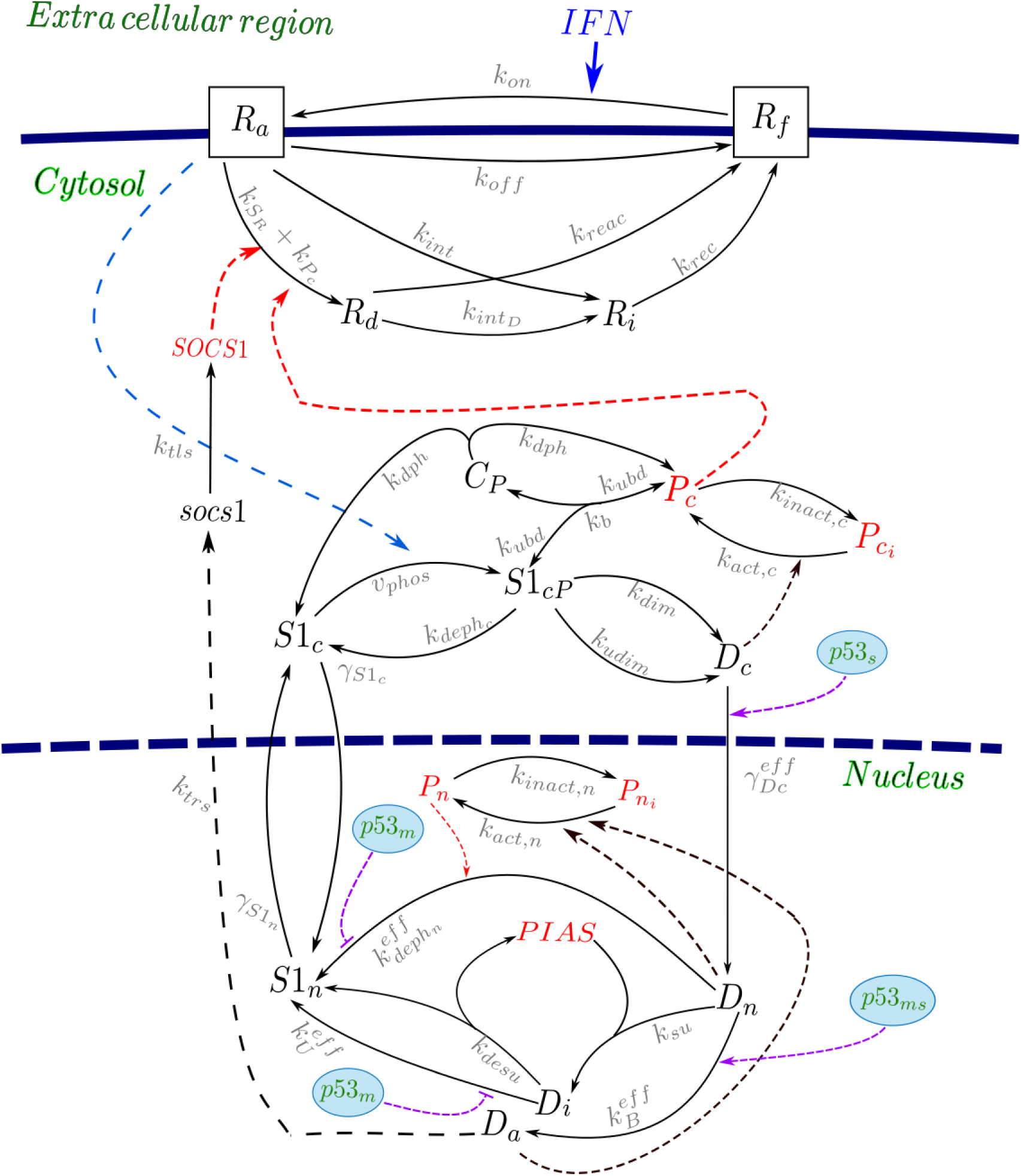
Schematic representation of the coupled JAK/STAT1–*p53* model. IFN binding activates receptor cycling and downstream STAT1 phosphorylation, dimerisation, and nuclear translocation. Nuclear STAT1 dimers (*D_n_*) bind DNA to form the transcriptionally active state (*D_a_*), which is regulated by PIAS1-mediated sequestration and phosphatase activity. Negative feedback is provided by SOCS1 and cytosolic phosphatases. *p53* modulates the pathway through memory (*p*53*_m_*) and instantaneous (*p*53*_s_*) components, affecting nuclear transport, DNA binding, and signal termination processes.

The *p53* –*Mdm2* oscillator used here is taken from our previous STAT1–*p53* model (Tshianyi et al., 2026). In that earlier model, STAT1 activity entered the *p53* module as a regulatory input. response.

**Table 1:** State variables and auxiliary inhibition factors of the coupled IFN–JAK/STAT–p53–effector–virus model.

| Symbol | Meaning |
| --- | --- |
| *Cytosolic receptor and STAT1 activation block* |  |
| $R_f$ | Free interferon receptor pool. |
| $R_a$ | Activated interferon-bound receptor pool. |
| $R_d$ | Deactivated receptor pool generated by phosphatase- and SOCS1-dependent inhibition. |
| $R_i$ | Internalised receptor pool. |
| $S1_c$ | Cytosolic unphosphorylated STAT1 monomer. |
| $S1_{cP}$ | Cytosolic phosphorylated STAT1 monomer. |
| $D_c$ | Cytosolic phosphorylated STAT1 dimer. |
| $C_P$ | Cytosolic phosphatase-STAT1 trapping complex. |
| $soc1$ | SOCS1 mRNA. |
| $SOCS1$ | SOCS1 protein acting as a negative regulator of receptor signalling. |
| $PTP_c$ | Active cytosolic phosphatase pool. |
| *Nuclear STAT1 and p53-coupling block* |  |
| $S1_n$ | Nuclear unphosphorylated STAT1 monomer. |
| $D_n$ | Nuclear phosphorylated STAT1 dimer. |
| $D_a$ | DNA-bound active STAT1 dimer. |
| $D_i$ | PIAS1-sequestered inactive STAT1 dimer. |
| $PTP_n$ | Active nuclear phosphatase pool. |
| $PIAS1_n$ | Nuclear PIAS1 pool mediating STAT1 sequestration. |
| $P_s$ | p53 memory variable controlling memory-dependent JAK/STAT modulation. |
| *p53-Mdm2 stress-response block* |  |
| $S_{p53}$ | Viral-stress signal driving p53 activation. |
| $p53$ | Nuclear p53 protein level. |
| $Mdm2_{RNA}$ | Mdm2 mRNA induced by p53. |
| $Mdm2$ | Mdm2 protein mediating p53 degradation. |
| *Endogenous IFN sensing block* |  |
| $S_v$ | Viral sensing variable induced by intracellular viral genome burden. |
| $E_v$ | Delayed exhaustion or adaptation variable limiting endogenous IFN sensing. |
| $IFN_{int}$ | Endogenously produced interferon signal. |
| $IFN_{ext}$ | Externally supplied interferon input. |
| $IFN$ | Total interferon signal combining external and endogenous sources. |
| *Effector and apoptotic commitment block* |  |
| $A_D$ | Integrated memory of DNA-bound STAT1 activity. |
| $IRF1$ | IRF1 transcription factor induced by STAT1 activity. |
| $M_{IRF1}$ | IRF1 memory variable. |
| $A_{IRF1}$ | Delayed IRF1-associated apoptotic activation variable. |
| $P_1$ | Antiviral effector associated with inhibition of viral genome replication. |
| $P_2$ | Antiviral effector associated with inhibition of viral protein production. |
| $P_3$ | Antiviral effector associated with enhanced viral protein loss. |
| $P_4$ | JAK/STAT-IRF1-associated apoptotic effector potentiated by p53. |
| $P_5$ | p53-autonomous apoptotic effector. |
| *Viral dynamics block* |  |
| $V_g$ | Intracellular viral genome burden. |
| $V_p$ | Intracellular viral protein burden. |
| $I_g$ | Antiviral inhibition factor acting on viral genome replication. |
| $I_p$ | Antiviral inhibition factor acting on viral protein production. |
| $I_{loss}$ | Antiviral enhancement factor increasing viral protein loss. |

A central variable in this coupling is the *p53*-dependent stress-memory state *P_s_*. This variable is not intended to represent a single molecular species, but a coarse-grained memory of prior *p53*-associated stress. It acts as a low-pass filtered stress-history variable: it accumulates when the *p53* module is active and relaxes when the stress signal decreases. This allows the model to distinguish an acute *p53* response from an established *p53*-dependent stress state. The introduction of *P_s_*is motivated by the fact that cellular decisions can depend not only on the instantaneous level of a signalling molecule, but also on the duration, persistence and temporal history of pathway activation. Distinct temporal profiles of *p53* activity have been associated with different downstream cell-fate programmes (Purvis et al., 2012), and prior IFN–JAK/STAT stimulation can leave signalling or transcriptional memory, including long-lasting desensitisation and IFN-*γ*-induced transcriptional memory (Kalliara et al., 2022; Tehrani et al., 2023). Thus, *P_s_* is introduced as a phenomenological representation of stress history rather than as evidence for a unique molecular carrier of *p53* memory. The JAK/STAT1 module describes receptor activation, STAT1 phosphorylation, dimerisation, nuclear import, DNA binding, and negative regulation by SOCS1, PIAS1, and phosphatases. Its main transcriptionally active output is the DNA-bound STAT1 state *D_a_*, which drives IRF1-associated responses and downstream effectors. The effect of *p53* is introduced through *P_s_* and a set of phenomenological gates that determine how this stress-memory state modifies nuclear decoding, downstream effector transmission, and apoptotic competence.

This *p53*-dependent regulatory layer contains five gates. The memory gate Φ_mem_(*P_s_*) represents the effect of accumulated *p53*-dependent stress history on DNA-bound STAT1 persistence by modulating DNA binding, DNA unbinding and nuclear dephosphorylation through effective rates such as 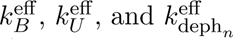. The synchronisation gate Φ_sync_ captures acute co-activation between STAT1 and *p53*, affecting nuclear import and early DNA-bound STAT1 activation. The effector-modulation gate Φ*_p_*_53_(*p*53) changes the transmission of STAT1–IRF1 activity to antiviral outputs, whereas Φ_apop_(*p*53) represents *p53*-dependent apoptotic competence. Finally, the viral-burden gate Φ*_V_* _4_(*V_g_*) ensures that the *P*_4_ route remains explicitly conditional on persistent intracellular viral burden.

These gates are phenomenological functions rather than single molecular reactions. They encode the hypothesis that *p53*-dependent stress does not primarily alter entry of the IFN signal into the JAK/STAT1 pathway, but reshapes how an activated STAT1 signal is stored, stabilised, transmitted to effectors and interpreted in an infection-dependent apoptotic context. This construction is motivated by reported interactions between STAT1 and *p53*, and by broader evidence that *p53* intersects with interferon-dependent signalling (Townsend et al., 2004; Youlyouz-Marfak et al., 2008; Będzińska et al., 2025). Because these gates aggregate processes for which direct quantitative constraints are unavailable, their individual parameters are not identifiable separately at the coarse-grained resolution of the model. The nominal calibration is therefore used as a representative parameter set. Qualitative conclusions are retained only when they persist across broad, biologically admissible variation of the memory, synchronisation, effector-modulation, apoptotic-competence and viral-burden-gating parameters (Supplementary Section S8).

The total IFN input combines an externally prescribed IFN signal and an endogenous virus-induced IFN signal. This formulation allows the same JAK/STAT1 module to be studied under externally imposed stimulation, infection-driven stimulation or mixed IFN-source conditions. In this framework, intracellular viral genomes *V_g_* are converted phenomenologically into two host-response inputs: endogenous IFN production and *p53*-associated checkpoint stress. Viral burden is therefore the infection-associated context in which JAK/STAT1 activation and *p53*-dependent stress signalling are jointly generated. This choice is biologically motivated by studies linking viral infection, IFN signalling and *p53*-dependent antiviral responses (Takaoka et al., 2003; Muñoz-Fontela et al., 2008; Rivas et al., 2010).

The viral-burden layer contains intracellular viral genomes *V_g_* and viral particles *V_p_*. Passive viral classes are defined by fixed traits: their ability to induce endogenous IFN, their ability to induce *p53*-associated stress, and their sensitivity to antiviral effectors. They are not treated as active agents with explicit adaptive strategies. Instead, they provide infection contexts in which the same host signalling architecture can be tested, ranging from efficient viral restriction to persistent viral burden.

The downstream effector layer contains three antiviral outputs, *P*_1_, *P*_2_, and *P*_3_, interpreted as coarse-grained functional axes rather than individual molecular species. They summarise antiviral restriction programmes downstream of STAT1–IRF1 activity and modulated by *p53*-dependent stress through Φ*_p_*_53_(*p*53) and Φ_mem_(*P_s_*). Functionally, *P*_1_ inhibits viral genome replication, *P*_2_ inhibits viral-particle production, and *P*_3_ enhances viral-particle loss. Their sustained activation defines the antiviral-state score *L*_AV_. This coarse-grained representation is consistent with the diversity of interferon-inducible restriction mechanisms acting on viral replication, translation, intracellular permissiveness and the infected-cell antiviral state (MacMicking, 2012; Schneider et al., 2014; Schoggins et al., 2011; Tretina et al., 2019).

The apoptotic layer separates two routes. The first route, *P*_4_, represents a JAK/STAT–IRF1-associated danger route. It is driven by sustained IRF1-associated signalling, explicitly gated by persistent viral burden through Φ*_V_* _4_(*V_g_*), and potentiated by *p53*-dependent apoptotic competence through Φ_apop_(*p*53). Thus, *P*_4_ is not activated by JAK/STAT1 signalling alone; it requires a persistent viral-burden context and sustained IRF1-associated activation, while *p53* increases its apoptotic competence. The second route, *P*_5_, represents a more direct *p53*-autonomous apoptotic programme driven by high and persistent *p53* activity. These two routes allow the model to distinguish JAK/STAT-associated apoptotic commitment, *p53*-autonomous commitment and dual-route commitment. The biological interpretation is consistent with the known involvement of IFN/STAT1/IRF1-linked mechanisms in apoptosis and with canonical *p53*-regulated apoptotic programmes (Fulda and Debatin, 2002; Tamura et al., 1995; Gao et al., 2010; Michalak et al., 2005).

These downstream variables are interpreted using the common readout framework defined below, which separates antiviral-state engagement, realised viral control and apoptotic-route selection.

### 2.2. The model architecture

The complete model is defined by six coupled ordinary-differential-equation blocks, denoted (*IFN*), (*C*), (*N*), (*p*53), (*E*), and (*V*). Each block is introduced with its associated auxiliary functions.

*Block* (*IFN*) *– composite IFN input and virus-induced endogenous IFN production.* The total IFN signal entering the JAK/STAT1 module is defined as the weighted sum of an externally prescribed IFN input and an endogenous virus-induced IFN input:

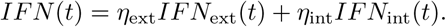

The external component *IFN*_ext_(*t*) represents an imposed IFN stimulus, whereas *IFN*_int_(*t*) is produced dynamically from intracellular viral burden. The weights *η*_ext_ and *η*_int_ set the relative contribution of the two IFN sources, allowing externally driven, virus-induced, or mixed IFN-source conditions to be studied within the same model.

The externally imposed IFN input is prescribed as

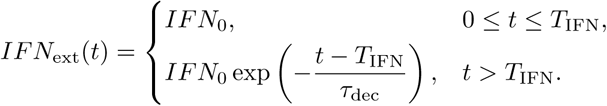

Thus, external IFN is maintained at amplitude *IFN*_0_ during the stimulation window and then decays exponentially after withdrawal.

The endogenous IFN component is generated by intracellular viral genomes through a sensing variable *S_v_*, an exhaustion variable *E_v_*, and the virus-induced IFN output *IFN*_int_. Viral genomes activate *S_v_* with strength *χ*_IFN_, while *E_v_* limits sustained sensing under persistent viral burden. The endogenous IFN dynamics are

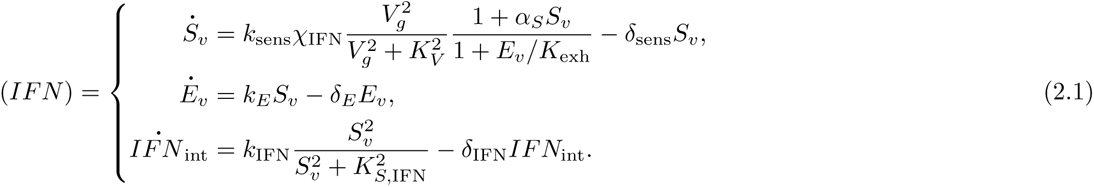

*Blocks* (*C*) *and* (*N*) *– IFN receptor–JAK/STAT1 pathway.* The IFN receptor–JAK/STAT1 pathway is represented by a cytosolic/receptor layer (*C*) and a nuclear STAT1-processing layer (*N*). Block (*C*) describes receptor activation, receptor desensitisation/internalisation, cytosolic STAT1 phosphorylation and dimerisation, SOCS1 feedback, and cytosolic PTP activity.

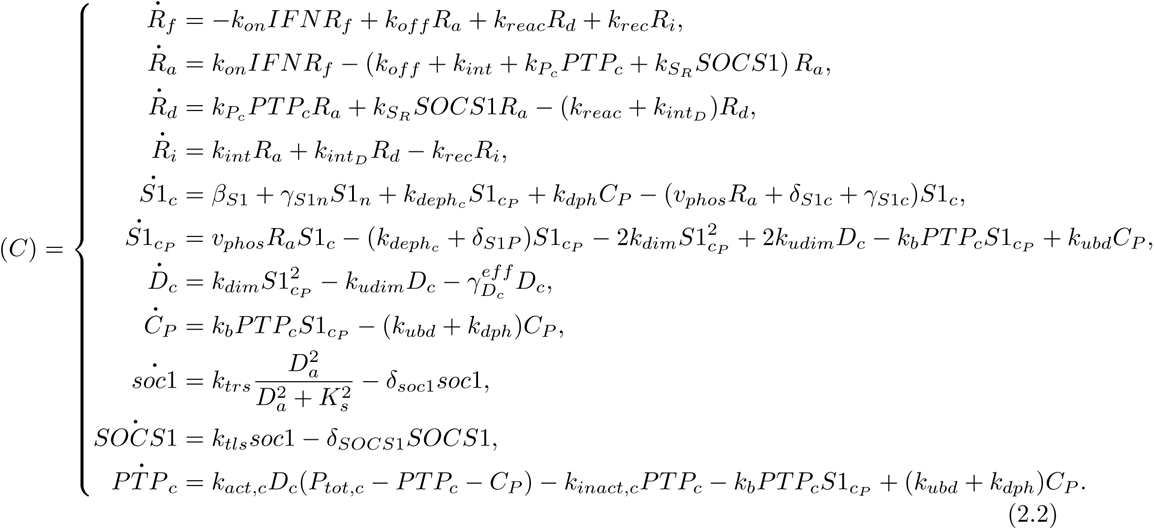

The auxiliary functions associated with block (*C*) are

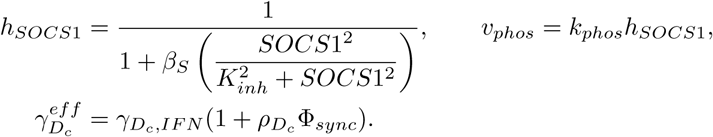

Block (*N*) describes nuclear STAT1 processing, SUMO-associated STAT1 regulation, nuclear PTP activity, PIAS1 dynamics, and the *p53*-dependent memory variable *P_s_*.

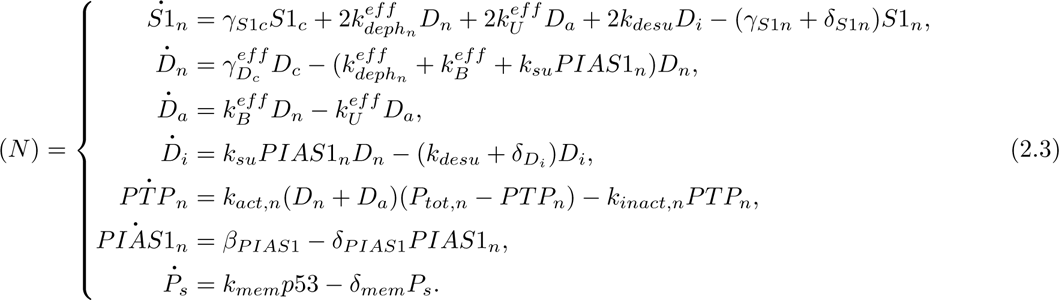

The auxiliary functions associated with block (*N*) are

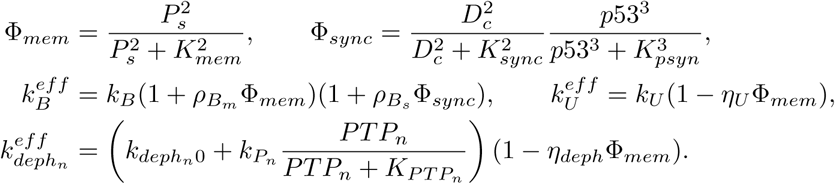

*Block* (*p*53) *– virus-induced p53–Mdm2 stress-response module.* This block describes virus-induced *p53* stress activation and the *p53* –*Mdm2* negative-feedback loop.

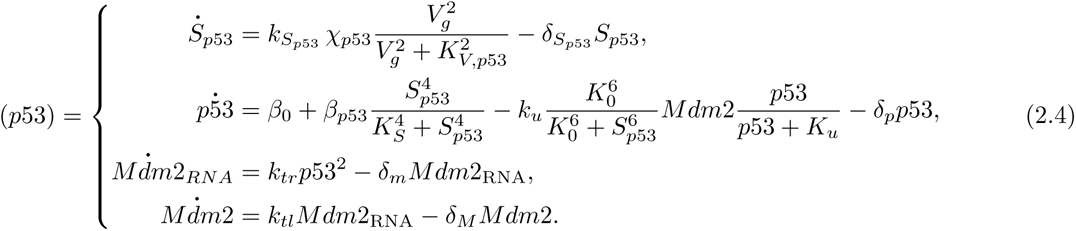

The interaction between the JAK/STAT1 pathway and the *p53* pathway is described in Figure 1.

*Block* (*E*) *– antiviral and apoptotic effector layer.* This block describes five downstream effector axes generated by the IFN-*γ*-centred JAK/STAT1–IRF1–*p53* network. The antiviral variables *P*_1_, *P*_2_, and *P*_3_ are coarse-grained functional outputs rather than single molecular species, but they represent distinct layers of antiviral restriction. *P*_1_ is the most IFN-sensitive effector axis: it is driven directly by DNA-bound STAT1 *D_a_* and represents rapidly inducible genome-restriction mechanisms that can respond to low IFN/JAK–STAT1 activity. *P*_2_ is driven by the accumulated STAT1 activity *A_D_*, and therefore represents a delayed restriction layer acting on viral-particle production. *P*_3_ is driven by IRF1 memory *M*_IRF1_, making it a later and more integrated antiviral layer that enhances viral-particle loss or reduces viral-particle persistence. Thus, *P*_1_, *P*_2_, and *P*_3_ do not simply duplicate one generic antiviral output; they encode increasing requirements for signal accumulation and transcriptional memory.

Biologically, these axes can be related to IFN-*γ*–STAT1/IRF1-induced restriction programmes, including GBP-family GTPases, IDO1, NOS2, RSAD2/viperin, ISG20, TRIM-family restriction factors, BST2/tetherin and autophagy- or ubiquitin-associated antiviral mechanisms (MacMicking, 2012; Schneider et al., 2014; Tretina et al., 2019). These examples are not meant as one-to-one identities for *P*_1_, *P*_2_, or *P*_3_, but as biological anchors for the three functional layers.

The apoptotic outputs are separated into two route-specific axes. *P*_4_ represents a JAK/STAT–IRF1-associated apoptotic danger route linked to sustained STAT1/IRF1 activity and explicitly gated by persistent viral burden through Φ*_V_* _4_(*V_g_*). It can be related to IRF1-associated death-receptor or caspase-linked programmes, including FAS/CD95, TNFSF10/TRAIL, CASP8 and related mediators. *P*_5_ represents a more direct *p53*-autonomous apoptotic axis, consistent with canonical *p53*-regulated effectors such as PUMA/BBC3, NOXA/PMAIP1, BAX and APAF1. Thus, the separation of *P*_1_–*P*_5_ reflects biologically distinct antiviral and apoptotic layers while remaining coarse-grained at the level of the model.

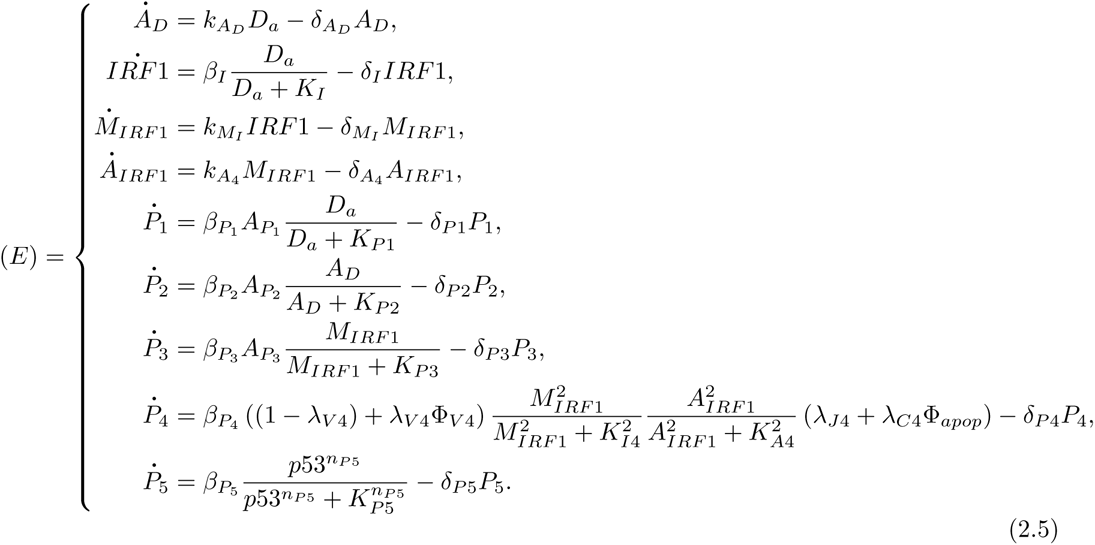

The auxiliary functions associated with this block are

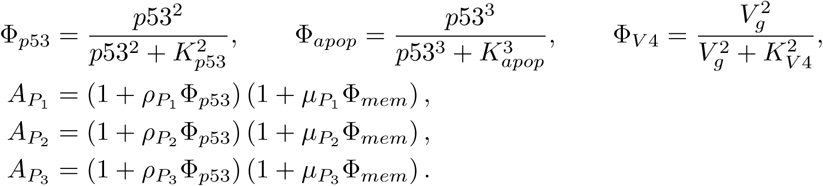

*Block* (*V*) *– viral replication and effector-mediated viral control.* This block describes intracellular viral genome replication and viral-particle production under antiviral effector pressure.

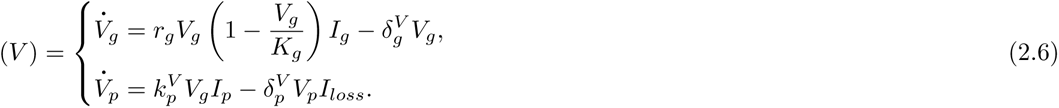

The auxiliary functions associated with this block are

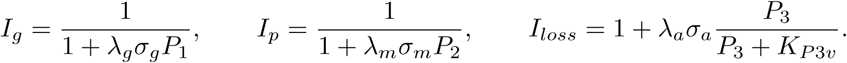

Together, the six blocks define a coupled IFN–JAK/STAT1–*p53* –virus system. Endogenous and external IFN inputs activate the receptor–STAT1 module; *p53*-dependent memory and synchronisation modify STAT1 nuclear processing and effector transmission; antiviral effectors feed back on viral genome replication and particle production; and persistent viral burden contributes to both endogenous IFN induction and *P*_4_-route activation. The complete architecture is summarised in Figure 2.

**Figure 2:**
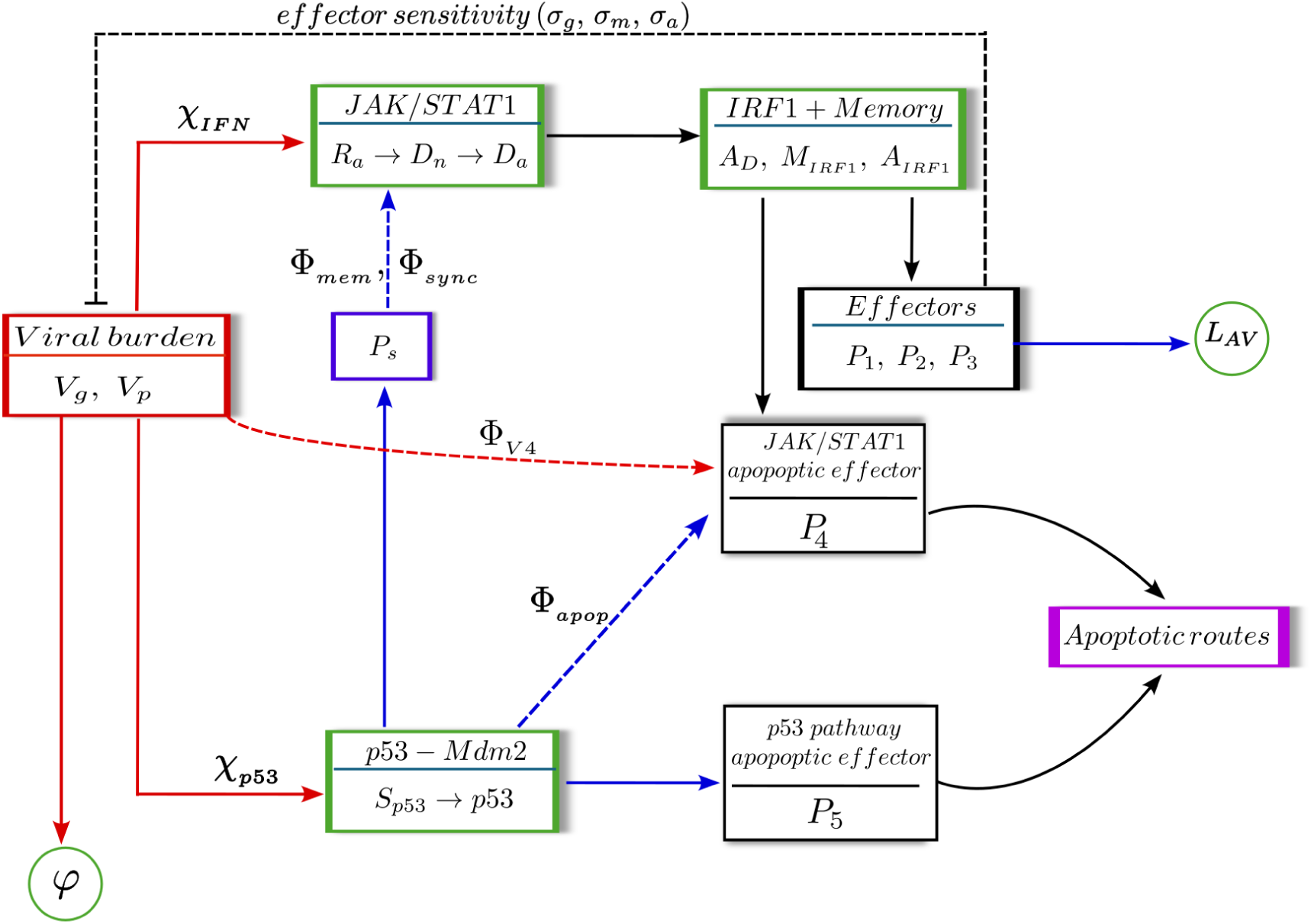
Schematic representation of the coupled IFN–JAK/STAT1–*p53* –effector–virus model. Viral burden, represented by *V_g_* and *V_p_*, provides two infection-dependent inputs: endogenous IFN induction through *χ*_IFN_, and checkpoint-stress induction through *χ_p_*_53_. Activated STAT1 is routed through nuclear STAT1 activity and the IRF1/memory layer, generating antiviral effectors *P*_1_, *P*_2_, and *P*_3_, whose sustained activation defines the antiviral-state score *L*_AV_. These effectors feed back on viral burden with class-dependent sensitivities (*σ_g_, σ_m_, σ_a_*), thereby determining the viral-control score *φ*. The *p53*-dependent memory variable *P_s_* modulates JAK/STAT1 signalling through Φ_mem_ and Φ_sync_. Apoptotic routing is represented by the JAK/STAT–IRF1-associated route *P*_4_, explicitly gated by persistent viral burden through Φ*_V_* _4_(*V_g_*) and potentiated by Φ_apop_(*p*53), and by the *p53*-autonomous route *P*_5_. Sustained activation of *P*_4_, *P*_5_, or both defines JAK/STAT-associated, *p53*-autonomous, or dual apoptotic routing.

*Prescribed p53 stress regimes.* The *p53* –*Mdm2* stress-response module is inherited from our previous STAT1–*p53* modelling study (Tshianyi et al., 2026). In that study, the stress amplitude *S*_0_ organised the *p53* response into distinct dynamical regimes rather than a single monotonic response curve. Figure 3 shows the reference regimes used here: near-basal, damped, sustained-oscillatory and plateau-like *p53* dynamics.

In the coupled model, these regimes are used as prescribed upstream *p53* backgrounds to test how *p53* reshapes IFN–JAK/STAT1 decoding, antiviral effector transmission and apoptotic routing. Unless otherwise stated, the JAK/STAT1-core condition corresponds to the absence of active *p53* stress, the sustained-oscillatory regime to *S*_0_ = 10, the upper-moderate damped regime to *S*_0_ = 17, and the plateau-like regime to *S*_0_ = 24.

When *p53* stress is induced by viral burden rather than prescribed externally, the virus-induced stress signal is treated as an effective input to the same *p53* –*Mdm2* module. This allows prescribed and virus-induced *p53* responses to be interpreted within a common stress-regime framework. Figure S6 illustrates how viral burden is converted into an effective stress input and mapped onto the corresponding *p53* dynamical regime.

### 2.3. Readout framework: antiviral state L*_AV_*, viral outcome φ, and apoptotic routing APO-J/P/D

We now move from signalling-level responses to simulations in which JAK/STAT1–*p53* activity is propagated to antiviral effectors, viral burden and apoptotic routes. To interpret these downstream simulations consistently, we define a common readout framework for three quantities: antiviral-state engagement, functional viral outcome and apoptotic routing. This framework is used in both prescribed-input simulations and virus-induced scenarios.

**Figure 3:**
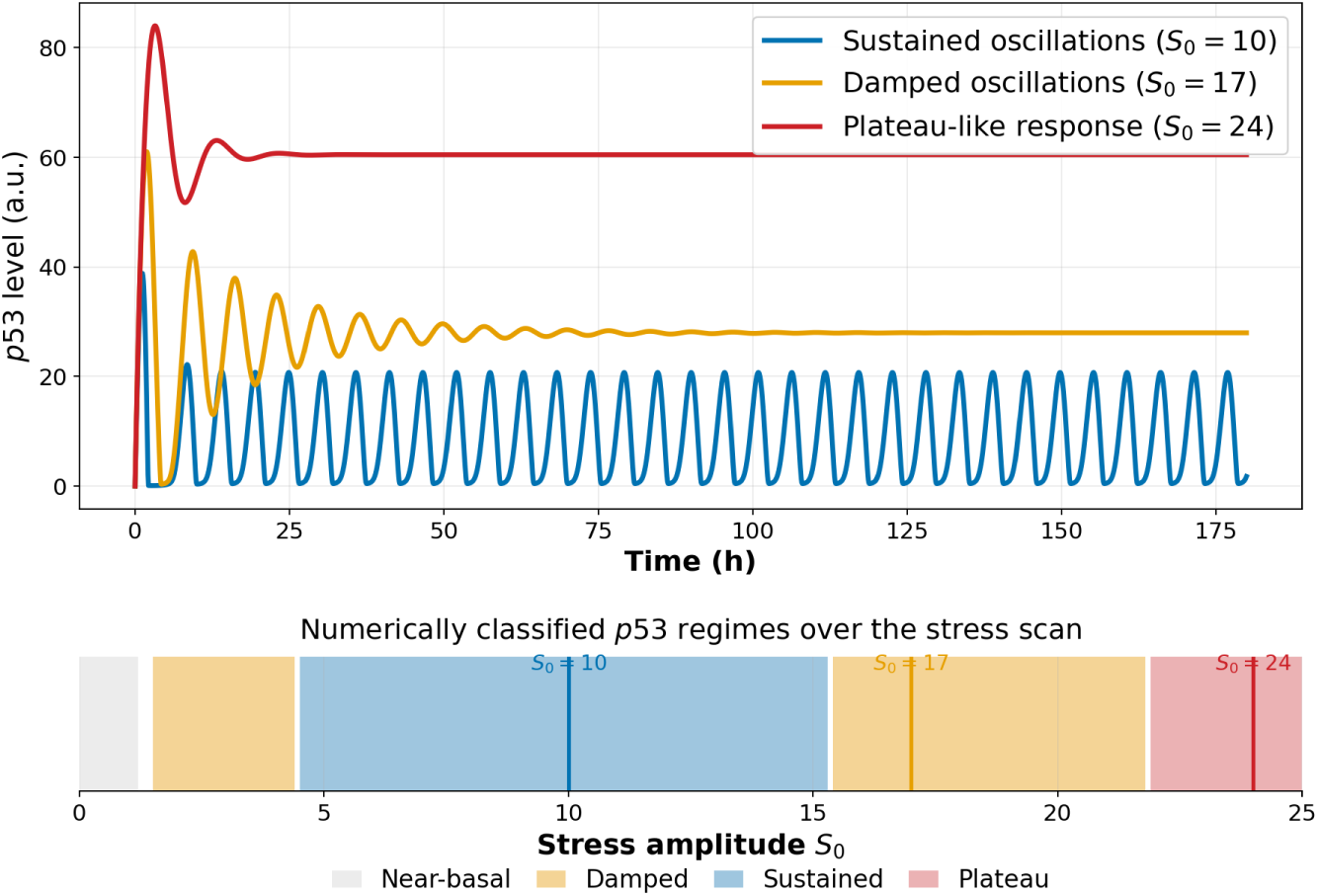
Standalone *p53* –Mdm2 stress-response regimes used as upstream *p53* backgrounds in the coupled antiviral-response model. The upper panel shows representative *p53* trajectories for *S*_0_ = 10 sustained oscillations, *S*_0_ = 17 damped oscillations, and *S*_0_ = 24 plateau-like activation. The lower panel shows the numerical classification of the stress-amplitude scan *S*_0_ ∈ [0, 25], with coloured bands denoting near-basal, damped, sustained and plateau-like regimes. Vertical lines indicate the reference stress values used in the simulations.

The purpose of this framework is to avoid treating signalling activation, effector engagement, viral control and apoptotic commitment as equivalent outcomes. A cell may activate a strong antiviral programme without clearing the virus, and apoptosis may occur through a JAK/STAT–IRF1-associated route, a *p53*-autonomous route, or both.

For each output *P_i_*, with *i* = 1*, . . .,* 5, sustained activation was defined by a threshold-and-duration rule. The output *P_i_* was considered active when it remained above its threshold *θ_P_* for at least a prescribed duration *τ_i_*:

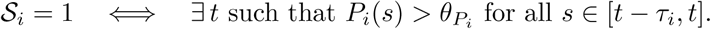

Otherwise, S*_i_*= 0.

The antiviral state was defined from the three antiviral effectors *P*_1_, *P*_2_, and *P*_3_:

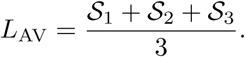

Thus, *L*_AV_ = 0 denotes a permissive state, 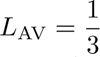 weak restriction, 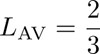 moderate restriction, and *L*_AV_ = 1 strong restriction. Apoptotic commitment was assessed separately from antiviral-state engagement. The variable *P*_4_ represents the JAK/STAT–IRF1-associated apoptotic route, whereas *P*_5_ represents the *p53*-autonomous apoptotic route. Apoptosis was assigned when either route satisfied its sustained-activation criterion:

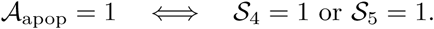

The route was classified as APO-J when only *P*_4_ was active, APO-P when only *P*_5_ was active, and APO-D when both routes were active.

Functional viral control was quantified from the late viral burden rather than from effector activation alone. For viral genomes *V_g_* and viral particles *V_p_*, we computed the mean level over the late observation window [96, 120] h:

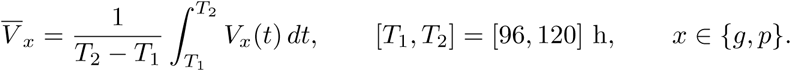

These late values were compared with the corresponding untreated reference scales,

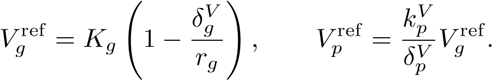

Genome-level and particle-level suppression scores were defined as

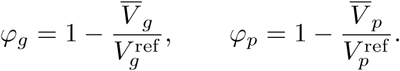

The global viral-control score was then defined conservatively as

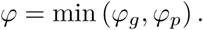

With this choice, strong viral control is assigned only when both viral genomes and viral particles are suppressed.

The viral outcome was classified from *φ*. The virus was considered cleared when *φ* ≥ 0.97 or when the late viral genome level was below the viral floor, controlled when 0.70 ≤ *φ <* 0.97, restricted when 0.30 ≤ *φ <* 0.70, and evaded when *φ <* 0.30. Therefore, *L*_AV_ measures activation of the host antiviral programme, whereas *φ* measures the realised effect of that programme on viral burden.

The thresholds and duration windows used for the main simulations were

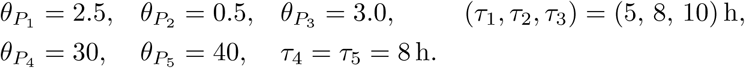

A threshold-robustness analysis for the apoptotic routes is provided in the Supplementary Material (Section S6.5, Supplementary Figs. S7–S9).

## 3. How *p53* reshapes post-phosphorylation JAK/STAT1 decoding of IFN stimulation

Given this downstream-coupling architecture, we next analyse how *p53*-dependent gates reshape the nuclear decoding of an IFN-induced STAT1 signal. To do so, we analysed the JAK/STAT1 response under various *p53*-dependent stress regimes, ranging from no *p53* activity to oscillatory, damped, and plateau-like *p53* profiles. We focused on three levels of the nuclear STAT1 response: the nuclear phosphorylated dimer pool *D_n_*(*t*), the DNA-bound active STAT1 state *D_a_*(*t*), and the transcriptional memory variable *A_D_*(*t*). We also tracked SOCS1 as the main STAT1-driven negative-feedback output. This decomposition separates nuclear STAT1 availability, transcriptionally active STAT1 binding, downstream memory, and feedback activation.

Biologically, this distinction is motivated by the fact that IFN signalling does not only generate rapid STAT1 activation, but also establishes transcriptional states whose duration and persistence can influence downstream cellular programmes (Stark et al., 1998; Stark and Darnell, 2012; Schneider et al., 2014). Moreover, *p53* has been implicated in antiviral immunity and in functional crosstalk with STAT1-dependent stress and apoptotic responses (Takaoka et al., 2003; Muñoz-Fontela et al., 2008; Rivas et al., 2010; Townsend et al., 2004). The mathematical properties of the model are provided in Section S2.1. The scalar response metrics used to compare the different simulation protocols quantify amplitude, integration, timing, and post-peak persistence; their definitions are provided in Section S3.

### 3.1. p53 redistributes post-phosphorylation STAT1 signalling towards the transcriptional layer

Because the gates act downstream of receptor activation, a direct effect on DNA-bound STAT1 dynamics is partly expected from the model structure: the memory gate Φ_mem_ modulates DNA binding through 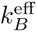, DNA unbinding through 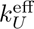, and nuclear dephosphorylation through 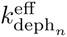, while the synchronisation gate Φ_sync_ modulates nuclear import through 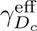. The question is therefore not whether these gates affect the nuclear layer, but how their combined action *redistributes* the IFN-induced STAT1 response between nuclear STAT1 availability, DNA-bound persistence, transcriptional memory, and feedback activation.

**Figure 4:**
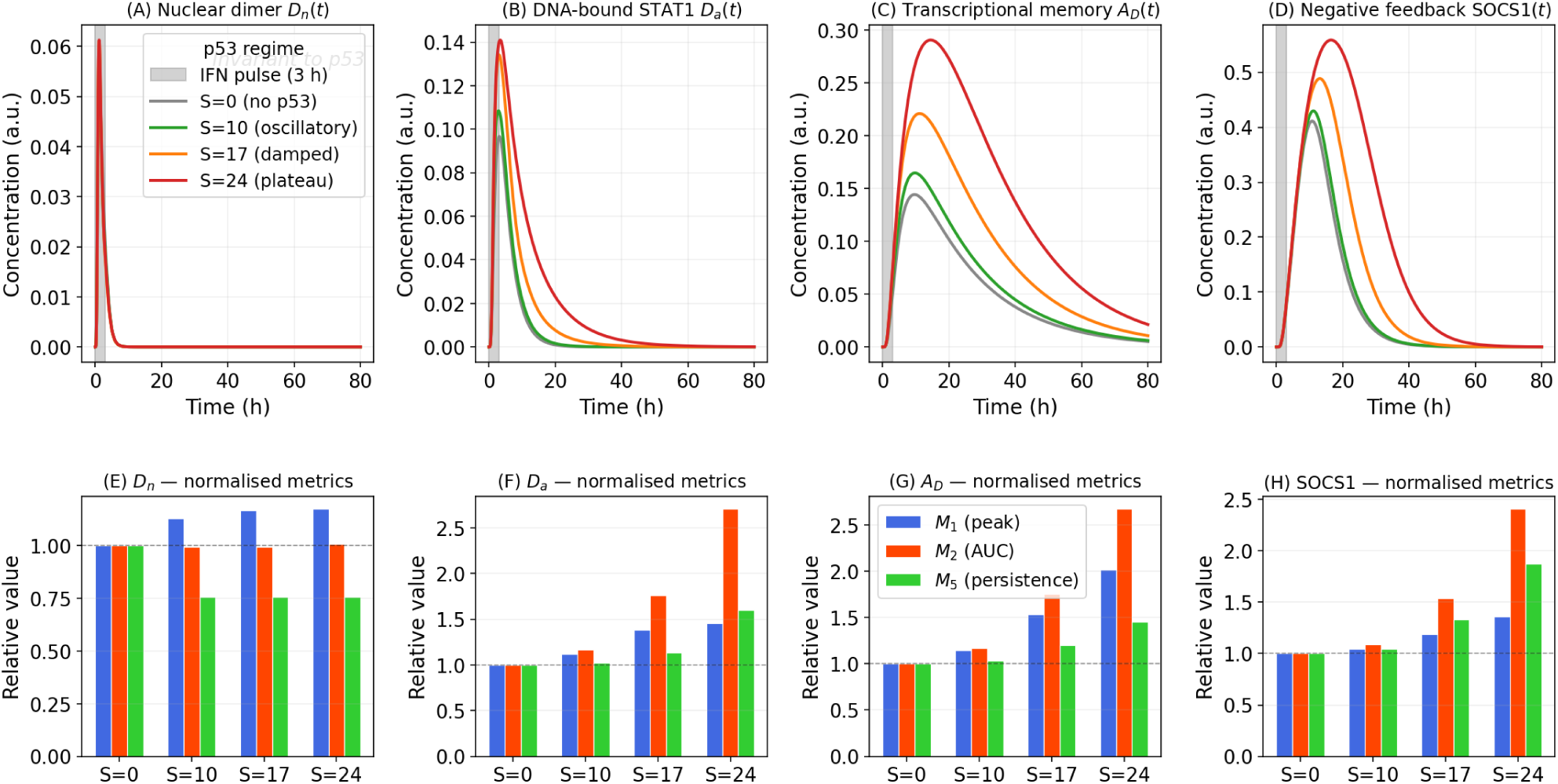
Time courses and relative response metrics of nuclear STAT1 signalling under different *p53* regimes following a transient IFN stimulation. The IFN=50 input is applied for 3 h, from *t* = 0 to *t* = 3 h, as indicated by the grey shaded region in panels (A)–(D). Panels (A)–(D) show the time courses of the nuclear STAT1 dimer *D_n_*(*t*), DNA-bound STAT1 *D_a_*(*t*), transcriptional memory *A_D_* (*t*), and SOCS1 negative feedback, respectively. Panels (E)–(H) report the corresponding response metrics for each output: peak amplitude *M*_1_, integrated response *M*_2_, and persistence *M*_5_. All metrics are expressed as relative values with respect to the *S* = 0 condition, corresponding to the absence of *p53*. Colours indicate the *p53* regimes: *S* = 0 without *p53*, *S* = 10 oscillatory, *S* = 17 damped, and *S* = 24 plateau-like. The grey vertical band indicates the duration of IFN activation.

Figure 4 shows that the nuclear phosphorylated STAT1 dimer pool *D_n_*(*t*) remains nearly invariant across the different *p53* regimes (panels A and E). Although the *p53*-dependent gates act on the nuclear STAT1 layer, they do not substantially expand the upstream pool of nuclear phosphorylated STAT1 dimers. The coupling therefore does not primarily operate by increasing the amount of available nuclear STAT1.

By contrast, the DNA-bound active STAT1 state *D_a_*(*t*) increases in amplitude and persistence as *p53* activity becomes stronger (panels B and F), consistent with the memory and synchronisation gates acting directly on DNA binding, unbinding, dephosphorylation, and nuclear import. The contrast between *D_n_*(*t*) and *D_a_*(*t*) is the key point: the same nuclear STAT1 availability is decoded into a stronger and more persistent transcriptionally active STAT1 state when *p53*-dependent coupling is present.

This redistribution is amplified at the level of the transcriptional memory variable *A_D_*(*t*) (panels C and G). The strongest changes are observed for the integrated response *M*_2_ and the post-peak persistence metric *M*_5_, showing that the main downstream consequence of the coupling is cumulative and temporal rather than instantaneous. The effect of *p53* is thus a reorganisation of STAT1 transcriptional decoding: post-phosphorylation coupling preferentially increases the integrated and persistent components of the transcriptionally active STAT1 response.

SOCS1 is also co-amplified alongside *D_a_* (panels D and H). This must be interpreted carefully, because SOCS1 can be regulated by *p53* in a context-dependent manner in biological systems (Będzińska et al., 2025; Sheikh and Sen, 2021). In the present model, however, no explicit *p53* →SOCS1 regulatory term is included. SOCS1 is kept as a STAT1-driven negative-feedback variable, with transcription driven by the DNA-bound STAT1 state according to

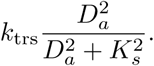

The SOCS1 increase observed here is therefore a model-intrinsic consequence of the larger and more persistent *D_a_*(*t*) signal: *p53* does not directly activate SOCS1 in the model; rather, the *p53*-dependent prolongation of DNA-bound STAT1 is sufficient to reinforce STAT1-driven negative feedback.

This signalling-level interpretation was preserved under broad variation of the *p53*-dependent memory and synchronisation gate parameters (Supplementary Figure S16). Across sampled parameter sets, plateau-like *p53* did not primarily increase the peak nuclear STAT1 dimer pool, but consistently increased DNA-bound STAT1 persistence, integrated DNA-bound STAT1 activity, and transcriptional memory.

In summary, *p53* coupling leaves the nuclear STAT1 dimer pool largely unchanged while redistributing signalling towards DNA-bound STAT1, transcriptional memory, and STAT1-driven negative feedback. This extends the window during which downstream STAT1-dependent programmes, including IRF1 induction and antiviral effector expression, can be engaged, while preserving an endogenous feedback structure that contributes to signal termination.

### 3.2. Impact of interferon timing on signal integration

Having shown that the *p53*-dependent gates redistribute nuclear STAT1 signalling towards DNA-bound activity and transcriptional memory, we next asked whether the timing of IFN stimulation modifies this effect. Some timing dependence is expected, since the memory variable *P_s_* accumulates with *p53* activity and relaxes over time; the relevant quantity is how strongly this dependence affects the amplitude, integration, time-to-peak, and post-peak persistence of *D_a_*(*t*).

We therefore varied the IFN onset time *t*_IFN_ relative to the *p53* stress state and quantified the resulting DNA-bound STAT1 response *D_a_*(*t*) using normalised peak, AUC, time-to-peak, and post-peak persistence metrics.

Figure 5 shows that the effect of IFN timing depends on the underlying *p53* regime. In the oscillatory regime, the peak and AUC of *D_a_*(*t*) remain close to the *t*_IFN_ = 0 reference, indicating that moderate *p53* oscillations have only a limited effect on the integrated STAT1 transcriptional output. By contrast, the damped and plateau-like regimes produce a stronger response, especially at the level of AUC and post-peak persistence. Thus, when IFN stimulation occurs after sufficient accumulation of *p53*-dependent memory, the same JAK/STAT1 module becomes more competent to sustain DNA-bound STAT1 activity.

The increase in time-to-peak with *t*_IFN_ should be interpreted with caution, because delaying IFN stimulation necessarily shifts the response in time. The most informative quantities are therefore the amplitude, AUC, and persistence metrics. These show that the main effect of prior *p53* stress is not simply to delay the STAT1 response, but to strengthen and prolong the DNA-bound STAT1 state once IFN-induced signalling reaches the nuclear layer.

Mechanistically, this follows from the accumulation of *P_s_*: through Φ_mem_, *P_s_* increases the effective DNA-binding rate 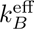, reduces the effective unbinding rate 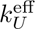, and stabilises the active nuclear STAT1 state by reducing 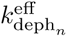 . The timing of IFN relative to *p53* determines whether this memory-dependent coupling is already available when the STAT1 signal enters the nucleus, thereby modifying the integrated and persistent components of the response.

Biologically, this provides a phenomenological representation of *p53*-dependent priming of STAT1-centred transcriptional activity, consistent with reported STAT1–*p53* functional crosstalk and with IFN-induced transcriptional memory at STAT1-regulated loci (Townsend et al., 2004; Youlyouz-Marfak et al., 2008; Kamada et al., 2018; Tehrani et al., 2023). The effect of *p53* is therefore not fixed by its amplitude alone, but also by its timing relative to IFN stimulation: a prior *p53* stress state can precondition the JAK/STAT1 nuclear layer, making the subsequent IFN response more integrated and persistent.

**Figure 5:**
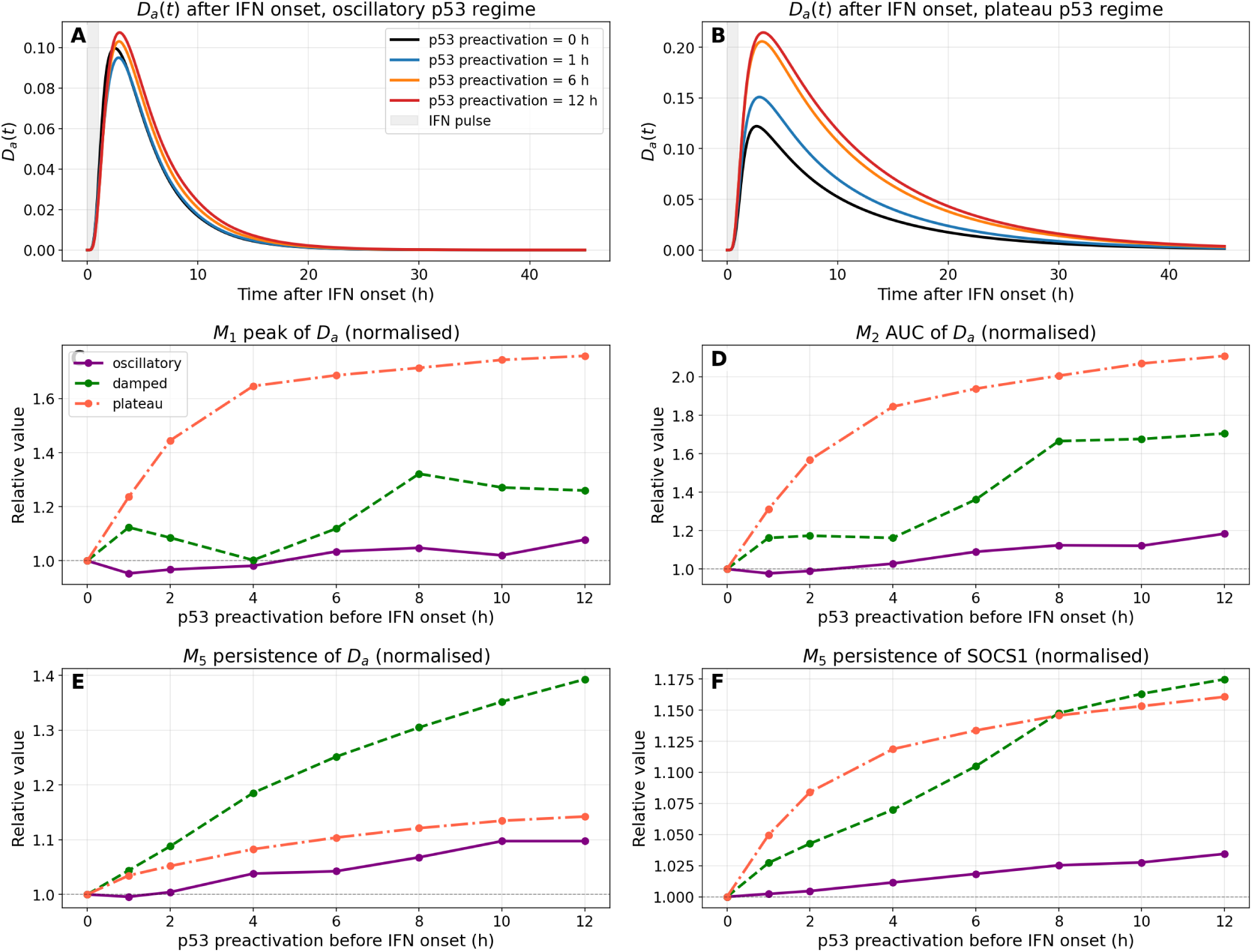
Effect of *p53* preactivation duration on the DNA-bound STAT1 response following a short IFN=50 pulse. The IFN stimulus is applied for 1 h, from IFN onset to 1 h after onset, and all time-course panels are aligned so that *t* = 0 corresponds to IFN onset. Panels A–B show *D_a_*(*t*) trajectories after IFN stimulation for selected *p53* preactivation times under oscillatory and plateau-like *p53* regimes, respectively; the grey shaded region indicates the 1 h IFN activation window. Panels C–F quantify how increasing *p53* preactivation before IFN onset modifies the STAT1 response: peak amplitude *M*_1_, area under the curve *M*_2_, *D_a_*(*t*) persistence *M*_5_, and SOCS1 persistence *M*_5_. All metric values are normalised to the no-preactivation condition, corresponding to IFN onset at *t*_IFN_ = 0. Colours indicate the *p53* regimes: oscillatory, damped, and plateau-like.

### 3.3. Memory and synchronisation have complementary effects

We next decomposed the coupling architecture to identify which components are responsible for these effects. This analysis is explicitly architectural: the memory-only and synchronisation-only scenarios were constructed to separate the contributions of the two implemented gates. The aim is therefore to determine whether the enhanced response arises mainly from acute co-activation, from accumulated memory, or from their combination.

We compared the core JAK/STAT1 response with memory-only, synchronisation-only, and full-coupling scenarios under the plateau-like *p53* regime, where the separation between the mechanisms is most clearly visible.

Figure 6 shows that the two coupling mechanisms have complementary effects on *D_a_*(*t*). The synchronisation component mainly enhances the early DNA-bound STAT1 response, increasing the initial amplitude of *D_a_*(*t*). By contrast, the memory component has a stronger effect on signal integration and post-peak persistence, indicating that it sustains the transcriptionally active STAT1 state after the initial response.

**Figure 6:**
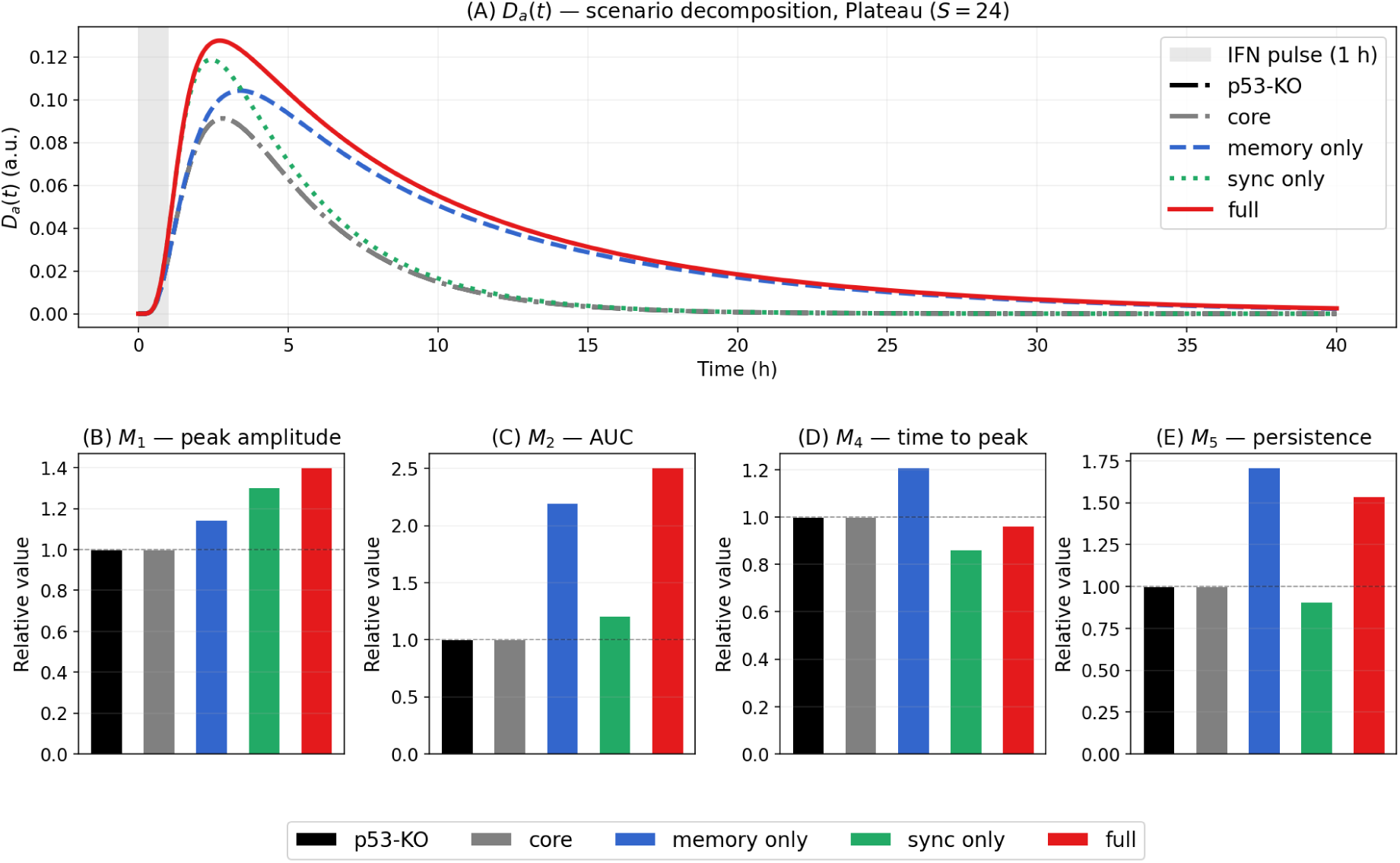
Scenario decomposition of the DNA-bound STAT1 response following transient IFN stimulation under the plateau-like *p53* regime (*S* = 24). IFN was applied for 1 h with amplitude 50. Panel (A) shows the time course of *D_a_*(*t*) for the *p53*-KO, core, memory-only, synchronisation-only, and full-coupling scenarios. Panels (B)–(E) report the corresponding normalised metrics: peak amplitude *M*_1_, AUC *M*_2_, time to peak *M*_4_, and post-peak persistence *M*_5_. All metrics are normalised to the *p53*-KO condition, indicated by the horizontal dashed line.

This separation is consistent with the definitions of the two gates: Φ_sync_ is an acute co-activation term depending on concurrent STAT1 and *p53* activity, whereas Φ_mem_ depends on the accumulated memory variable *P_s_*. The informative result is that the full response cannot be reproduced by one mechanism alone. Synchronisation provides early amplification, memory provides persistence, and the full-coupling scenario combines both behaviours. Thus, the *p53*-dependent regulation of JAK/STAT1 is not controlled by a single effective gain, but by the combination of an acute amplifier and a slower persistence module.

### 3.4. Prior stress delays STAT1 recovery

We next asked whether the *p53*-dependent persistence of nuclear STAT1 activity affects the capacity of the JAK/STAT1 pathway to respond to repeated IFN stimulation. The previous analyses showed that the implemented memory gate can prolong DNA-bound STAT1 activity and transcriptional memory. Here, we asked whether this persistence also delays the return of the pathway to a fully re-inducible state. Unlike the direct increase in DNA-bound persistence, this recovery behaviour is not imposed by a single gate; it emerges from the interaction between memory accumulation, pathway reset, and repeated IFN stimulation.

To quantify re-inducibility, we used the nuclear STAT1 dimer pool *D_n_*(*t*) as an upstream readout of pathway restart. In contrast to *D_a_*(*t*), which reflects DNA binding, residence time, and transcriptional persistence, *D_n_*(*t*) reports whether a new pool of phosphorylated STAT1 dimers can be generated and transported to the nucleus after a previous IFN pulse. A complementary analysis using *D_a_*(*t*), together with a return-to-baseline quantification of transcriptional persistence, is provided in Supplementary Section S5 (Figure S3 and Figure S4).

**Figure 7:**
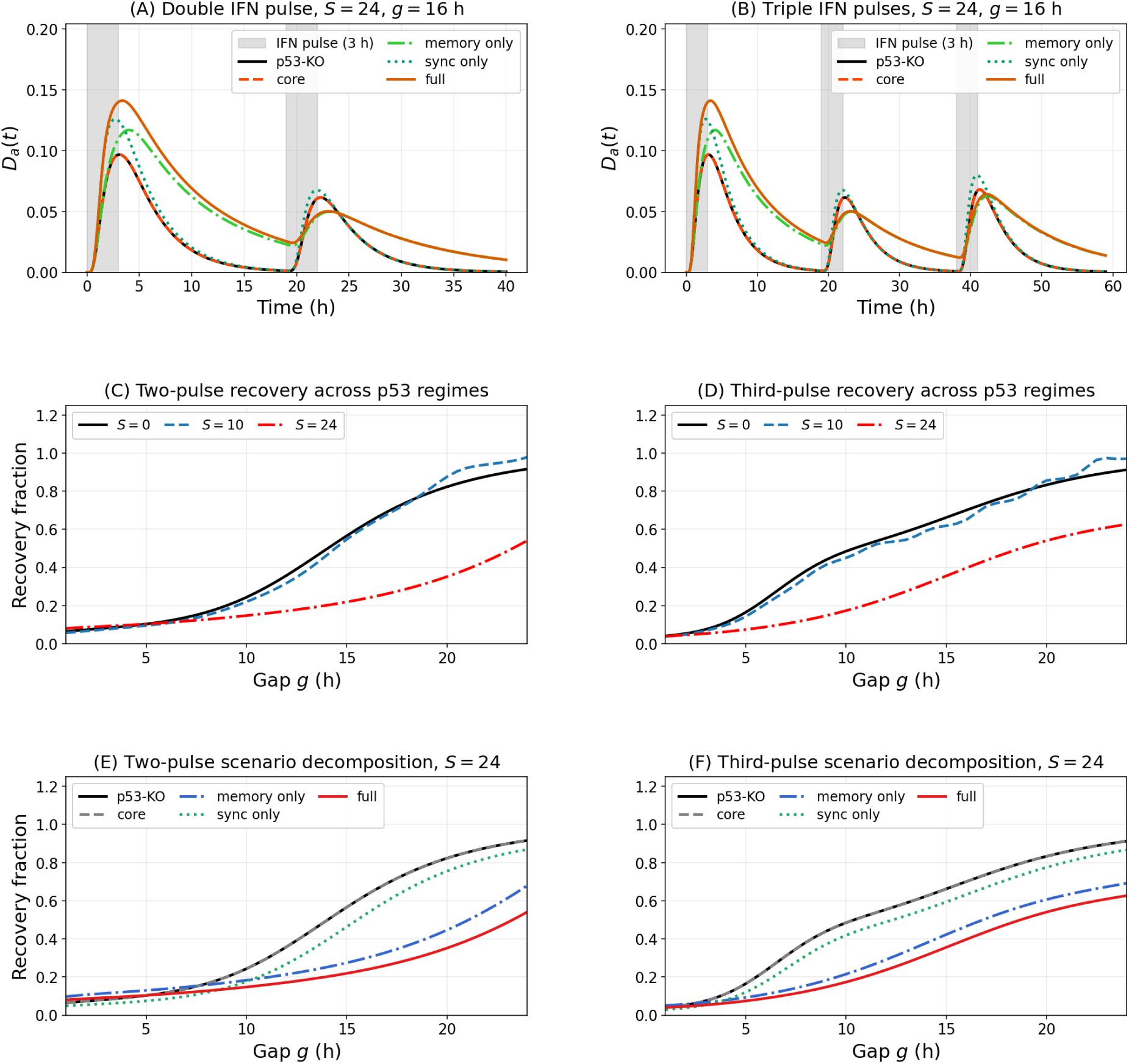
Recovery analysis of the DNA-bound STAT1 response *D_a_*(*t*) under repeated IFN stimulation. Each IFN pulse has the same amplitude, *IFN* = 50, and the same duration, 3 h. Panels (A)–(B) illustrate the time-course protocol at *S* = 24 for the scenario decomposition: a double-pulse protocol and a triple-pulse protocol, respectively, using an equal interpulse gap *g* = 16 h. The grey shaded regions denote the 3 h IFN activation windows. For the double-pulse protocol, the second pulse is applied after a gap *g* following the end of the first pulse; for the triple-pulse protocol, the same gap *g* separates the first and second pulses and the second and third pulses. Panels (C)–(D) quantify the two-pulse and third-pulse recovery fractions across *p53* regimes in the full model, for *S* = 0, *S* = 10, and *S* = 24. Panels (E)–(F) show the corresponding scenario decomposition at *S* = 24, comparing the *p53*-KO, core, memory-only, synchronisation-only, and full-coupling scenarios. Recovery fractions are computed as the peak *D_a_* response induced by the tested pulse, after subtraction of the matched no-tested-pulse control, normalised to the peak *D_a_* response elicited by the first pulse. Thus, R*_Da_* (*g*) measures second-pulse recovery, whereas R_3,*D*_*_a_* (*g*) measures third-pulse recovery as functions of the interpulse gap *g*.

For the *k*-th pulse, with *k* ∈ {2, 3}, and for an interpulse gap *g*, the incremental recovery fraction was defined as

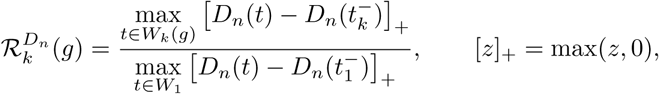

where *W*_1_ is the response window after the first IFN pulse, *W_k_*(*g*) is the response window after the *k*-th pulse, and 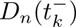 is the level of nuclear STAT1 dimers immediately before the *k*-th stimulation. Values close to one indicate full recovery, whereas lower values indicate refractoriness.

Figure 7 shows that *D_n_*(*t*) remains transient after a single IFN pulse across the *p53* regimes, indicating that the pathway is not locked in a permanently activated nuclear STAT1 state. However, repeated stimulation reveals a clear *p53*-dependent refractory effect. At short interpulse gaps, the second and third pulses generate only a fraction of the initial *D_n_* response, and recovery progressively improves as the gap lengthens. The hierarchy between regimes is consistent: the no-*p53* condition recovers fastest, the oscillatory regime is intermediate, and the plateau-like regime remains refractory for the longest time.

The decomposition of the *S* = 24 condition identifies the memory component as the main contributor to this delayed recovery. The memory-only and full-coupling scenarios show the strongest reduction in second- and third-pulse recovery, whereas the core and synchronisation-only scenarios recover more efficiently. This result is consistent with the construction of Φ_mem_, but it is not a simple restatement of the memory gate: the observed refractoriness depends on how memory-dependent persistence affects pathway reset and re-induction after repeated stimulation.

This behaviour is compatible with experimental evidence that interferon signalling can exhibit long-lasting desensitisation upon repeated stimulation (Kalliara et al., 2022; Sarasin-Filipowicz et al., 2009), and with reports that interferon stimulation can establish transcriptional memory (Kamada et al., 2018; Tehrani et al., 2023). It is also consistent with evidence that *p53*-activating stress can modulate STAT1 signalling in a context-dependent manner (Youlyouz-Marfak et al., 2008; Townsend et al., 2004; Będzińska et al., 2025). However, these studies do not directly demonstrate that *p53*-activating stress delays the recovery of nuclear STAT1 dimers after repeated IFN stimulation. This delayed recovery is therefore a specific prediction of the present model. The delayed-recovery hierarchy was also robust to gate-parameter uncertainty (Supplementary Figure S17). Across sampled parameter sets, the recovery deficit remained lowest in the no-*p53* condition, intermediate under oscillatory *p53*, and highest under plateau-like *p53*. Thus, the persistence–recovery trade-off is not restricted to the nominal gate-parameter calibration. Together, these simulations suggest that the temporal order of stress and IFN exposure can determine whether JAK/STAT1 signalling remains readily re-inducible. When *p53*-dependent memory is already established before IFN restimulation, the pathway becomes less able to regenerate a full nuclear STAT1 dimer response at short interpulse gaps. By contrast, when IFN stimulation occurs before the memory layer has fully accumulated, the early response is better preserved. Thus, prior *p53*-activating stress is predicted to promote a more refractory STAT1 state, whereas early or concurrent IFN exposure can preserve stronger initial re-induction before memory-dependent refractoriness emerges.

## 4. Prescribed *p53* and IFN inputs reveal how *p53* modulates antiviral response

Having characterised how the *p53*-dependent coupling layer modifies post-phosphorylation STAT1 decoding, we next examine how this altered decoding is transmitted to downstream antiviral effectors, viral control and apoptotic routing. The aim is not to re-establish the signalling-level memory and persistence mechanisms, which are part of the coupling architecture, but to determine how they shape functional outcomes once the response is propagated to the effector and viral-burden layers.

We first analysed a controlled prescribed-input setting in which both the IFN stimulus and the *p53* regime were imposed externally. This provides a host-side baseline: the IFN source is fixed, the *p53* dynamics are prescribed, and the downstream consequences of *p53*-dependent JAK/STAT1 modulation can be examined without feedback from virus-induced IFN production or infection-induced checkpoint stress.

The IFN stimulus was applied for 120 h and then withdrawn exponentially. Three upstream conditions were compared: the JAK/STAT1 core condition, an oscillatory *p53* profile, and a plateau-like *p53* profile. The analysis was organised into transcriptional-memory outputs (*A_D_, M*_IRF1_*, A*_IRF1_), antiviral effectors (*P*_1_*, P*_2_*, P*_3_), and viral/apoptotic outputs (*V_g_, V_p_, P*_4_*, P*_5_). Antiviral-state engagement, viral outcome and apoptotic commitment were classified using the common readout framework defined above, with *L*_AV_ reported as a normalised score taking the values 0, 1*/*3, 2*/*3, and 1.

### 4.1. Prescribed p53 amplifies the memory and effector layers

Figure 8 examines how prescribed *p53* alters the IFN-induced memory–effector architecture. Here, *A_D_* denotes accumulated DNA-bound STAT1 activity, *M*_IRF1_ a slower IRF1-memory state, and *A*_IRF1_ a delayed IRF1-associated activation layer. Because *p*53-dependent gates can amplify memory and effector transmission, the relevant question is how this coupling is distributed across the memory and effector layers.

**Figure 8:**
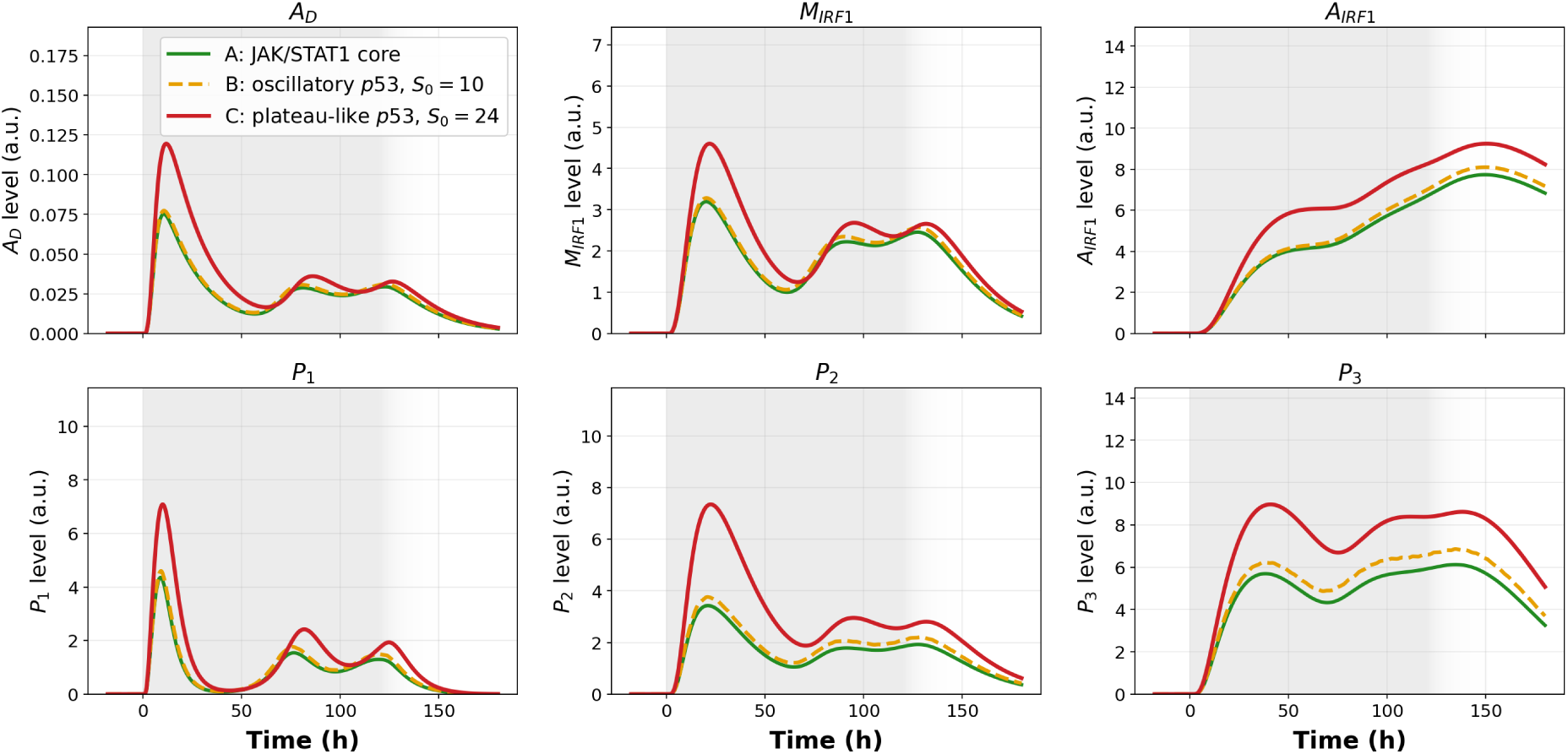
Effect of prescribed *p53* regimes on the memory and effector layers under a fixed prolonged external IFN stimulus (IFN = 20 a.u. from 0 to 120 h). The upper row shows the transcriptional and memory variables *A_D_*, *M*_IRF1_, and *A*_IRF1_. The lower row shows the antiviral effectors *P*_1_, *P*_2_, and *P*_3_. Curves compare the JAK/STAT1 core condition, oscillatory *p53*, and plateau-like *p53*.

The JAK/STAT1 core condition already initiates the antiviral programme under the prescribed IFN stimulus, and prescribed *p53* increases the intensity with which this programme is transmitted through the memory and effector layers. Crucially, this amplification is not uniform. The plateau-like *p53* regime produces the strongest increase in *A_D_*, *M*_IRF1_, and *A*_IRF1_, indicating that sustained *p53* activity preferentially reinforces the integrated, memory-associated layer. At the effector level, the increase is modest for *P*_1_, which is closer to the proximal STAT1 response, whereas it becomes more pronounced for *P*_2_ and especially *P*_3_, which depend on more integrated memory and IRF1-associated signals.

The imposed coupling is thus filtered through the network: the largest effects appear in variables that integrate signalling over longer timescales. The oscillatory regime is intermediate, and the plateau-like regime gives the largest downstream output. This establishes the first downstream consequence of the coupling: sustained *p53* preferentially strengthens the memory-dependent and IRF1-associated components of the antiviral programme.

### 4.2. p53 preactivation is buffered at the effector level

We then asked whether activating *p53* before IFN onset further modifies the downstream memory and effector layers. Since the *p53*-dependent memory gate can be partly engaged before IFN stimulation, some effect at the STAT1 level is expected; the question is whether this upstream timing effect is transmitted proportionally to *A_D_*, *M*_IRF1_, *A*_IRF1_, and the antiviral effectors.

**Figure 9:**
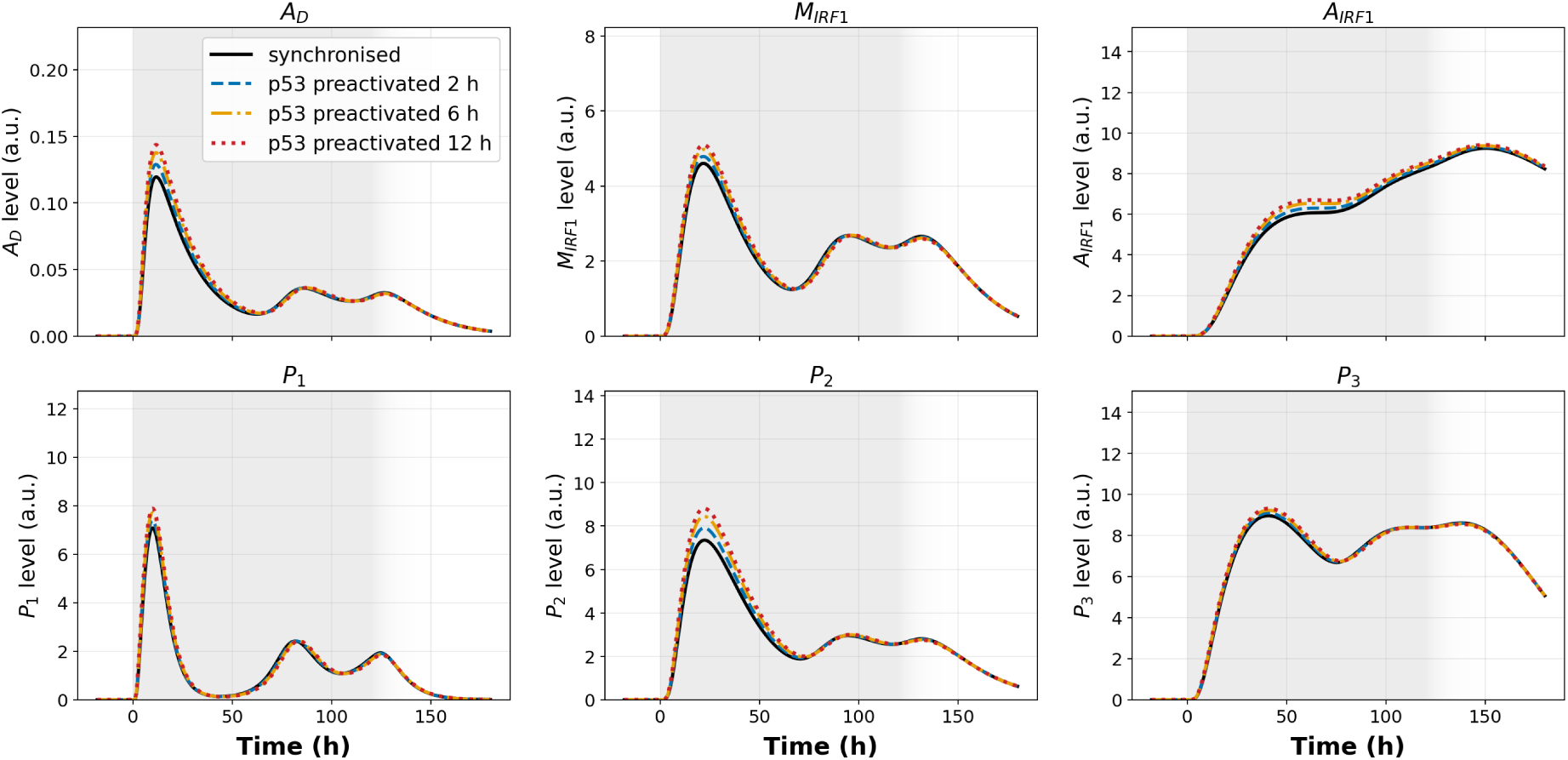
Effect of *p53* preactivation on the memory and effector layers under the same prolonged external IFN stimulus (IFN = 20 a.u. from 0 to 120 h). The synchronised condition is compared with *p53* preactivation 2 h, 6 h, and 12 h before IFN onset. The upper row shows *A_D_*, *M*_IRF1_, and *A*_IRF1_, and the lower row shows *P*_1_, *P*_2_, and *P*_3_.

Figure 9 shows that the downstream memory–effector layer is only weakly affected by the duration of *p53* preactivation. A small increase is visible when *p53* is activated before IFN onset, especially for *A_D_*, *M*_IRF1_, *P*_1_, and *P*_2_, but the trajectories obtained with 2 h, 6 h, and 12 h of preactivation remain close to one another. Thus, although preactivation can increase the upstream STAT1 response, this gain is not transmitted linearly to the downstream effector programme.

This buffering is not imposed by a single *p53* gate, but emerges as the signal passes through IRF1 accumulation, memory integration, effector production and effector turnover. These downstream layers smooth differences generated at the STAT1 level, so that larger changes in DNA-bound STAT1 persistence produce only modest changes in *P*_1_, *P*_2_, and *P*_3_. The effect is saturation-like: once *p53*- dependent modulation is present at IFN onset, extending the preactivation window provides little additional effector gain.

This behaviour was preserved under broad gate-parameter variation (Supplementary Figure S18). Although DNA-bound STAT1 remained sensitive to the duration of *p53* preactivation, the antiviral effectors showed a smaller relative spread across preactivation delays. Thus, *p53* timing is more visible at the signalling layer than at the final effector layer: prior *p53* activity can prime STAT1 decoding, but the memory–effector module partially buffers this priming before it reaches the antiviral outputs.

### 4.3. External IFN duration gates delayed P*_4_*-associated apoptosis

We finally asked whether the *p53*-dependent amplification of the memory–effector layer is sufficient to improve viral control and to route apoptosis under an externally prescribed IFN input. In this controlled setting, IFN is imposed independently of viral burden: the external IFN pulse is fixed from 0 to 120 h, and therefore does not rise or decay as a function of *V_g_*. This makes the prescribed-IFN setting useful as a duration test: it separates the effect of sustained JAK/STAT–IRF1 drive from the infection-coupled IFN feedback analysed later.

Figure 10 shows that increasing or prolonging the externally imposed IFN response does not make antiviral control and apoptotic routing equivalent. In the sensitive viral class, the viral burden is rapidly reduced and the system reaches a cleared outcome. Because *V_g_* does not persist, the *P*_4_-associated route remains inactive even when the external IFN amplitude is high. Under plateau-like *p53*, the model can still trigger early *P*_5_-associated commitment, reflecting the prescribed *p53*-dependent apoptotic arm rather than a *P*_4_-mediated danger route.

**Figure 10:**
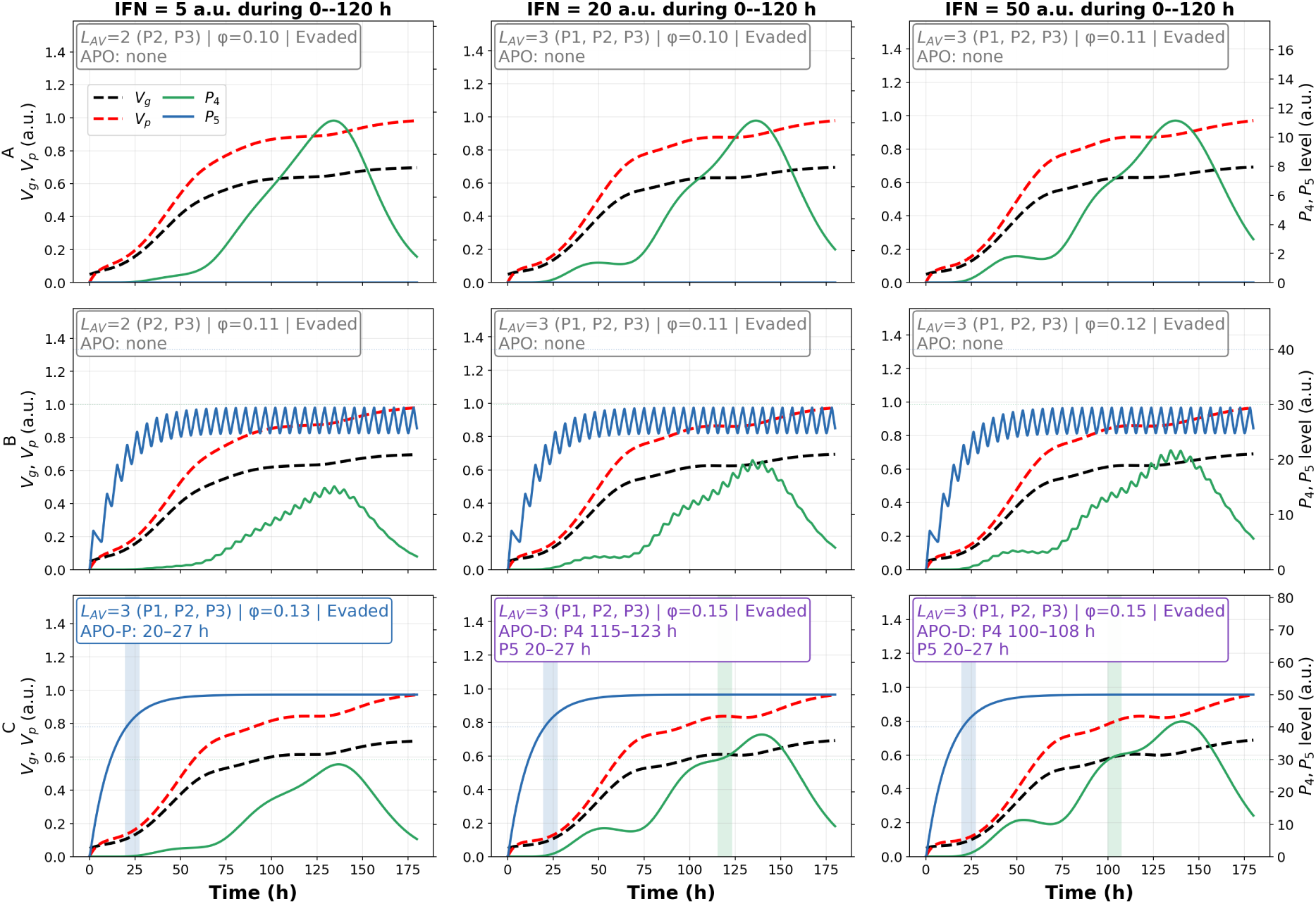
Class-specific prescribed-IFN simulations showing antiviral-state engagement, viral outcome, and apoptotic commitment under prolonged external IFN input. Rows are labelled by upstream regime: A, JAK/STAT1 core; B, oscillatory *p53* (*S*_0_ = 10); and C, plateau-like *p53* (*S*_0_ = 24). Columns correspond to external IFN amplitudes applied from 0 to 120 h, followed by exponential withdrawal. Black and red dashed curves denote *V_g_* and *V_p_*, respectively; green and blue curves denote *P*_4_ and *P*_5_. The curve legend is shown in the first panel only. Decision boxes report *L*_AV_, the viral-control score *φ*, the realised viral outcome, and the selected apoptotic route. Box colours encode apoptotic routing: grey, no apoptosis; green, APO-J; blue, APO-P; and violet, APO-D.

The intermediate viral class shows a different behaviour. The host response is strongly engaged, and increasing the external IFN amplitude improves the viral-control score *φ*, especially under plateau-like *p53*. However, this improvement remains distinct from *P*_4_-associated apoptotic commitment. Even with a prolonged external IFN pulse, *P*_4_ does not automatically cross its sustained apoptotic threshold in the intermediate class. Thus, in this prescribed-input setting, high IFN-driven antiviral activation is not by itself sufficient to route the system through *P*_4_.

The highly resistant viral class reveals the condition under which delayed *P*_4_-associated apoptosis can appear. Viral burden persists despite strong antiviral-state engagement, keeping the viral-burden gate Φ*_V_* _4_(*V_g_*) available. In the plateau-like *p53* regime, extending the external IFN pulse to 120 h allows the JAK/STAT–IRF1-associated *P*_4_ output to accumulate long enough to satisfy the sustained-threshold rule at higher IFN amplitudes. The resulting APO-D cases combine early *P*_5_ commitment, driven by prescribed sustained *p53*, with delayed *P*_4_ commitment, driven by prolonged IFN/JAK–STAT–IRF1 signalling in the presence of persistent viral burden.

Because external IFN is decoupled from viral burden, its amplitude alone does not guarantee *P*_4_-apoptosis: sustained JAK/STAT–IRF1 signalling must coincide with persistent *V_g_*, through the Φ*_V_* _4_(*V_g_*) gate, and with sufficient *p53*-dependent apoptotic amplification. *P*_4_ therefore behaves as a duration-sensitive, context-dependent danger route, whereas *P*_5_ reports the imposed *p53*-autonomous arm. More broadly, this prescribed-input setting fixes the baseline propagation logic of the model: prescribed *p53* strengthens the memory and effector layers, but its functional impact depends on viral sensitivity, so that a high *L*_AV_ can coexist with poor viral control in resistant classes. The next section relaxes the external drive and lets IFN production and *p53* activation arise dynamically from viral burden.

## 5. Antiviral response under dual IFN-source control

The prescribed-input analysis showed that high levels of *p53* can amplify the transmission of IFN-induced STAT1 activity to IRF1-associated memory and antiviral effectors. We next asked how this amplification is modified when the IFN signal contains two components: an externally supplied IFN input and a virus-induced IFN input generated by viral sensing. Here, *p53* remains prescribed, so the analysis isolates the role of IFN-source composition from virus-induced *p53* activation.

The two IFN components differ in their relation to infection dynamics. The external component is imposed and can activate JAK/STAT1 independently of the current viral burden, whereas the virus-induced component remains coupled to *V_g_*. As a result, virus-induced IFN carries temporal information about viral persistence, while external IFN mainly provides a prescribed upstream drive. The key question is therefore not whether persistent viral burden can maintain virus-induced IFN, which follows directly from the model structure, but how this source-dependent timing reshapes the downstream readouts. We specifically ask whether the virus-coupled IFN component changes the relationship between normalised antiviral-state engagement *L*_AV_, functional viral control *φ*, and route-specific apoptotic commitment. This distinction is important because sustained antiviral activation does not necessarily imply viral clearance, especially for poorly sensitive viral classes.

The two apoptotic routes also decode the IFN–*p53* network differently. The *P*_4_ route requires sustained IFN–IRF1 signalling in a persistent viral-burden context, through the Φ*_V_* _4_(*V_g_*) gate, whereas the *P*_5_ route is mainly controlled by the imposed *p53* regime in this prescribed-*p53* setting.

### 5.1. A virus-coupled IFN component sustains the IRF1–P*_4_* danger route under persistent viral burden

We next combined an externally supplied IFN component with a virus-induced one, asking how the virus-coupled component changes the downstream interpretation of IFN signalling when viral burden is unresolved. Because the external component is imposed by protocol while the virus-induced component stays coupled to *V_g_*, the two carry different temporal information, and this difference—rather than total IFN amplitude—is expected to shape viral control and apoptotic routing (Schneider et al., 2014; Muñoz-Fontela et al., 2008; Rivas et al., 2010).

Figure S11, in the supplementary, resolves this at the trajectory level and Figure 11 generalises it across the virus-induced IFN capacity IFN_max_; both show that the same dual-source architecture is read differently depending on viral class. Where the class remains effector-sensitive—sensitive and intermediate classes—increasing IFN_max_ raises antiviral engagement and translates into higher *φ*, so that *L*_AV_ and *φ* rise together. The highly resistant class shows the opposite pattern: *L*_AV_ stays high while *φ* remains low. This dissociation is not imposed by the IFN-source structure but emerges from strong host activation combined with weak viral sensitivity: the virus-coupled component maintains the danger signal, yet the activated effectors still fail to suppress genome replication or particle production.

The apoptotic layer separates an expected component from a contingent one. The *P*_5_ route tracks the prescribed *p53* regime, so early *P*_5_-associated commitment under plateau-like *p53* is expected and generates a broad APO-P region. The *P*_4_ route is contingent: although the persistence of the virus-coupled IFN component is structural, route selection is not, since it depends on whether the maintained IFN–IRF1 signal keeps *P*_4_ above threshold long enough through Φ*_V_* _4_(*V_g_*), and on whether *P*_5_ has already been recruited. This is why, in the highly resistant conditions, unresolved viral burden keeps the IFN–IRF1 danger signal active and lets *P*_4_ accumulate with delay: under oscillatory *p53* the persistent signal appears mainly as APO-J, whereas under plateau-like *p53* the early *P*_5_ pressure combines with delayed *P*_4_ recruitment to give dual-route APO-D.

The informative outcome is therefore not that virus-coupled IFN persists—which is structural—but that this persistence can change the apoptotic route selected by a system in which *p53* is held fixed. IFN-source composition is thus not reducible to a total-amplitude effect: whether the response ends in viral control, *P*_4_-, *P*_5_- or dual-route apoptosis depends jointly on viral effector sensitivity and on the prescribed *p53* regime.

**Figure 11:**
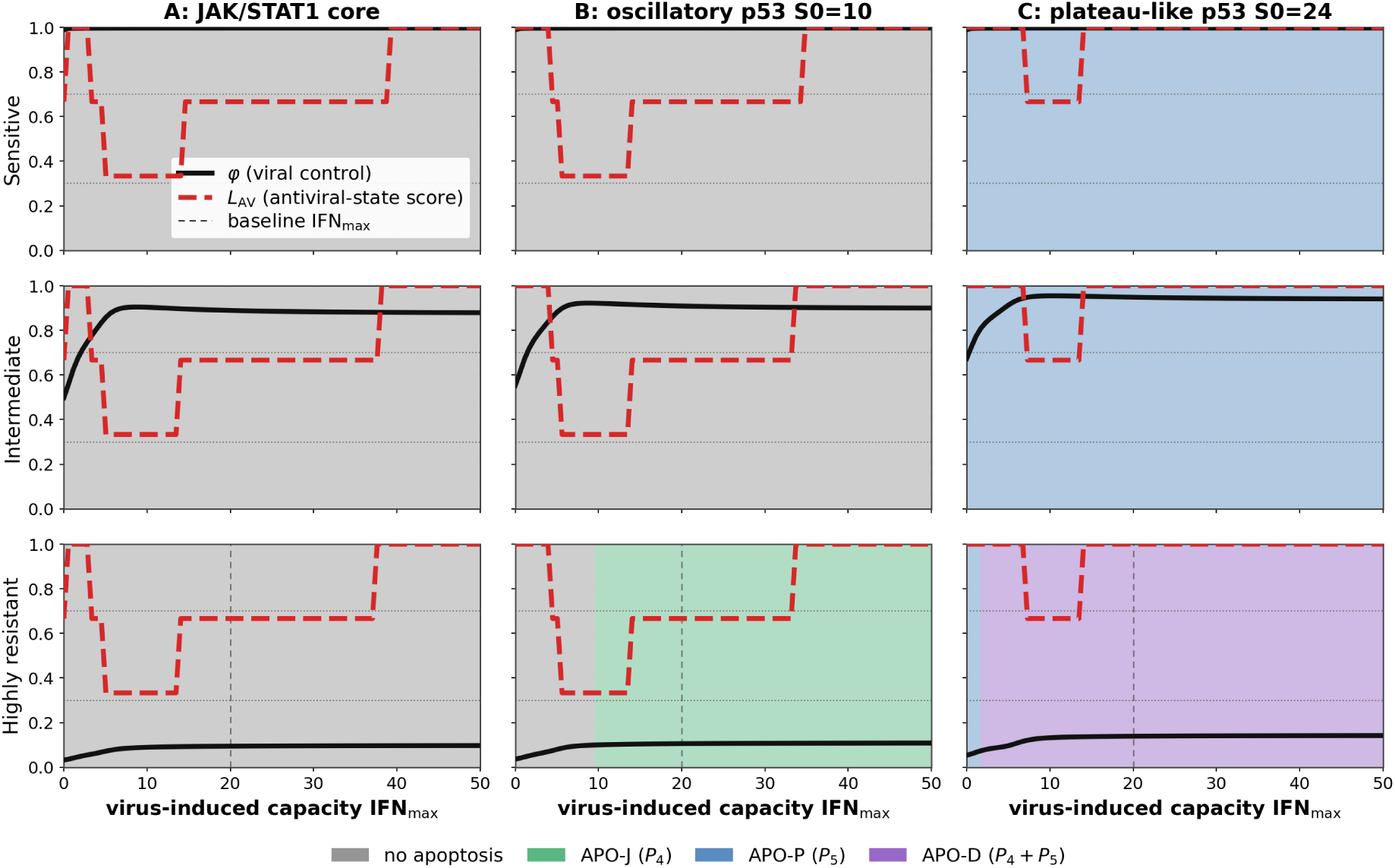
Response-map summary of antiviral control and apoptotic routing under dual IFN input. Rows correspond to viral effector-sensitivity classes: sensitive, intermediate, and highly resistant. Columns correspond to prescribed *p53* regimes: JAK/STAT1 core, oscillatory *p53* (*S*_0_ = 10), and plateau-like *p53* (*S*_0_ = 24). In each panel, the horizontal axis gives the virus-induced IFN capacity IFN_max_. Background colours indicate the apoptotic route selected by the sustained-threshold rule: no apoptosis, APO-J for *P*_4_-associated commitment, APO-P for *P*_5_-associated commitment, and APO-D for dual *P*_4_ + *P*_5_ commitment. The same apoptotic-route colour code is used as in the response-map and time-course decision figures. The solid black curve shows the viral-control score *φ*, whereas the dashed red curve shows the antiviral-state score *L*_AV_, with discrete levels 0, 1*/*3, 2*/*3, and 1. Horizontal dotted lines mark reference readout levels used to interpret weak, intermediate, and strong responses. Vertical dashed lines indicate the baseline class-specific IFN_max_ values used in the corresponding time-course simulations.

### 5.2. Constant-sum IFN-source composition separates imposed IFN exposure from virus-coupled IFN feedback

To test whether the relative contribution of the two IFN sources matters beyond the configured maximal IFN budget, we performed a constant-sum IFN-source composition scan. The external and virus-induced components were constrained to share the same maximal IFN budget, set to the saturation scale of the model:

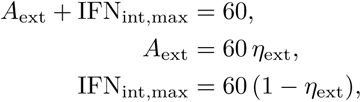

with 0 ≤ *η*_ext_ ≤ 1. Thus, *η*_ext_ = 0 corresponds to a purely virus-induced IFN configuration, whereas *η*_ext_ = 1 corresponds to a purely externally imposed IFN configuration. The corresponding virus-induced production rate is

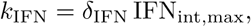

with *δ*_IFN_ = 0.12. This scan is therefore a redistribution of the same maximal IFN budget between an imposed external source and a virus-coupled endogenous source, not a change in total IFN capacity.

Because the external component is imposed and withdrawn while the virus-induced component stays coupled to viral burden, late IFN necessarily declines as *η*_ext_ increases. The question is not whether the virus-coupled component maintains IFN longer—it does, by construction—but how replacing virus-coupled feedback by imposed exposure changes late viral burden and apoptotic-route selection.

**Figure 12:**
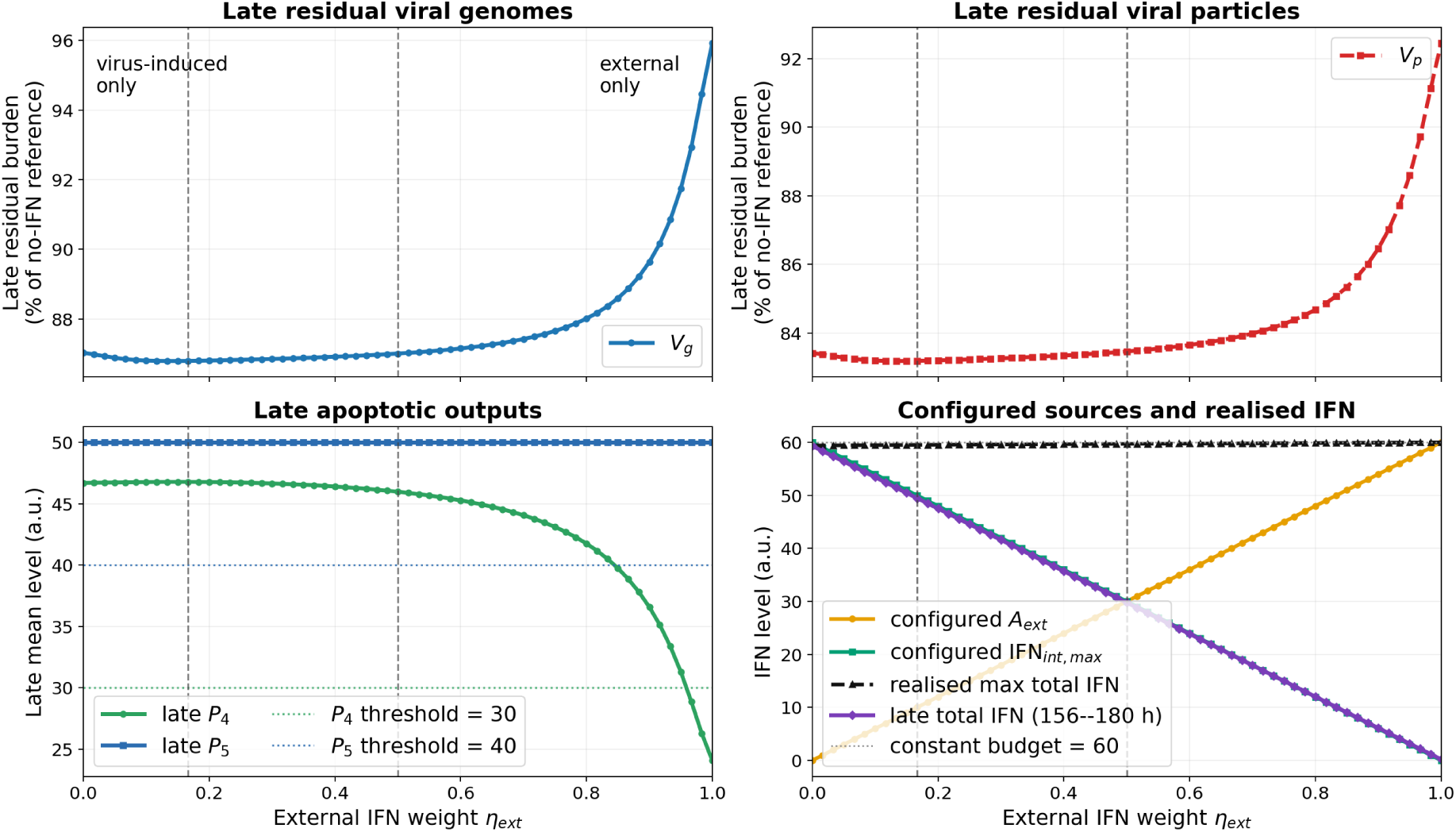
Constant-sum IFN-source composition under plateau-like prescribed *p53* in the highly resistant viral class. The external IFN weight *η*_ext_ varies from 0 to 1, with *A*_ext_ = 60*η*_ext_ and IFN_int,max_ = 60(1 − *η*_ext_), so that *A*_ext_ + IFN_int,max_ = 60 for all source compositions. The virus-induced production rate is *k*_IFN_ = *δ*_IFN_IFN_int,max_, with *δ*_IFN_ = 0.12. The upper panels show the late residual viral genome and particle burdens, expressed as percentages of the no-IFN reference. The lower-left panel shows the late apoptotic outputs *P*_4_ and *P*_5_. The lower-right panel shows the configured IFN-source composition together with the realised maximal and late total IFN levels. Dashed vertical markers indicate the minimal external reference *A*_ext_ = 10 and the equal-composition point *η*_ext_ = 0.5. The late window is 96–120 h.

Figure 12 shows the downstream consequences of this redistribution. Because the maximal IFN budget is fixed, the differences across the scan cannot be attributed to a larger configured IFN capacity; they reflect the replacement of a virus-coupled feedback source by an externally imposed one. As *η*_ext_ increases, late total IFN decreases, since a larger fraction of the budget is assigned to the withdrawn external component.

The informative result is how this late IFN loss affects the viral and apoptotic readouts. In the highly resistant class, residual *V_g_* and *V_p_* stay high across the scan, confirming poor control even at a large configured IFN budget. However, residual burden is lowest when the virus-induced component dominates and rises towards the external-only limit. For a resistant viral class, maintaining a feedback link between viral burden and IFN production is therefore more effective than replacing the same maximal budget with an imposed pulse that becomes disconnected from late infection.

The apoptotic outputs separate the two source-dependent mechanisms. The *P*_5_ route stays high and nearly invariant across the scan, consistent with its dependence on the prescribed plateau-like *p53* input. In contrast, *P*_4_ decreases as *η*_ext_ increases, because *P*_4_ requires sustained IFN–IRF1 signalling together with the viral-burden-gated danger route: when the IFN budget is mainly virus-induced, persistent viral burden maintains the upstream danger signal and supports higher late *P*_4_; as the budget is transferred to the external source, the late IFN signal weakens after withdrawal and the drive to *P*_4_ is reduced.

The constant-sum scan thus clarifies the role of IFN-source composition. The two sources differ in temporal coupling—one imposed by protocol, the other linked to viral burden—and this difference, at fixed total budget, changes late residual viral burden and selectively affects the *P*_4_ route while leaving the *P*_5_ route largely governed by prescribed *p53*. Source composition therefore cannot be reduced to a single total IFN amplitude.

## 6. Virus-induced IFN and virus-induced *p53* couple antiviral control to apoptotic routing

We next considered a fully infection-coupled configuration in which viral burden induces both IFN production and *p53*-activating stress. In this setting, *V_g_* is no longer only a downstream target of the antiviral programme; it also acts as an upstream input that sustains endogenous IFN production, checkpoint-stress activation, and the viral-burden gate required for *P*_4_-associated commitment. These dependencies are structural. The informative question is how they translate into downstream outcomes once viral effector sensitivity, viral persistence, IFN-induction capacity, *p53*-stress induction, and route-specific thresholds are combined.

This configuration provides a direct infected-cell counterpart to the previous controlled settings. The IFN branch represents viral sensing followed by JAK/STAT1 activation, whereas the *p53* branch represents infection-associated stress, which may arise from replication stress, oxidative or metabolic perturbation, damage-like signalling, or other disturbances associated with persistent infection. The analysis asks how viral classes shape the duration and intensity of infection-coupled IFN and *p53* signals, and how this coupling affects antiviral-state activation, functional viral control, and apoptotic-route selection.

Analyses in which viral-to-stress conversion is combined with both external and virus-induced IFN sources are provided in the Supplementary Material, Section S7.4. The calibration curves for viral-to-IFN and viral-to-*p53* stress conversion are shown in Supplementary Figure S5 and Figure S6.

### 6.1. Viral persistence converts p53 into a burden-dependent amplifier of the memory–effector layer

We first examined how the fully infection-coupled setting shapes the memory–effector layer. Since *V_g_* drives both endogenous IFN production and *p53*-associated stress, some late reinforcement of the response is expected when viral burden persists; the informative question is how this maintained input is *distributed* across the memory and effector layers, and how this distribution depends on viral class.

Figure 13 illustrates the highly resistant, strongly IFN-inducing configuration. The early response is driven mainly by virus-induced IFN and JAK/STAT1 activation, so the separation between *p53* regimes is initially limited. The later separation reflects the infection-coupled structure: persistent viral burden maintains the *p53*-associated stress input and lets the *p53*-dependent gates act over a longer time window.

This reinforcement is not uniform across the network. *A_D_* and *P*_1_, which are closer to the proximal STAT1-dependent response, retain a stronger early component and discriminate less between weak *p53* conditions. By contrast, *M*_IRF1_, *A*_IRF1_, *P*_2_, and especially *P*_3_ show larger late differences between regimes: these variables integrate signalling over longer timescales and therefore retain more information about persistent viral burden and accumulated *p53*-dependent modulation.

The magnitude of this late separation depends on viral resistance. In an intermediate class, viral burden persists long enough to reveal a moderate *p53*-dependent reinforcement; in a highly resistant class, viral burden remains elevated for longer and continuously supplies both IFN-inducing and *p53*-activating inputs, producing the clearest separation between the core, sustained, damped, and plateau-like conditions. Viral resistance therefore has a dual consequence: it reduces the functional efficiency of the antiviral effectors, but it prolongs the upstream signals that drive their production.

Viral IFN-induction capacity adds a second layer of control. Under high IFN induction, the memory–effector response shows rapid initial activation followed by later *p53*-dependent divergence; under low IFN induction, the early phase is weaker or delayed and the outputs accumulate more progressively. Persistent viral burden can still support late activation, especially under stronger *p53* regimes, but it does not fully replace a strong early IFN–JAK/STAT1 drive. Persistence and IFN-induction capacity are thus not equivalent: persistence maintains the stimulus, whereas IFN-induction capacity determines how efficiently viral burden is converted into an early antiviral response.

**Figure 13:**
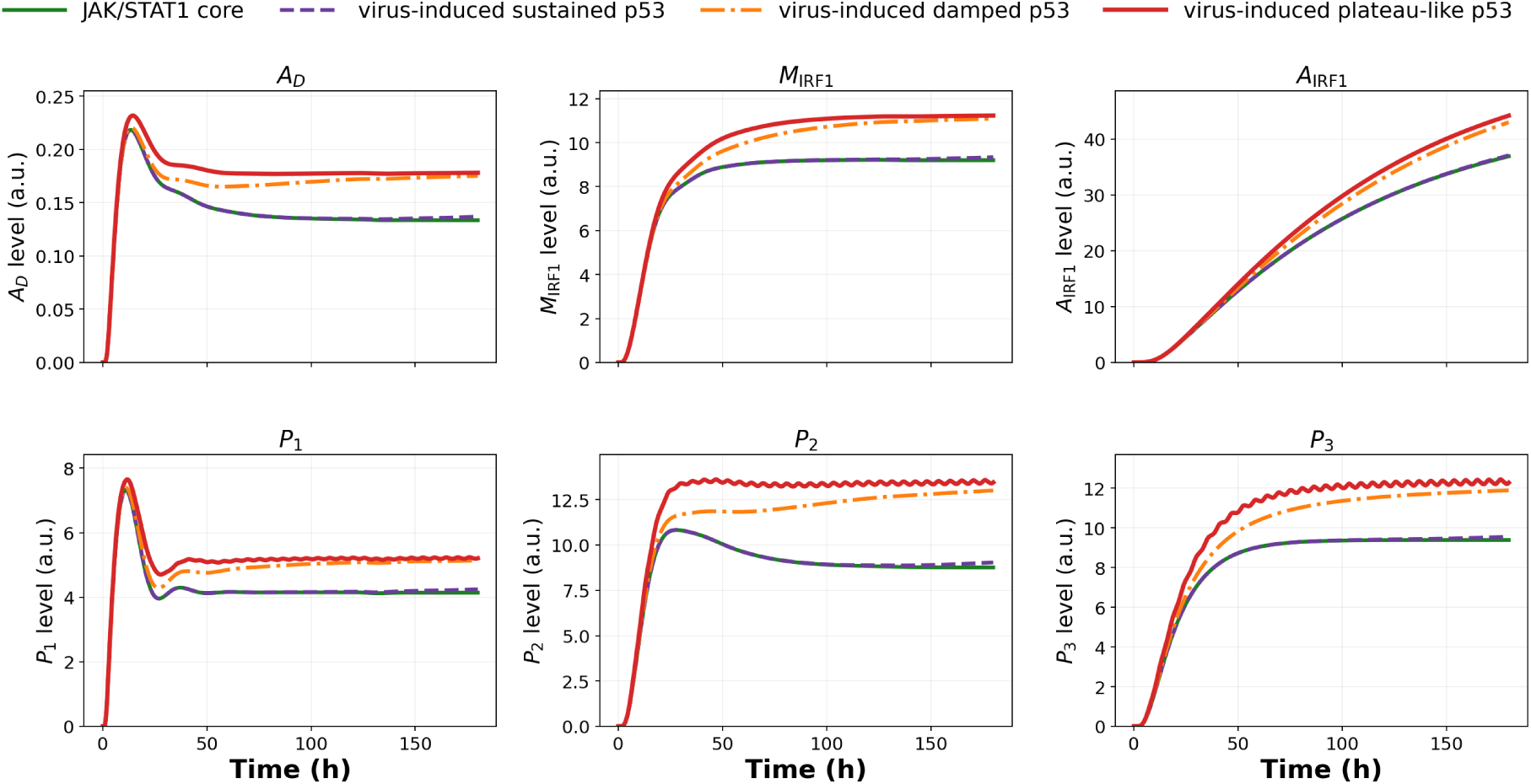
Memory and effector deployment under virus-induced IFN and virus-induced *p53*. The figure shows a highly resistant, strongly IFN-inducing viral context. Curves compare the JAK/STAT1 core condition with virus-induced sustained, damped, and plateau-like *p53* regimes. The upper row shows the memory variables *A_D_*, *M*_IRF1_, and *A*_IRF1_. The lower row shows the antiviral effectors *P*_1_, *P*_2_, and *P*_3_.

The memory–effector layer therefore acts as a dynamical readout of viral class: its trajectories encode several viral properties at once—how efficiently the virus induces IFN, how strongly it induces checkpoint stress, how strongly it resists the effector programme, and how long it persists. This provides the mechanistic link to apoptotic routing, because the same persistent viral burden that reinforces *M*_IRF1_, *A*_IRF1_, *P*_2_, and *P*_3_ also maintains the context in which *P*_4_- or *P*_5_-associated commitment can emerge. Additional scenario simulations are provided in Supplementary Figure S12 and Figure S13.

### 6.2. Virus-induced p53 accelerates P*_4_*-apoptosis only under persistent viral burden

We next asked how virus-induced *p53* reshapes the conversion of JAK/STAT activity into antiviral restriction and apoptotic-route selection. We focused first on the highest calibrated non-saturating viral-to-IFN condition, corresponding to *χ*_IFN_ = 0.1003 and a realised maximal virus-induced IFN level of approximately 45. Under this fixed IFN-induction context, we varied the viral-to-*p53*-stress gain, *χ_p_*_53_ = 0.2, *χ_p_*_53_ = 0.8, and *χ_p_*_53_ = 1.2, across three viral classes: sensitive, intermediate, and highly resistant.

Increasing *χ_p_*_53_ strengthens the conversion of persistent viral burden into *p53*-associated stress, whereas *P*_4_ remains conditional on sustained JAK/STAT–IRF1 signalling in a persistent viral-burden context through Φ*_V_* _4_(*V_g_*). The selected apoptotic route therefore depends on whether viral burden is cleared before commitment, maintained long enough to support *P*_4_, or sustained together with sufficiently strong *p53* stress to recruit the *p53*-autonomous *P*_5_ route.

Figure 14 shows that the effect of virus-induced *p53* is conditioned by whether viral burden persists long enough to keep the *P*_4_ gate open. In the sensitive class, infection is cleared for all three values of *χ_p_*_53_: viral genomes decline rapidly and the danger route cannot remain open long enough for *P*_4_-commitment. Strong antiviral engagement and high IFN induction therefore do not imply apoptosis when the virus is efficiently cleared.

The intermediate class shows the clearest dependence of *P*_4_-timing on virus-induced *p53*. Viral control is incomplete, so *V_g_* persists long enough to maintain the *P*_4_ gate, and as *χ_p_*_53_ increases *P*_4_-commitment occurs earlier. The route, however, is not set by *χ_p_*_53_ alone: at the highest *χ_p_*_53_, *P*_5_ is also recruited, producing dual-route apoptosis. This class is thus a transition regime, where persistent viral burden supports *P*_4_ while stronger virus-induced *p53* stress adds *P*_5_.

**Figure 14:**
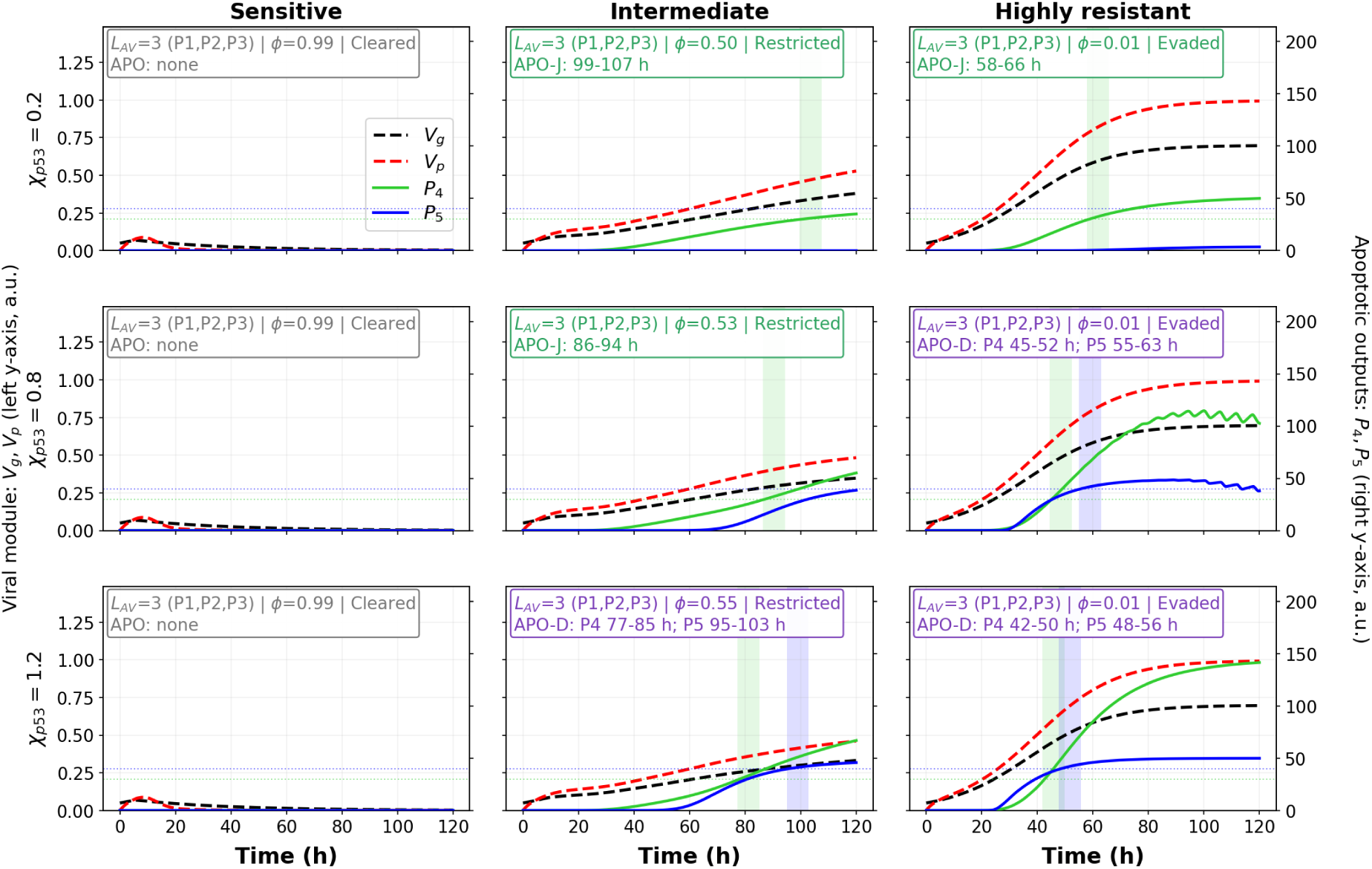
Time-course responses under the highest calibrated non-saturating virus-induced IFN condition, with *χ*_IFN_ = 0.1003 and realised maximal IFN_int_ ≈ 45. Rows show increasing viral-to-*p53*-stress gain, *χ_p_*_53_ = 0.2, 0.8, and 1.2. Columns compare sensitive, intermediate, and highly resistant viral classes. Black and red dashed curves show viral genomes *V_g_* and viral particles *V_p_*, respectively. Green and blue curves show the apoptotic outputs *P*_4_ and *P*_5_. Shaded regions indicate sustained threshold crossing for apoptotic commitment. Decision boxes report the antiviral state, the viral outcome *φ*, and the selected apoptotic route.

The highly resistant class maintains the strongest viral burden and sustains the *P*_4_ gate most robustly: *P*_4_-associated apoptosis is selected even at low *χ_p_*_53_. As *χ_p_*_53_ increases, *P*_5_ is recruited and the response shifts towards dual-route commitment. Resistant infection thus supplies a persistent *P*_4_-permissive danger context, while strong virus-induced *p53* stress adds the *p53*-autonomous *P*_5_ component.

Figure S15 in the supplementary extends this across the continuous *χ*_IFN_ range. Increasing *χ*_IFN_ raises the capacity of viral burden to drive endogenous IFN, and increasing *χ_p_*_53_ raises its capacity to drive *p53*-associated stress; the map shows how these combine with viral sensitivity and route-specific thresholds to produce distinct antiviral and apoptotic outcomes.

The sensitive class stays non-apoptotic across the tested *χ_p_*_53_ values, consistent with the time-course analysis: viral clearance closes the *V_g_*-dependent *P*_4_ gate before sustained commitment is reached, so increasing antiviral engagement does not imply apoptosis once viral burden is resolved.

The intermediate class occupies a transition regime. Viral control is partial and persistent *V_g_* keeps the *P*_4_ gate open, so the response is mainly *P*_4_-associated at low *χ_p_*_53_ and shifts, over part of the parameter range, towards dual-route apoptosis as *χ_p_*_53_ increases. This is one of the main results of the analysis: the same persistent viral burden that supports delayed *P*_4_-commitment can, under stronger virus-induced *p53* stress, combine with *P*_5_-associated commitment.

The highly resistant class is dominated by persistent viral burden, so the *P*_4_ gate remains active over a broad range of *χ*_IFN_. Increasing *χ_p_*_53_ then recruits *P*_5_ and shifts the response towards APO-D, indicating that highly resistant infection sustains both arms of the apoptotic programme: a *V_g_*-gated JAK/STAT–IRF1 danger route and a virus-induced *p53*-dependent route.

Together, the time-course and response-map analyses show how the infection-coupled architecture translates viral persistence into functional routing. Viral burden is both a target of antiviral effectors and an input to the host-response network: persistent *V_g_* sustains virus-induced IFN, virus-induced *p53* stress, and the *P*_4_ danger gate Φ*_V_* _4_(*V_g_*).

This creates a class-dependent separation between viral control and apoptotic routing. Sensitive viruses are cleared before sustained apoptotic commitment is reached. Intermediate viruses occupy a transition region in which delayed *P*_4_-associated commitment can emerge and, at higher *χ_p_*_53_, shift towards dual *P*_4_*/P*_5_ commitment. Highly resistant viruses maintain the persistent viral-burden context required for *P*_4_ and, when virus-induced *p53* stress is sufficiently strong, also recruit the *p53*-autonomous *P*_5_ route.

This organisation was preserved under gate-parameter variation (Supplementary Section S8, indicating that the routing landscape is organised by viral persistence and viral-to-*p53* stress induction rather than by a single nominal gate-parameter choice.

## 7. Conclusion

This study provides a dynamical interpretation of how *p53* reshapes an IFN-*γ*-centred JAK/STAT1 antiviral-response network. The central conclusion is not that *p53* simply strengthens or weakens JAK/STAT1 signalling, but that it changes how an activated STAT1 signal is temporally decoded and functionally routed. By placing the *p53*-dependent coupling downstream of STAT1 activation, the model isolates a specific hypothesis: *p53*-associated stress history modifies nuclear STAT1 processing, transcriptional persistence, downstream effector transmission and apoptotic competence, rather than acting only through IFN availability or receptor-proximal phosphorylation.

At the signalling level, *p53*-dependent coupling redistributes the IFN-induced response towards DNA-bound STAT1 persistence, integrated transcriptional output and STAT1-driven feedback. This produces a persistence–recovery trade-off: the same memory-dependent mechanism that prolongs the transcriptionally active STAT1 state also delays pathway re-inducibility after repeated IFN stimulation. Thus, prior *p53*-associated stress can make the JAK/STAT1 response more persistent but less rapidly recoverable.

When this reshaped signalling core is propagated to the downstream memory–effector layer, the effect of *p53* is filtered rather than transmitted as a simple gain. Prescribed *p53* strengthens IRF1-associated memory and antiviral effector production, but increasing *p53* preactivation does not produce a proportional increase in all downstream antiviral outputs. The antiviral programme therefore transforms upstream STAT1 priming through memory accumulation, IRF1-associated integration, effector production and turnover.

At the functional level, the model separates antiviral-state engagement from realised viral control. The normalised antiviral-state score *L*_AV_ measures sustained activation of the host effector programme, whereas the viral-control score *φ* measures whether that programme actually suppresses late viral burden. These two quantities need not coincide. A strong antiviral state can coexist with viral evasion when the viral class is poorly sensitive to the induced effectors, whereas sensitive viral classes can be cleared under a comparatively modest antiviral state. Infection outcome is therefore determined by the match between host effector activity and viral sensitivity, not by IFN amplitude or STAT1 activation alone.

The origin of the IFN and *p53* inputs provides a further level of organisation. Externally imposed IFN or *p53* stimulation probes the intrinsic host-side capacity of the pathway, whereas virus-induced IFN and virus-induced *p53* stress remain dynamically coupled to viral persistence. Persistent viral burden is therefore both a target of antiviral restriction and an input that sustains endogenous IFN production, checkpoint stress and apoptotic routing. Consequently, signal composition cannot be reduced to a single total IFN amplitude: changing the balance between imposed and virus-coupled sources can modify late viral burden and route-specific apoptotic commitment.

Apoptotic commitment is route-specific. The *P*_4_ route represents a JAK/STAT–IRF1-associated danger programme that requires sustained antiviral signalling in a persistent viral-burden context through Φ*_V_* _4_(*V_g_*). The *P*_5_ route represents a more direct *p53*-autonomous apoptotic programme driven by sustained checkpoint activation. Thus, *p53* does not decide apoptosis alone. It conditions apoptotic competence, whereas viral persistence determines whether this competence is routed towards JAK/STAT–IRF1-associated, *p53*-autonomous or dual apoptotic commitment. This explains the class-dependent behaviour of the model: sensitive viral classes are cleared before sustained commitment develops, intermediate classes can support delayed *P*_4_-associated commitment, and highly resistant classes maintain a persistent danger context that can recruit both *P*_4_ and *P*_5_.

These conclusions should be interpreted within the coarse-grained nature of the model. The variables *P*_1_–*P*_5_ are functional axes rather than molecular species, and the threshold-based decisions are phenomenological criteria for sustained activation. Viral classes are passive infection contexts defined by fixed traits: IFN-induction capacity, *p53*-stress induction and sensitivity to antiviral effectors. The model does not resolve detailed pathogen sensing, *p53* target-gene networks, cell-to-cell variability, paracrine IFN exchange, immune-cell recruitment or tissue-level dynamics. It should therefore be read as a mechanistic framework for qualitative dynamical organisation, not as a point-calibrated model for a specific virus or cell type.

Despite these limitations, the model makes experimentally testable predictions. First, perturbing *p53* should affect STAT1 DNA-bound persistence, temporal integration and recovery more strongly than the early receptor-proximal phosphorylation peak. Second, antiviral-state engagement and viral control should be dissociable across viral classes with different effector sensitivities. Third, *p53* preactivation should show sub-proportional transmission to antiviral effector outputs once the memory–effector layer is engaged. Fourth, JAK/STAT–IRF1-associated apoptotic commitment should depend on persistent viral burden, whereas *p53*-autonomous commitment should track sustained checkpoint stress.

Overall, the *p53*-dependent gates encode a local hypothesis about post-phosphorylation coupling, but their importance lies in their system-level consequences. In this framework, *p53* converts the JAK/STAT1 module from a pathway that simply responds to interferon into a context-dependent decision module that integrates signalling history, effector sensitivity and viral persistence to determine antiviral control and apoptotic fate.

---

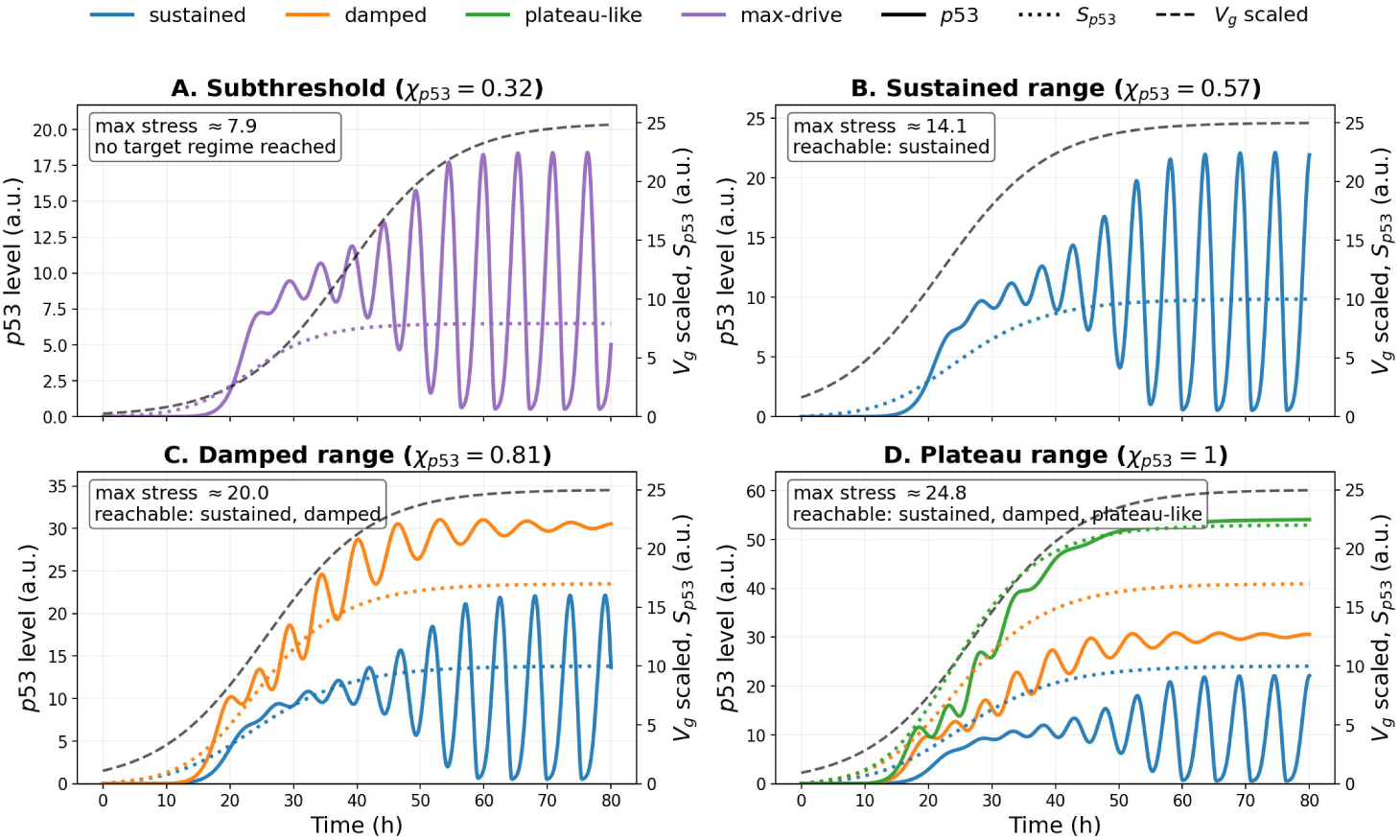

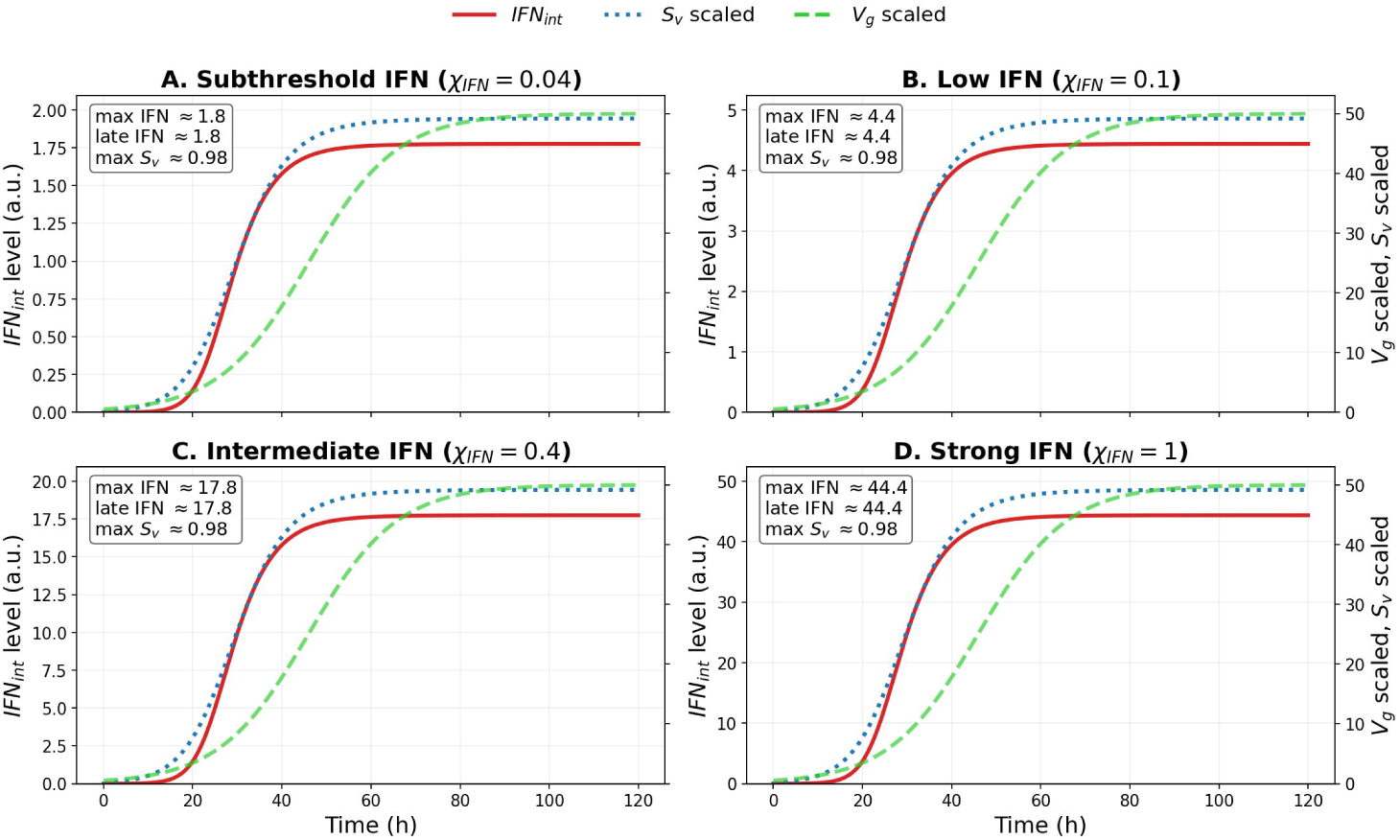

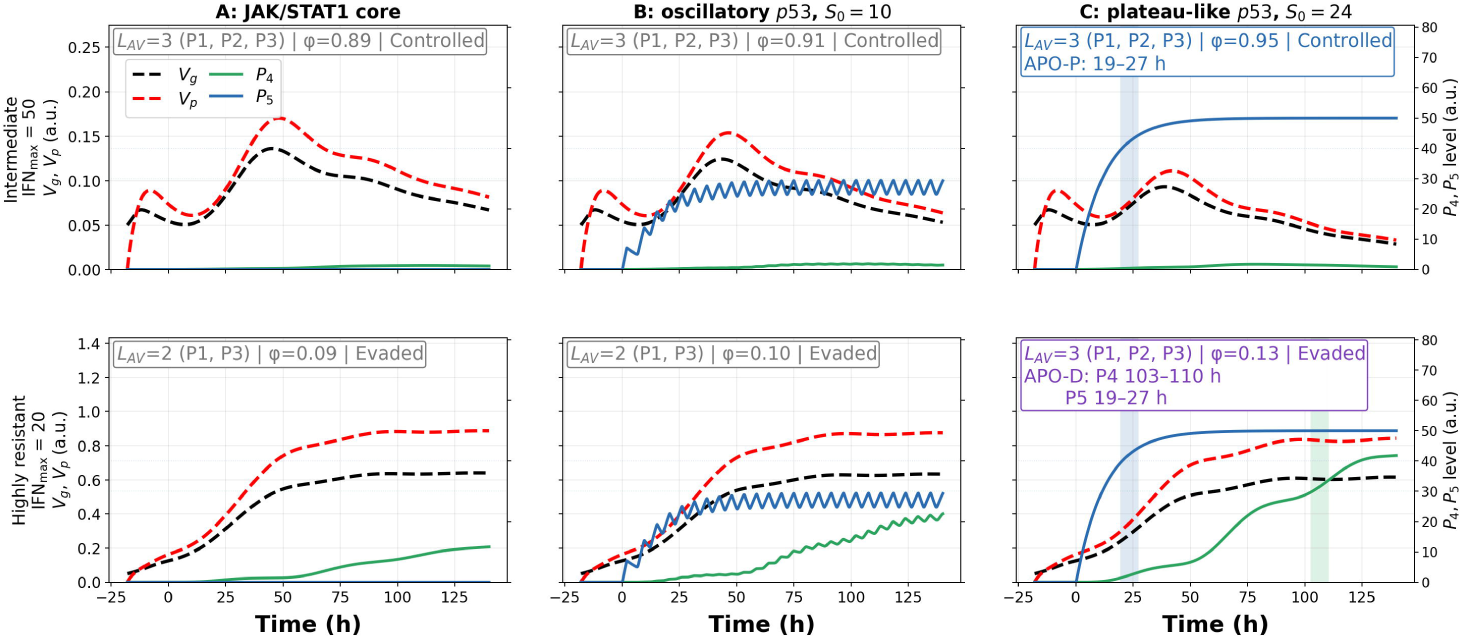

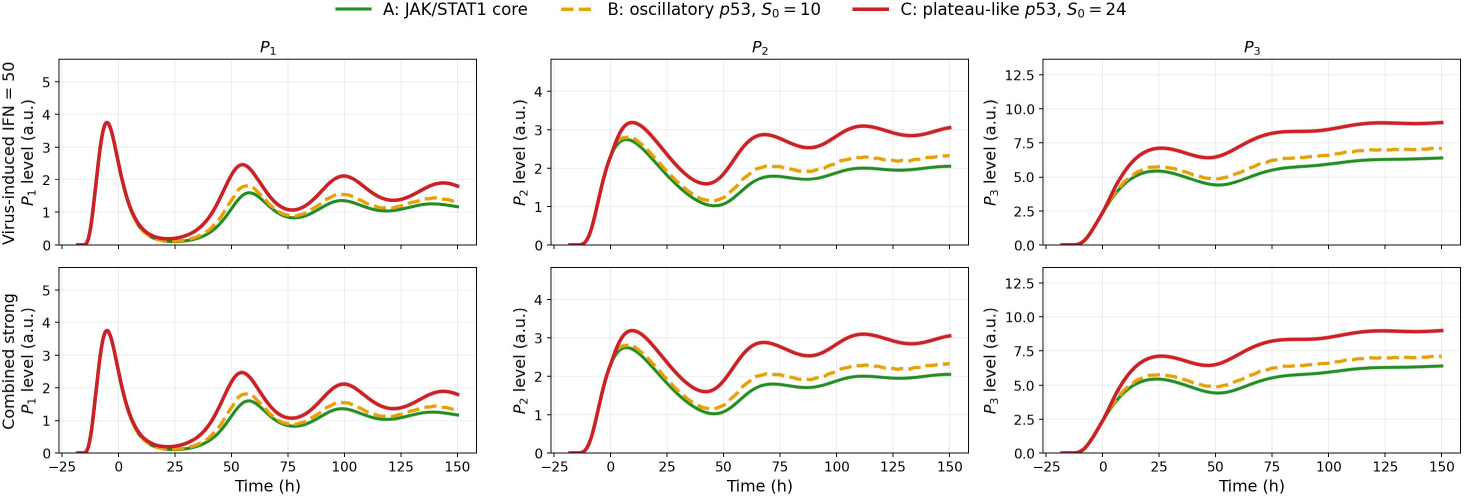

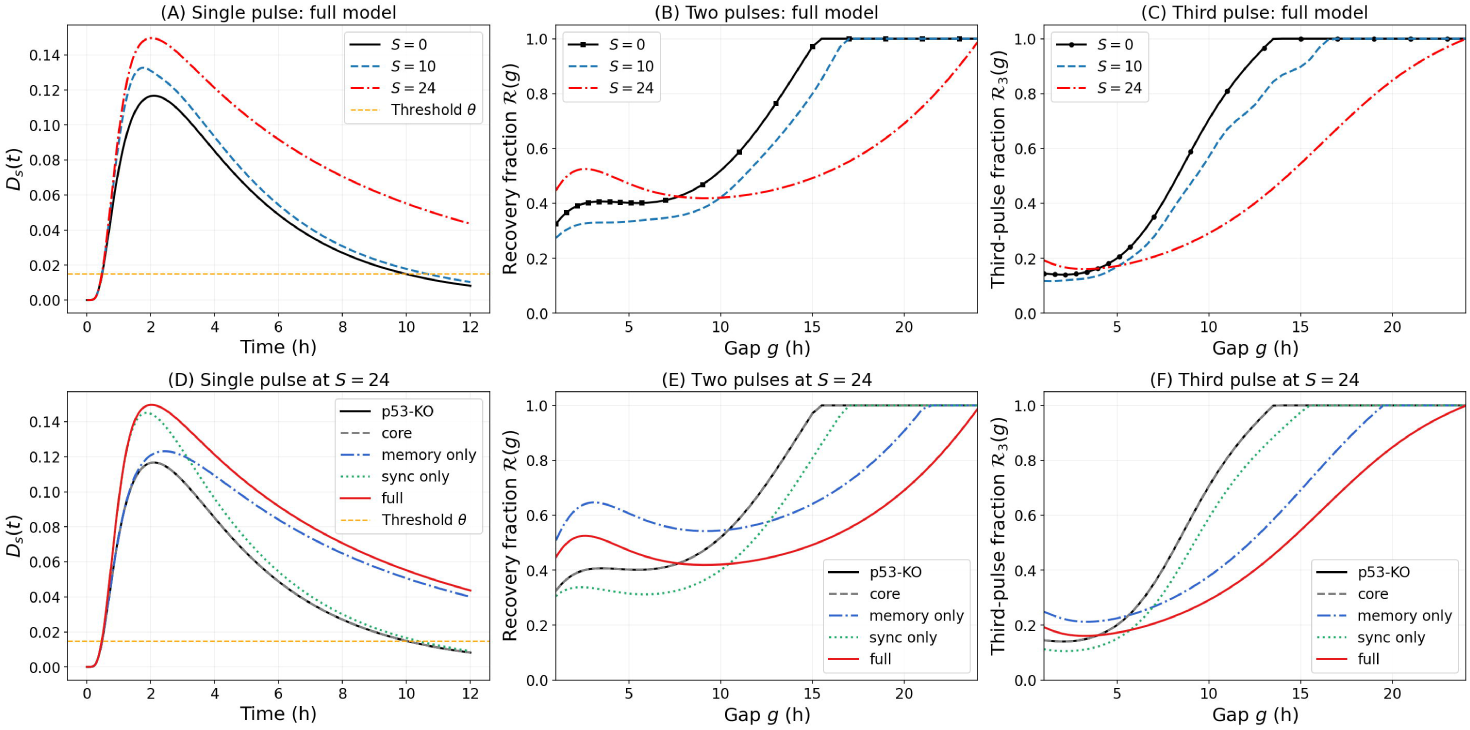

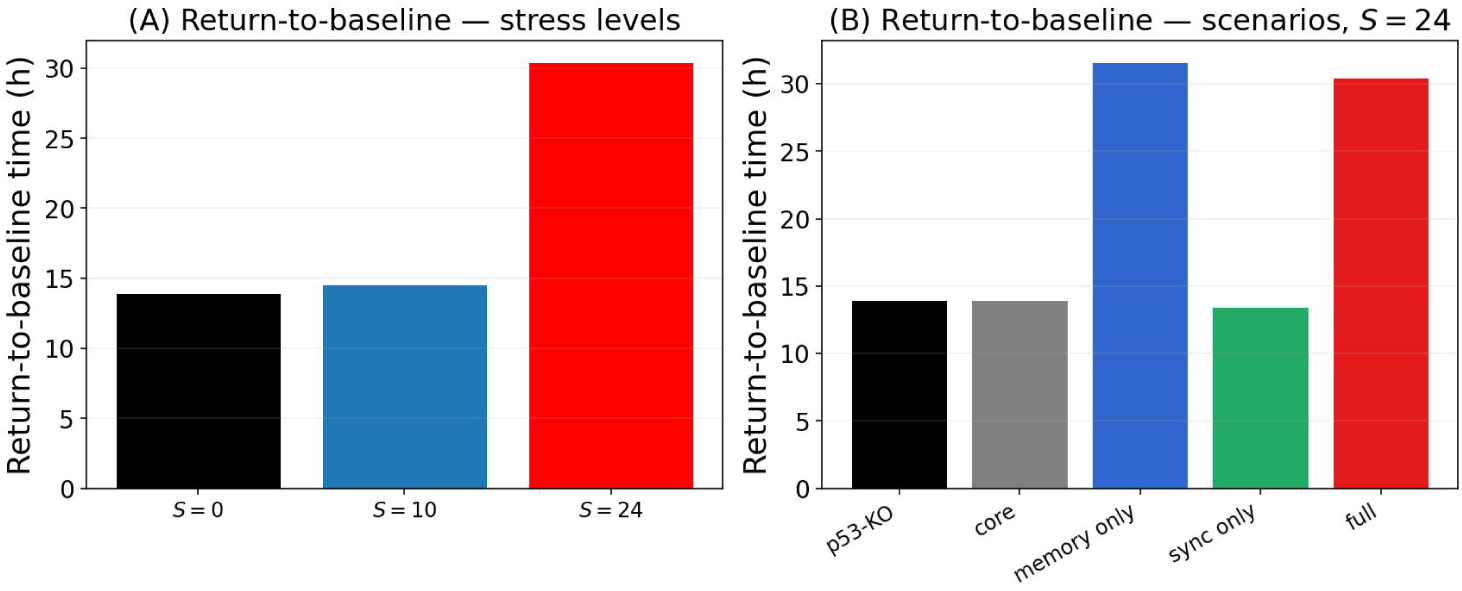

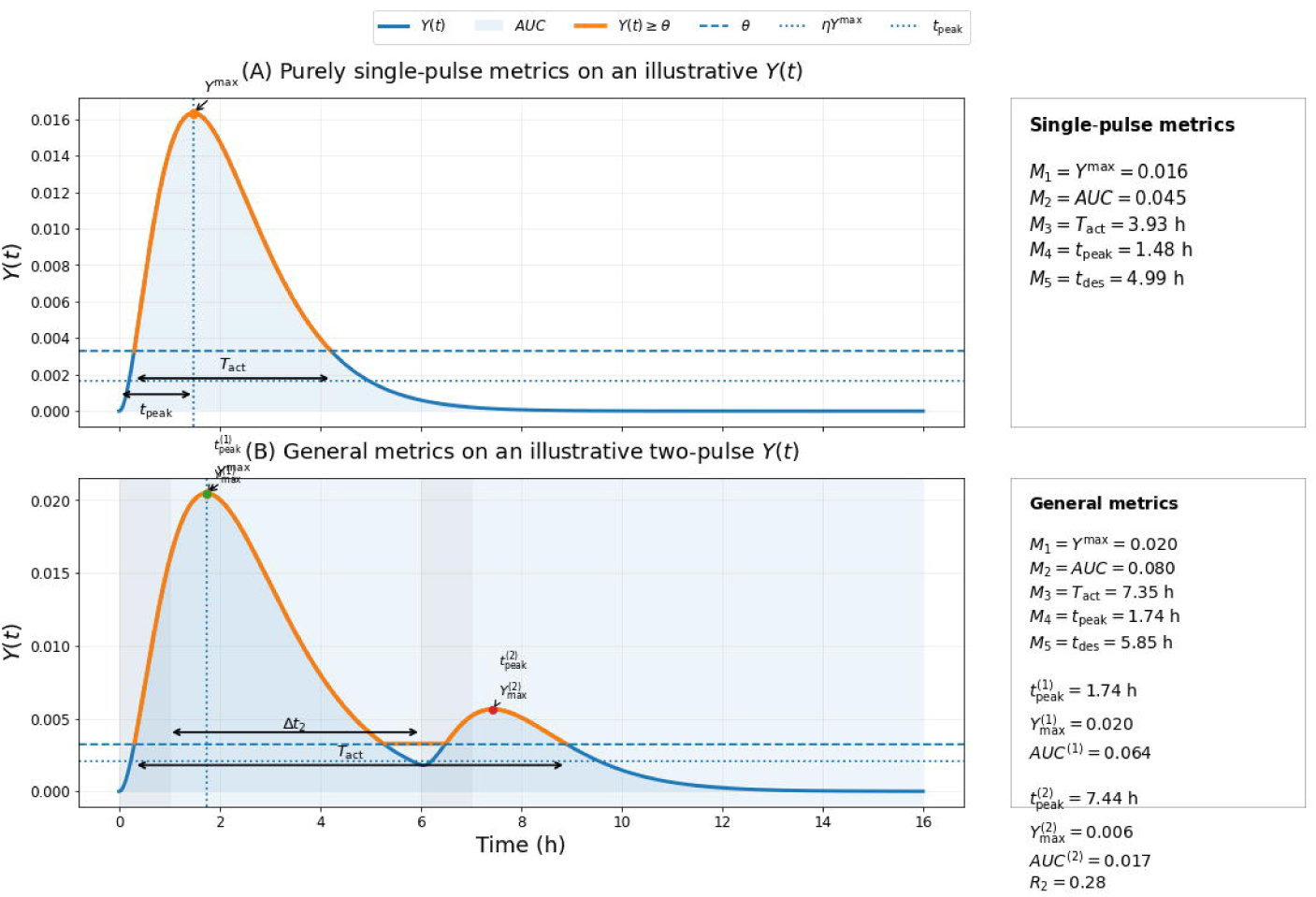

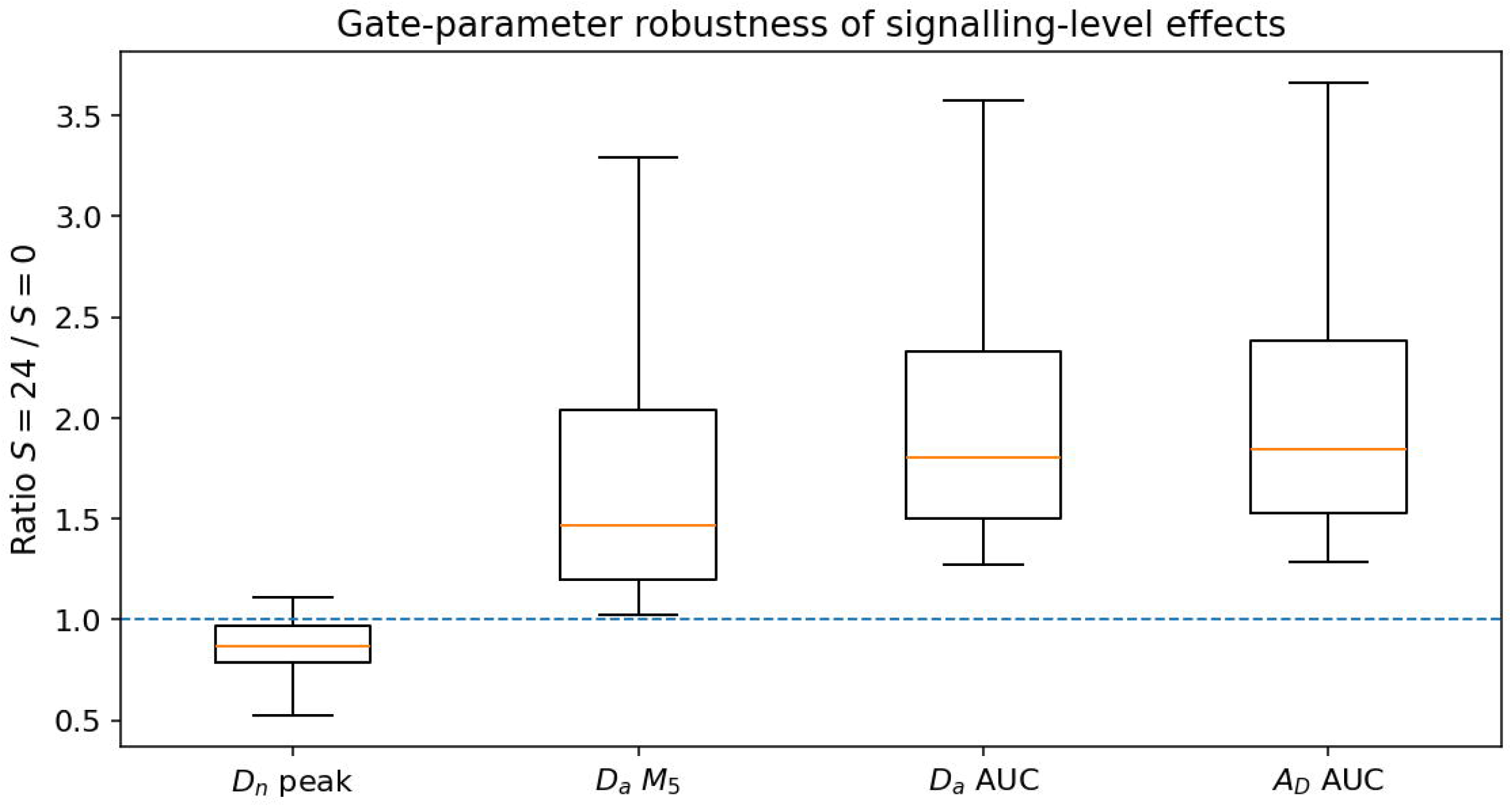

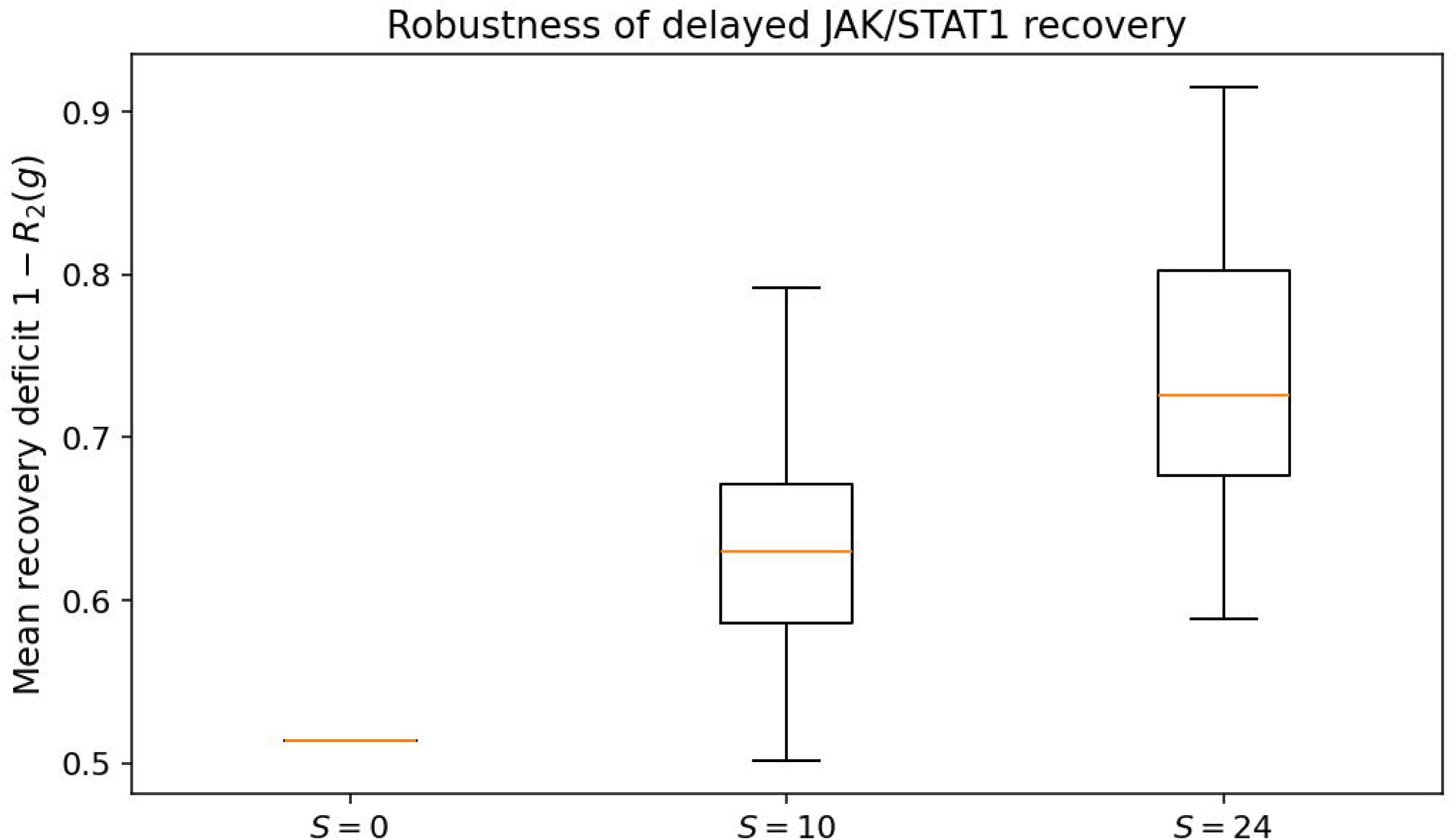

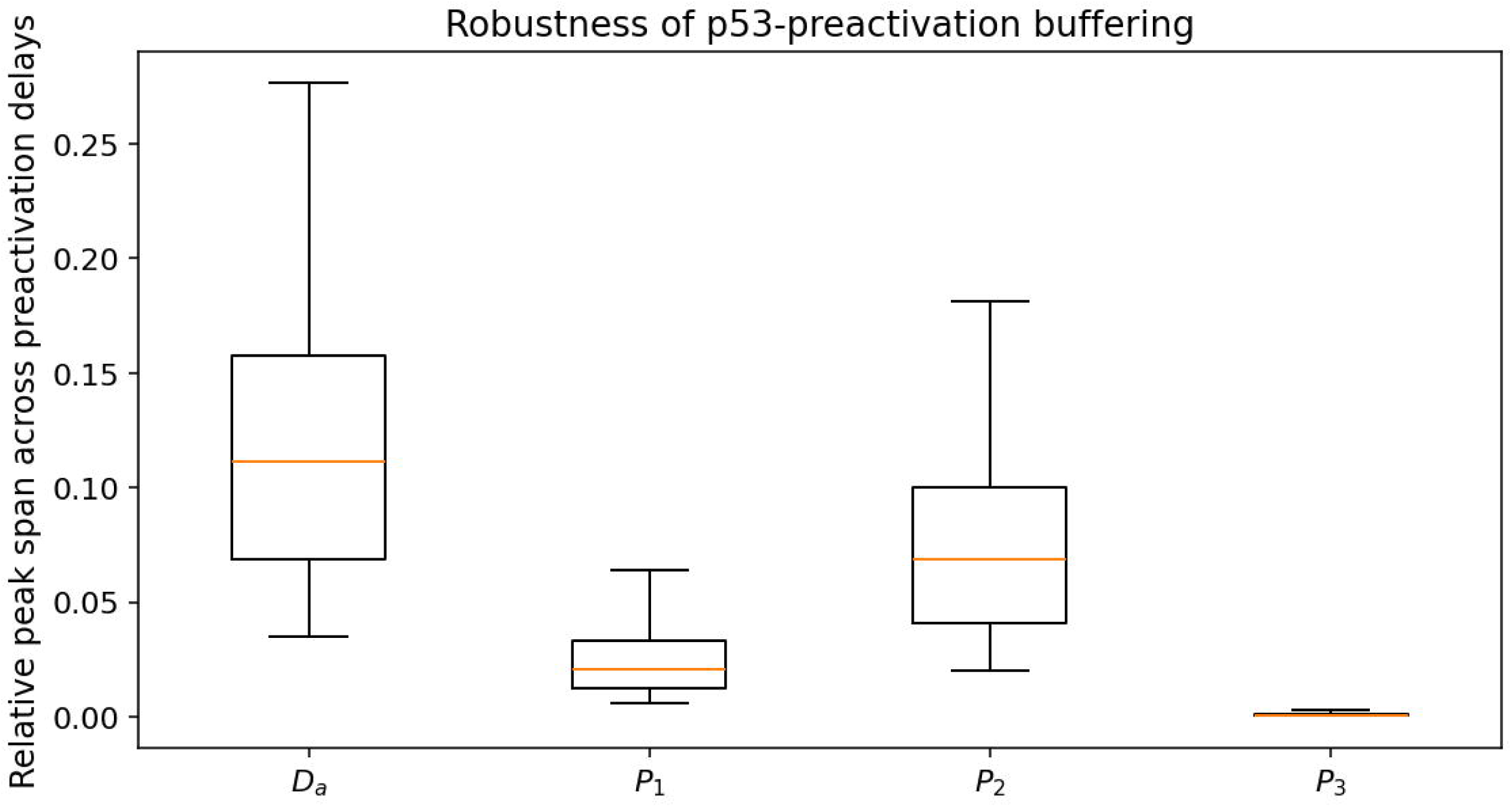

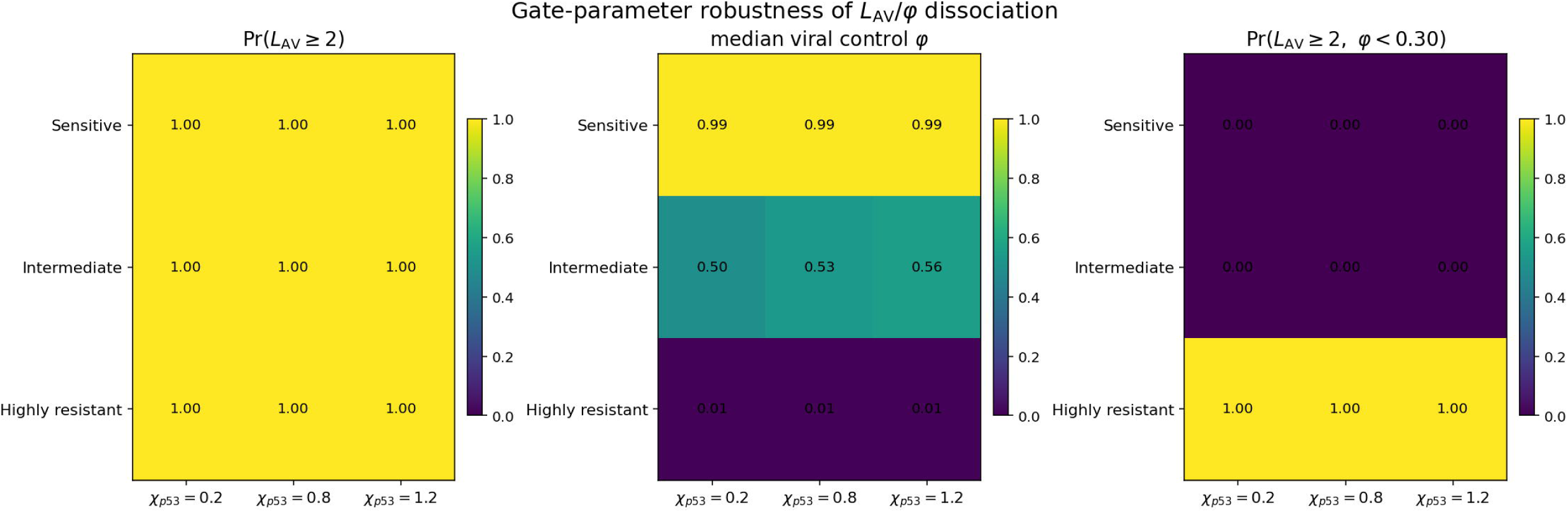

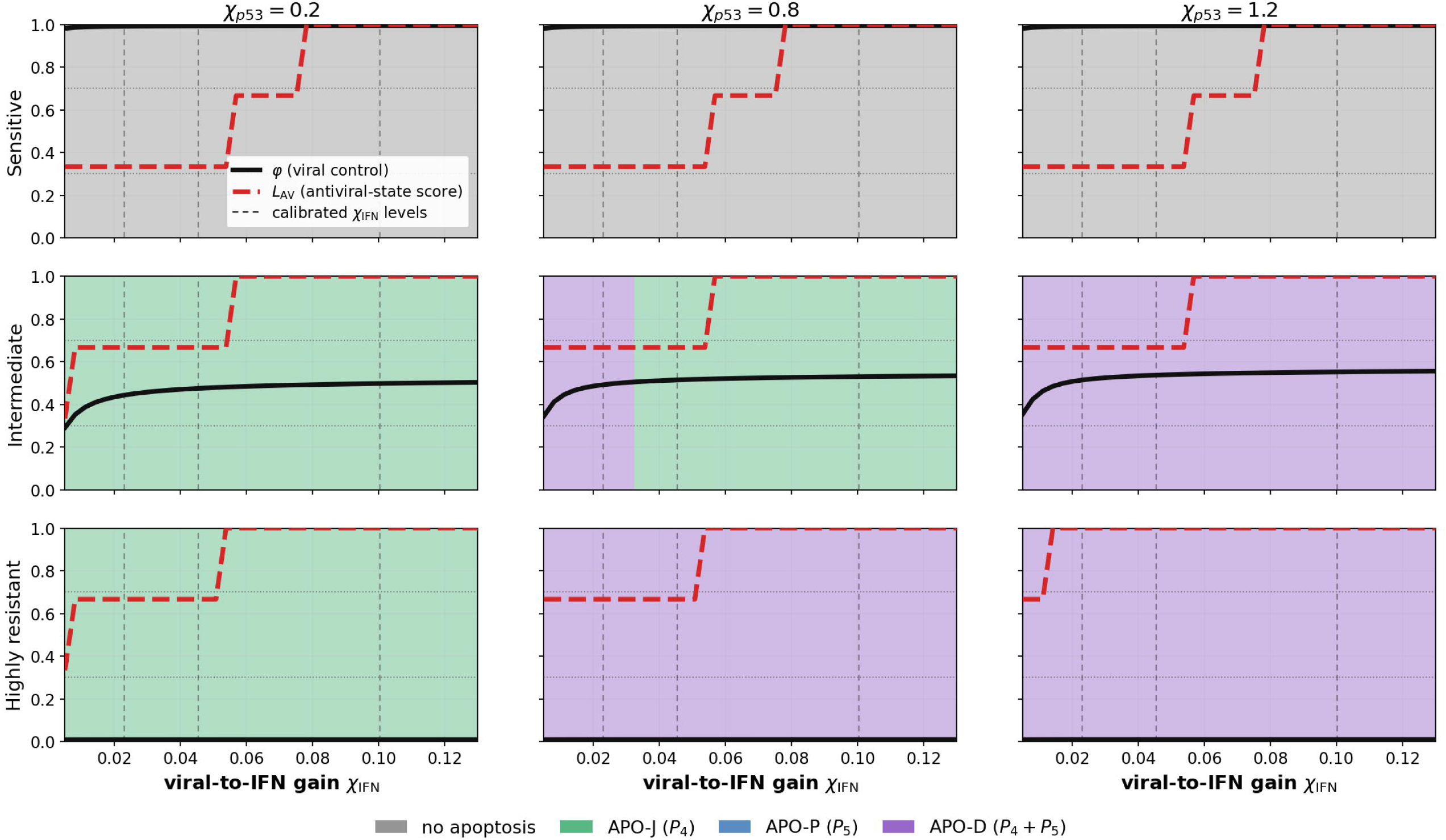

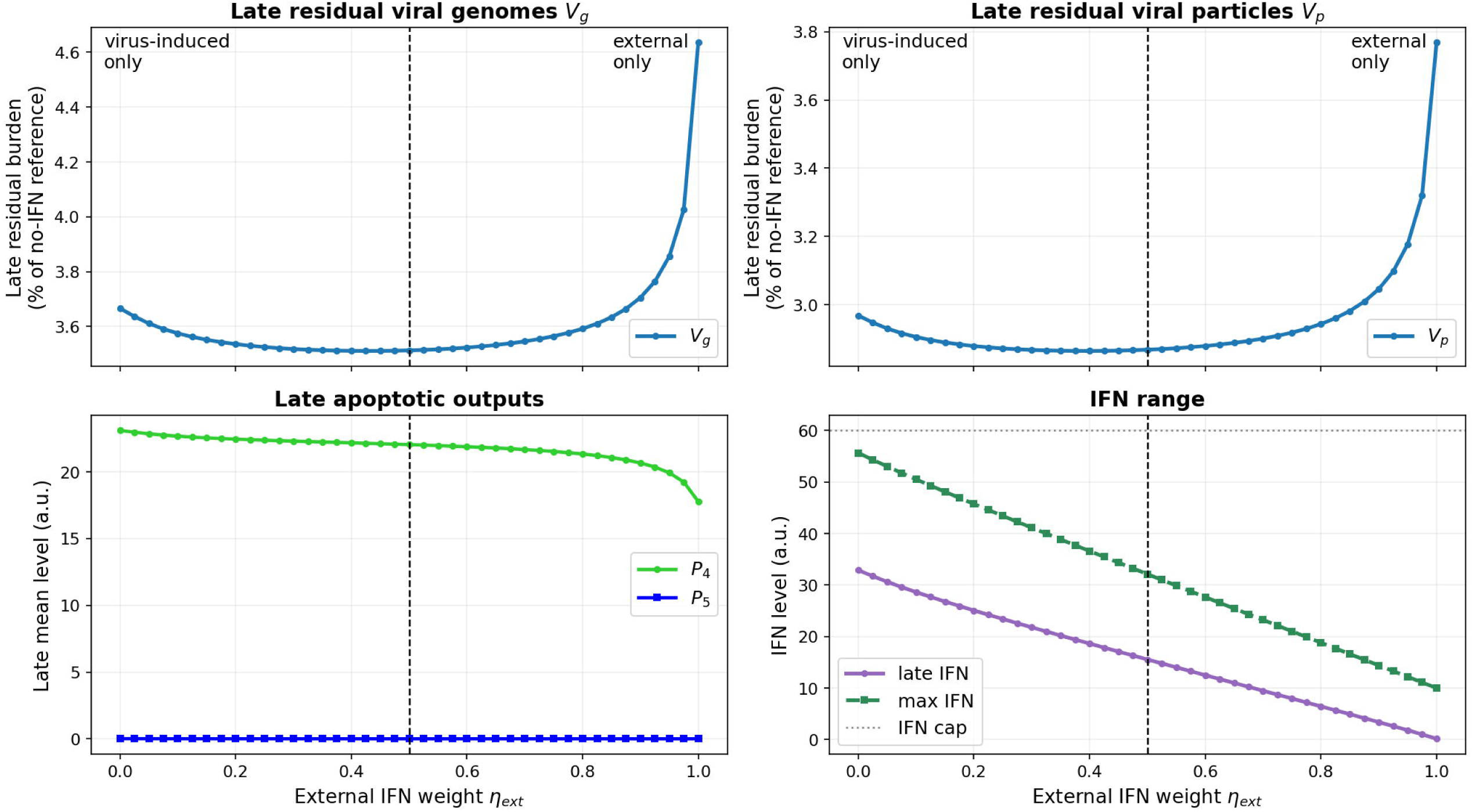

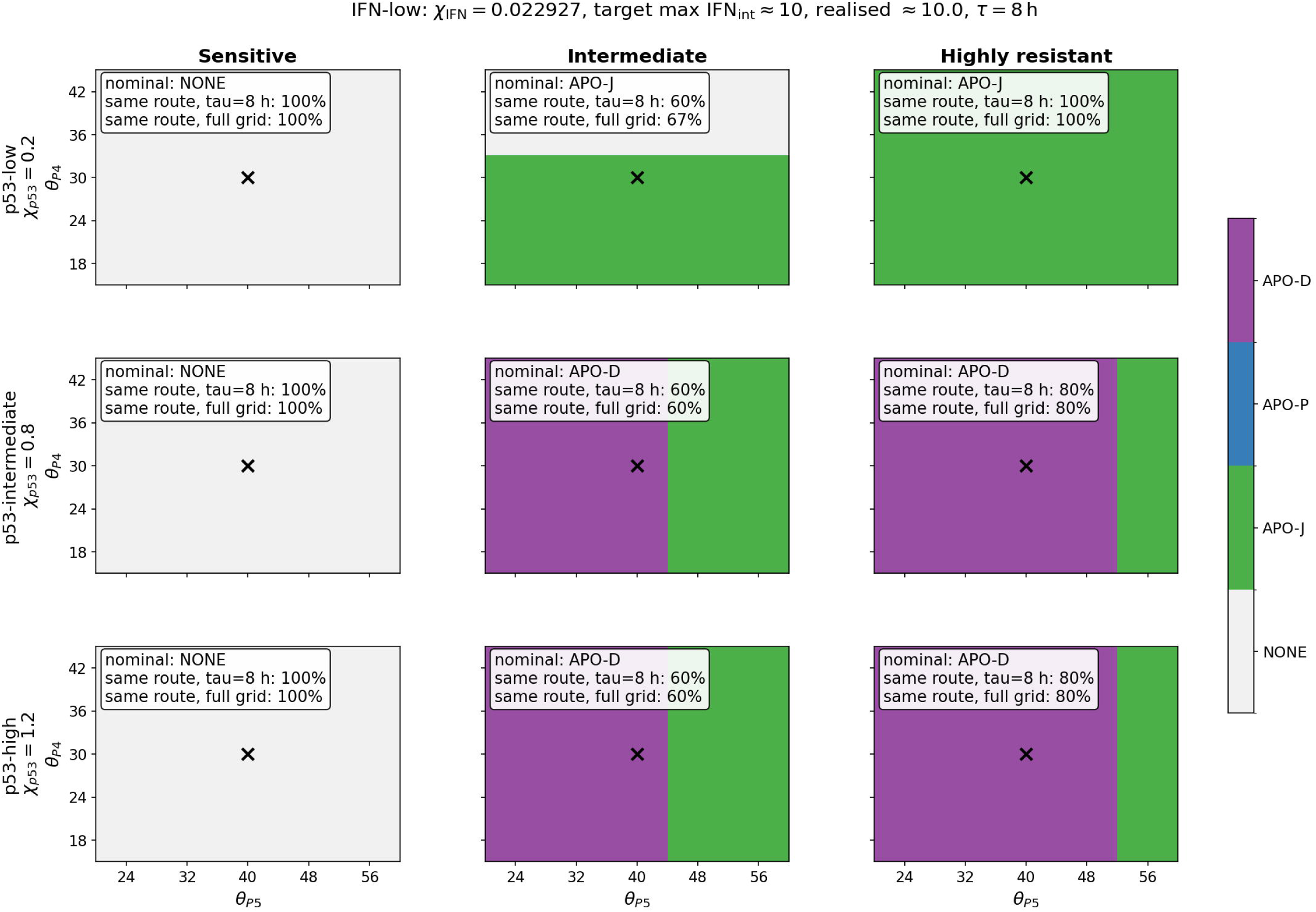

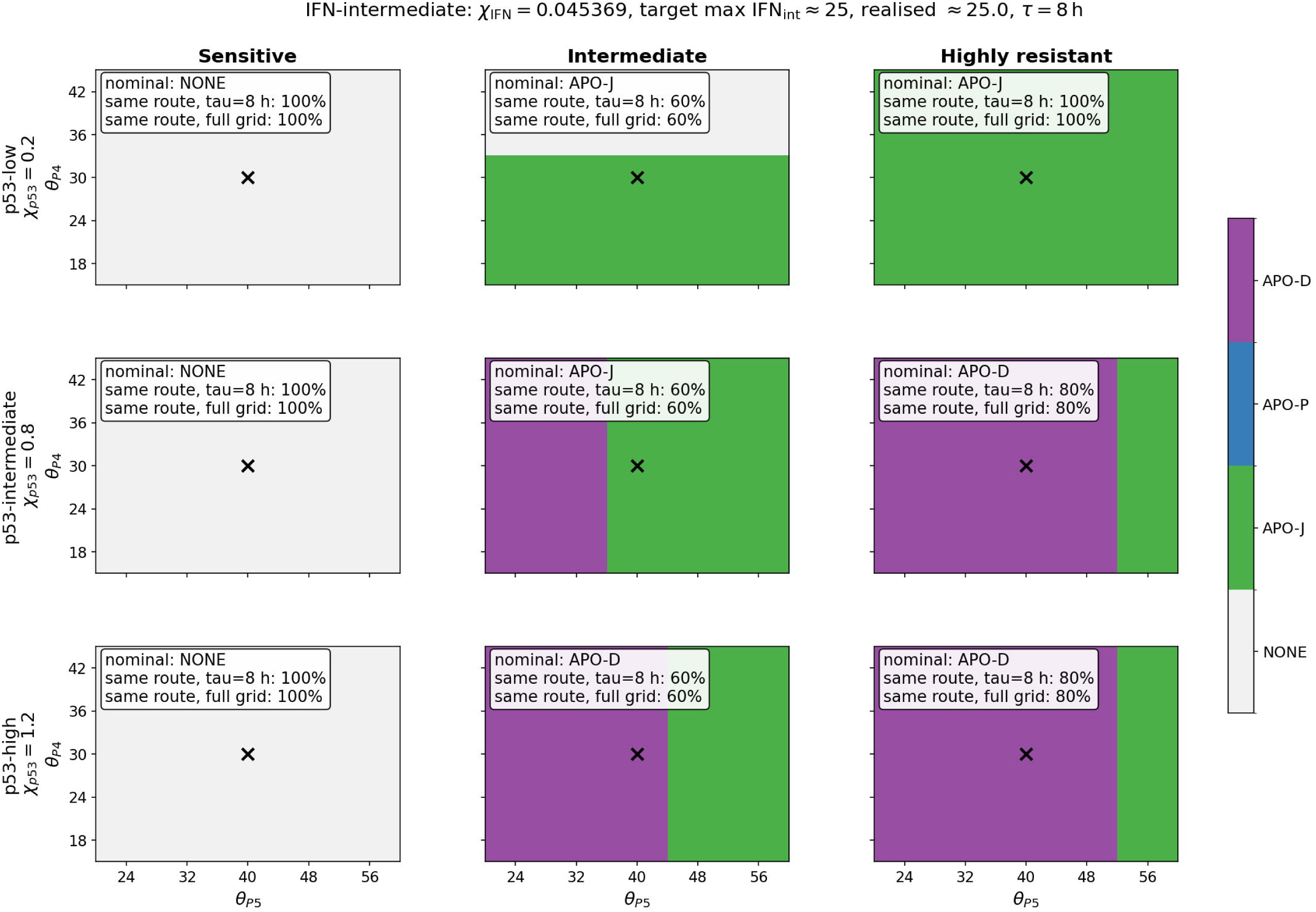

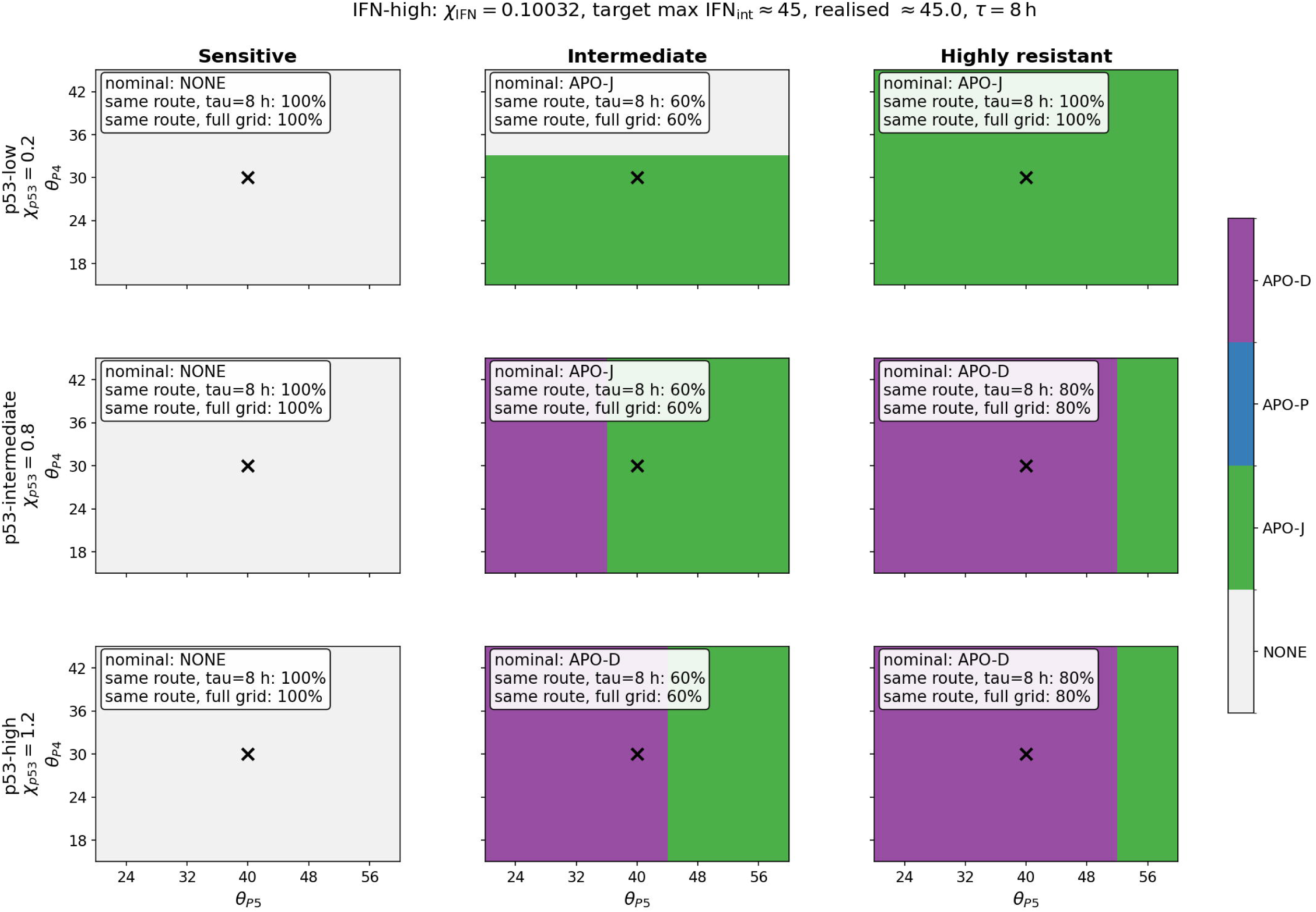

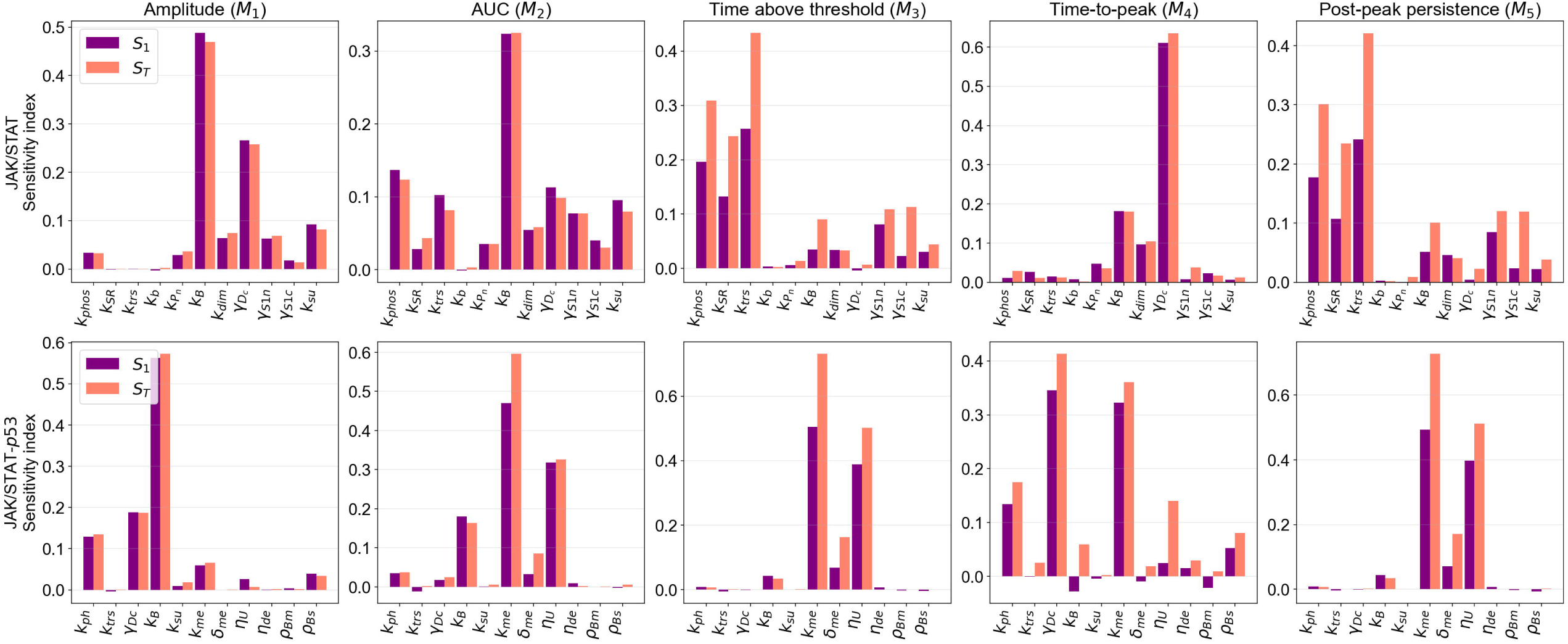

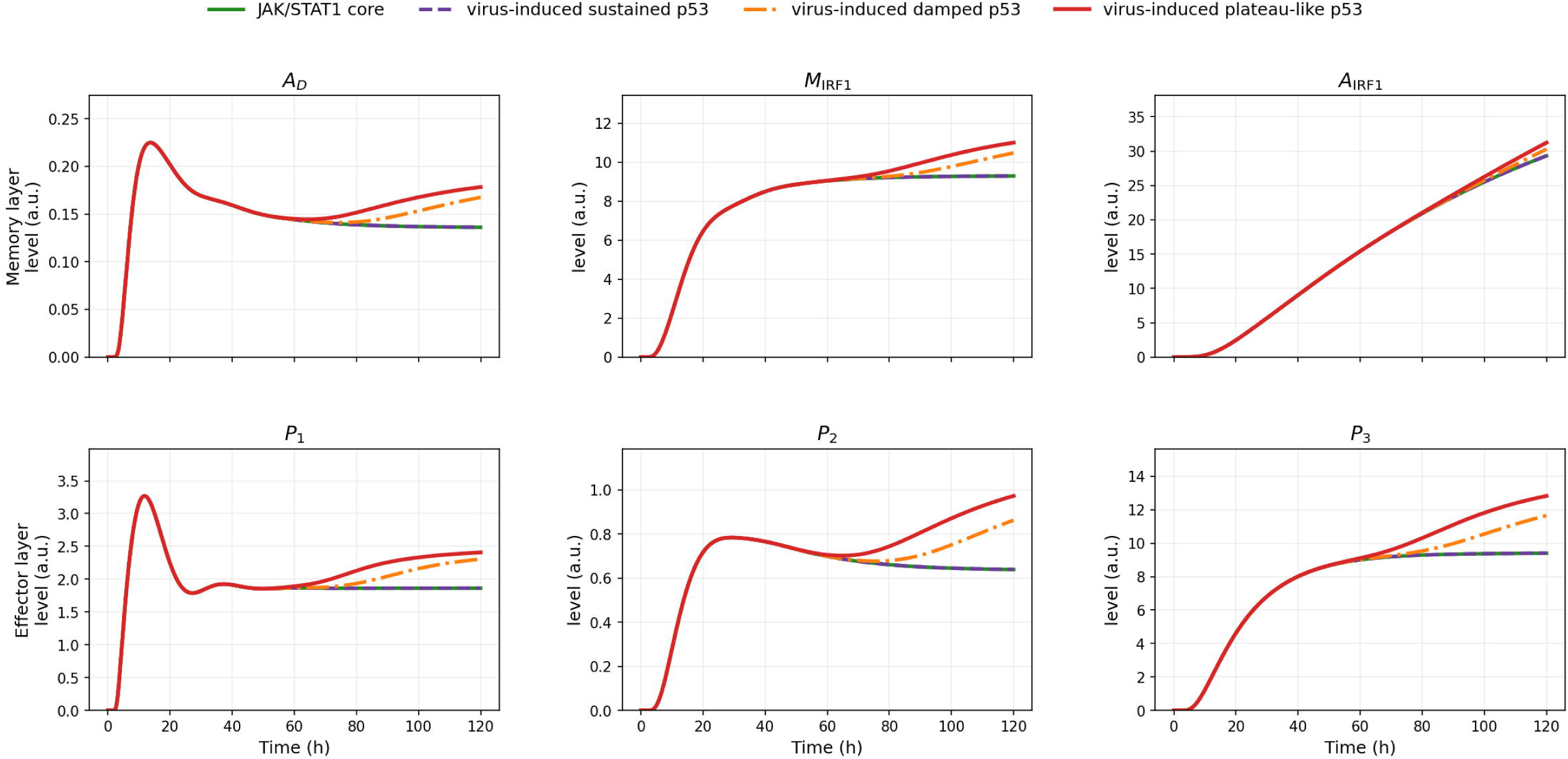

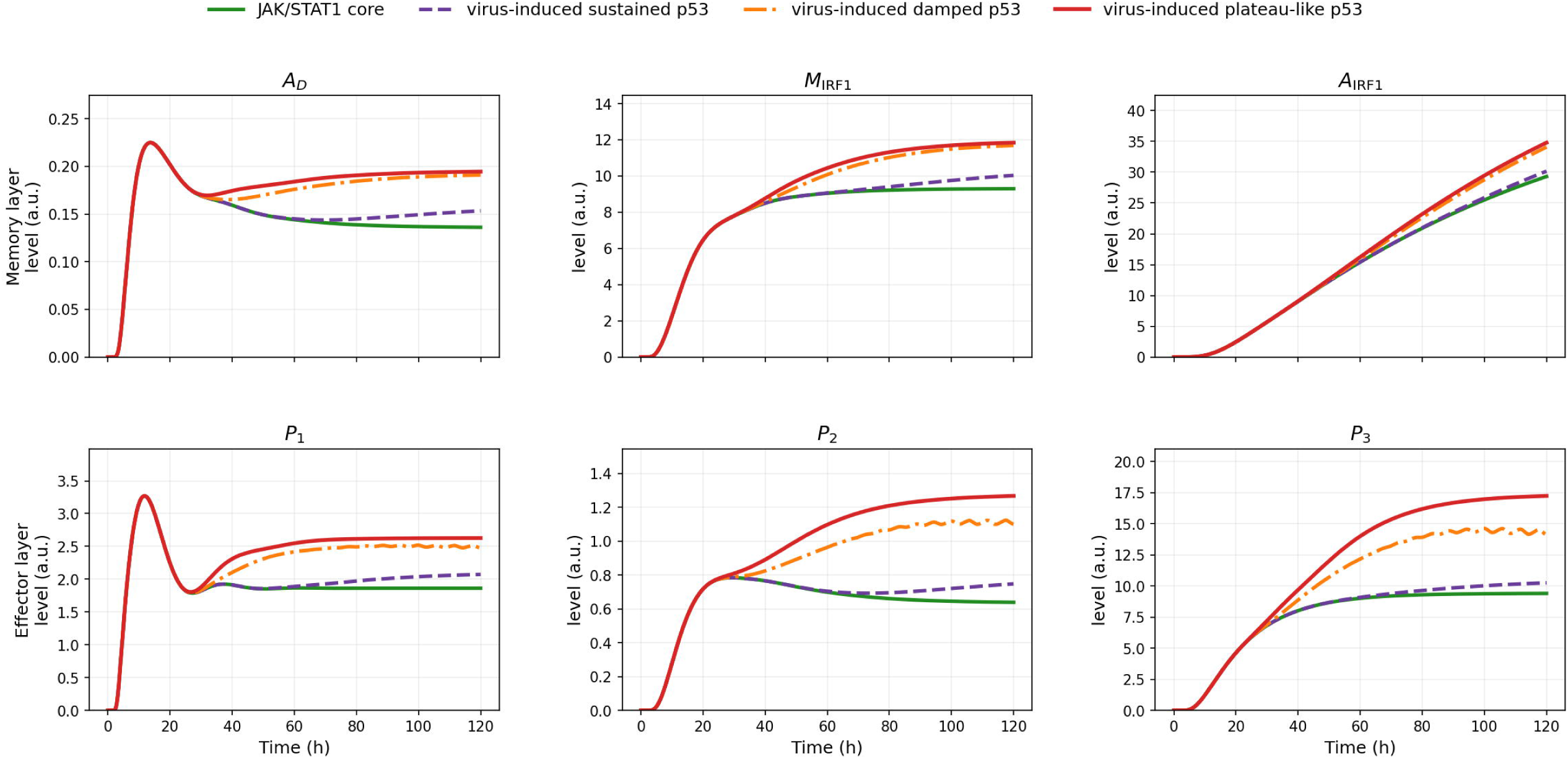

## Notes

### Competing Interest Statement

The authors have declared no competing interest.

