## supplementary material for "How *p53* stress memory could redirect JAK/STAT1 antiviral signalling: a model-based prediction"


### S1. Model description

The model is organised into six equation blocks, grouped into five functional modules: endogenous IFN sensing $(IFN)$, cytosolic and nuclear JAK/STAT1 signalling $(C)$–$(N)$, the *p53–Mdm2* stress-response module $(p53)$, the downstream effector and apoptotic-commitment layer $(E)$, and the intracellular viral-dynamics module $(V)$. The full model is written in block form in the main text, with the six dynamical blocks $(IFN)$, $(C)$, $(N)$, $(p53)$, $(E)$, and $(V)$ given in Equations (2.1)–(2.6). The corresponding auxiliary functions are introduced immediately after the block in which they are used. The parameter values used in the simulations are summarised in Tables S1–S2.

The construction combines components derived from our previous STAT1–*p53* model with new coupling, antiviral-effector and viral-burden extensions. The *p53–Mdm2* oscillator is derived from our earlier dynamical model of STAT1–*p53* coordination, in which STAT1 activity modulated the *p53* network and influenced the balance between p21- and PUMA-associated cell-fate programmes under stress (Tshianyi et al., 2026). Here, the STAT1-dependent contribution to the *p53* module is kept as a fixed background configuration, so that the present model does not impose a fully bidirectional JAK/STAT1–*p53* feedback loop. This allows the analysis to focus on the complementary direction: how *p53*-dependent stress signalling reshapes a JAK/STAT1-centred response.

The JAK/STAT1 module is extended by a *p53*-dependent regulatory layer acting through memory, synchronisation, effector-amplification and apoptotic-competence gates. These gates are phenomenological functions rather than single molecular reactions. They formalise the hypothesis that prior or concurrent *p53*-associated stress can modify post-phosphorylation STAT1 decoding, including nuclear accumulation, DNA-bound persistence, transcriptional memory, downstream effector engagement and apoptotic competence. This modelling choice is motivated by reported interactions between STAT1 and *p53*, and by broader evidence that *p53* intersects with interferon-dependent signalling (Townsend et al., 2004; Youlyouz-Marfak et al., 2008; Będzińska et al., 2025).

The viral component is introduced as a functional extension of the coupled host signalling core. Intracellular viral genomes $V_g$ are converted into two host-response inputs: endogenous IFN production and *p53*-associated checkpoint stress. The parameters $\chi_{\text{IFN}}$ and $\chi_{p53}$ control these two conversions independently. Thus, the model can distinguish viral contexts that efficiently induce IFN from those that preferentially induce checkpoint stress. The total IFN input combines an external source and a virus-induced source,

$$IFN(t) = \eta_{\text{ext}} IFN_{\text{ext}}(t) + \eta_{\text{int}} IFN_{\text{int}}(t),$$

where $\eta_{\text{ext}}$ and $\eta_{\text{int}}$ define the IFN-source composition.

Throughout the model, repeated mathematical structures have the same interpretation. First-order loss terms have the form $-\delta_x x$, with $\delta_x > 0$. Saturating activation terms are represented by Hill-type functions. Inhibitory functions reduce effective activity, whereas amplification factors


*Corresponding author.
*Email addresses:* (Francis Didier Tshianyi M.K.), (Rebecca Walo Omana), (Apollinaire Ndondo Mboma), (Didier Kumwimba Seya), (Didier Gonze)
[1] Present address: Unit of Theoretical Chronobiology, ULB, Brussels, Belgium.

multiply basal production, binding, transport or degradation processes. This convention allows recurrent kinetic mechanisms to be used consistently across the model.

*Endogenous IFN sensing*

The endogenous IFN module converts intracellular viral burden into a virus-induced IFN input. Viral genomes $V_g$ activate the sensing variable $S_v$ through a saturating term scaled by $\chi_{\mathrm{IFN}}$, interpreted as the viral-to-IFN induction gain. Low values correspond to weak activation of the sensing axis, whereas high values correspond to efficient conversion of viral burden into IFN production.

The sensing variable $S_v$ contains a positive-feedback term $(1 + \alpha_S S_v)$, representing amplification of innate sensing once the response is engaged. This amplification is limited by the delayed variable $E_v$, which represents coarse-grained exhaustion or adaptation under sustained viral burden. The endogenous IFN signal $IFN_{\mathrm{int}}$ is then produced from $S_v$ through a saturating activation function and decays with rate $\delta_{\mathrm{IFN}}$. Thus, unlike an externally prescribed IFN pulse, $IFN_{\mathrm{int}}$ depends dynamically on the infection trajectory.

*Cytosolic and nuclear JAK/STAT1 signalling*

The JAK/STAT1 module converts the total IFN signal into nuclear STAT1 transcriptional activity. In the cytosolic block ($C$), free receptors $R_f$ bind IFN and enter the activated state $R_a$. Activated receptors promote STAT1 phosphorylation through the term $v_{\mathrm{phos}} R_a S1_c$, with $v_{\mathrm{phos}} = k_{\mathrm{phos}} h_{\mathrm{SOCS1}}$. The factor $h_{\mathrm{SOCS1}}$ implements SOCS1-mediated negative feedback, so that increasing SOCS1 decreases the effective phosphorylation rate.

Phosphorylated STAT1 monomers $S1_{c_P}$ form cytosolic dimers $D_c$, which are transported to the nucleus. In the nuclear block ($N$), nuclear STAT1 dimers $D_n$ can bind DNA to form the transcriptionally active state $D_a$, unbind back to the nuclear dimer pool, undergo dephosphorylation, or be sequestered by PIAS1 into $D_i$. The variable $D_a$ is the central transcriptional readout of the JAK/STAT1 module and drives SOCS1, IRF1 and the downstream effector layer.

*p53-dependent modulation of JAK/STAT1 signalling*

The interaction between *p53* and JAK/STAT1 is represented by coarse-grained gates acting at different levels of the STAT1 response. The variable $P_s$ is a slow *p53*-dependent memory variable. It accumulates with *p53* activity and relaxes with rate $\delta_{\mathrm{mem}}$. Through $\Phi_{\mathrm{mem}}$, $P_s$ increases the effective DNA-binding rate $k_B^{\mathrm{eff}}$, decreases the effective unbinding rate $k_U^{\mathrm{eff}}$, and reduces the effective nuclear dephosphorylation rate $k_{\mathrm{deph}_n}^{\mathrm{eff}}$. Consequently, the same nuclear STAT1 input can generate a more persistent DNA-bound active state $D_a$ when prior *p53* activity has accumulated.

The synchronisation gate $\Phi_{\mathrm{sync}}$ represents acute co-activation between cytosolic STAT1 dimers and *p53*. It depends jointly on $D_c$ and $p53$, and increases the effective nuclear import term $\gamma_{D_c}^{\mathrm{eff}}$. The model therefore separates a slow *p53*-memory effect, mediated by $\Phi_{\mathrm{mem}}$, from a transient co-activation effect, mediated by $\Phi_{\mathrm{sync}}$.

Additional *p53*-dependent functions act downstream of the STAT1 core. The gate $\Phi_{p53}$ contributes to antiviral-effector amplification, whereas $\Phi_{\mathrm{apop}}$ represents *p53*-dependent apoptotic competence and potentiates the JAK/STAT–IRF1-associated apoptotic route. These functions do not modify receptor-level IFN input directly; they change how IFN-induced STAT1 activity is decoded at the transcriptional, effector and apoptotic-output levels.

*p53–Mdm2 stress response*

The *p53* module is driven by the viral-stress signal $S_{p53}$, which increases with viral genome burden $V_g$ and is scaled by the viral-to-stress gain $\chi_{p53}$. Keeping $\chi_{p53}$ independent from $\chi_{\mathrm{IFN}}$ allows the model to distinguish infection contexts with different balances between IFN induction and checkpoint-stress activation.

The $p53$ equation contains basal synthesis, stress-induced synthesis, *Mdm2*-dependent degradation and basal degradation. Stress increases $p53$ production through a high-order saturating function of $S_{p53}$, and also modulates the *Mdm2*-dependent degradation term through the factor involving $K_0$. The variables $Mdm2_{\mathrm{RNA}}$ and $Mdm2$ form the delayed negative-feedback loop: $p53$ induces $Mdm2_{\mathrm{RNA}}$, which produces $Mdm2$, and $Mdm2$ promotes $p53$ degradation. This architecture allows weak, oscillatory, damped or plateau-like *p53* regimes depending on the viral-stress input.

*STAT1–IRF1 memory and antiviral effector layer*

The effector layer translates nuclear STAT1 activity into antiviral and apoptotic outputs. The variables $A_D$, IRF1, $M_{\text{IRF1}}$ and $A_{\text{IRF1}}$ describe transcriptional and IRF1-associated memory downstream of $D_a$. Specifically, $A_D$ integrates DNA-bound STAT1 activity, IRF1 is induced by $D_a$, $M_{\text{IRF1}}$ integrates IRF1 activity over a slower timescale, and $A_{\text{IRF1}}$ captures delayed IRF1-associated activation.

The antiviral effectors $P_1$, $P_2$, and $P_3$ read different upstream signals. $P_1$ responds directly to $D_a$, $P_2$ responds to accumulated STAT1 activity $A_D$, and $P_3$ responds to IRF1 memory $M_{\text{IRF1}}$. Their production rates are multiplied by the *p53*-dependent amplification factors

$$A_{P_i} = (1 + \rho_{P_i}\Phi_{p53})\,(1 + \mu_{P_i}\Phi_{\text{mem}})\,, \qquad i \in \{1, 2, 3\}.$$

Thus, IFN-induced JAK/STAT1 activity remains the primary driver of the antiviral programme, while *p53* changes the effective gain and persistence of effector production.

The three antiviral effectors act on distinct steps of the viral life cycle: $P_1$ inhibits viral genome replication through $I_g$, $P_2$ inhibits viral-particle production through $I_p$, and $P_3$ enhances viral-particle loss through $I_{\text{loss}}$. This structure separates the antiviral programme into coarse-grained restriction axes and is consistent with the diversity of interferon-inducible mechanisms acting on viral replication, translation, intracellular permissiveness and infected-cell antiviral state (MacMicking, 2012; Schneider et al., 2014; Schoggins et al., 2011; Tretina et al., 2019).

*Apoptotic effectors and route-inclusive commitment*

The apoptotic layer separates two downstream routes. The variable $P_4$ represents a JAK/STAT–IRF1-associated apoptotic programme. Its production requires IRF1 memory $M_{\text{IRF1}}$, delayed IRF1-associated activation $A_{\text{IRF1}}$, and the viral-burden gate $\Phi_{V4}(V_g)$. The factor $(\lambda_{J4} + \lambda_{C4}\Phi_{\text{apop}})$ separates a basal JAK/STAT–IRF1-associated contribution from a *p53*-potentiated contribution. Since $\lambda_{V4} = 1$ in the retained calibration, $P_4$ remains explicitly conditional on persistent viral burden through $\Phi_{V4}(V_g)$.

The variable $P_5$ represents a *p53*-autonomous apoptotic programme. Its production depends directly on a high-threshold Hill function of $p53$, making it sensitive to sustained, high-amplitude *p53* activity and less responsive to weak or transient checkpoint activation. The biological interpretation of these two routes is consistent with the known involvement of IFN/STAT1/IRF1-linked mechanisms in apoptosis and with canonical *p53*-regulated apoptotic programmes (Fulda and Debatin, 2002; Tamura et al., 1995; Gao et al., 2010; Michalak et al., 2005).

Apoptotic commitment is route-inclusive and duration-dependent. A brief threshold crossing is not sufficient. Route $i \in \{4, 5\}$ is considered committed only if $P_i$ remains above its threshold $\theta_{P_i}$ for at least a route-specific duration $\tau_i$:

$$\mathcal{S}_i = 1 \quad \Longleftrightarrow \quad \exists t \text{ such that } P_i(s) > \theta_{P_i} \text{ for all } s \in [t - \tau_i, t].$$

The apoptotic decision is then

$$\mathcal{A}_{\text{apop}} = 1 \quad \Longleftrightarrow \quad \mathcal{S}_4 = 1 \text{ or } \mathcal{S}_5 = 1.$$

The route is classified as JAK/STAT-associated when only $P_4$ satisfies the criterion, *p53*-autonomous when only $P_5$ satisfies it, and dual-route when both criteria are satisfied.

*Viral dynamics and antiviral feedback*

The viral module tracks intracellular viral genomes $V_g$ and viral particles $V_p$. Viral genomes grow logistically with intrinsic replication rate $r_g$ and carrying capacity $K_g$, while viral particles are produced from viral genomes at rate $k_p^V$. The antiviral effectors feed back onto the viral module through three inhibition or loss terms:

$$I_g = \frac{1}{1 + \lambda_g\sigma_g P_1}, \qquad I_p = \frac{1}{1 + \lambda_m\sigma_m P_2}, \qquad I_{\text{loss}} = 1 + \lambda_a\sigma_a \frac{P_3}{P_3 + K_{P3v}}.$$

The sensitivity parameters $\sigma_g$, $\sigma_m$, and $\sigma_a$ define how vulnerable a viral class is to effector-mediated control. A viral class may strongly induce IFN or checkpoint stress, yet remain poorly controlled if its replication or particle-production steps are weakly sensitive to $P_1$, $P_2$, or $P_3$.

The viral module closes the host–virus feedback loop. Viral genomes induce $IFN_{\text{int}}$ and $S_{p53}$, while IFN-, STAT1-, IRF1- and *p53*-dependent effectors restrict viral genome replication, viral-particle production and viral-particle persistence. The model therefore separates five viral traits: viral-to-IFN induction ($\chi_{\text{IFN}}$), viral-to-checkpoint-stress induction ($\chi_{p53}$), sensitivity to genome restriction ($\sigma_g$), sensitivity to particle-production inhibition ($\sigma_m$), and sensitivity to particle loss ($\sigma_a$).

*Interpretation of the model architecture*

The model separates three levels of outcome: antiviral-state engagement, functional viral control and apoptotic commitment. The variables $P_1$, $P_2$, and $P_3$ define the antiviral state of the host programme; late viral burden, measured through $V_g$ and $V_p$, defines the functional viral outcome; and $P_4$ and $P_5$ define route-specific apoptotic commitment.

These outcomes need not coincide. A strong antiviral state may fail to suppress a poorly sensitive viral class, whereas apoptosis may be triggered through the JAK/STAT–IRF1-associated route, the *p53*-autonomous route, or both. The model therefore provides a framework for analysing how viral burden is translated into IFN signalling, checkpoint activation, antiviral restriction and apoptotic routing.

### S2. Mathematical properties

We explain here the model structure and give the mathematical properties of the model described in Equations (2.1)–(2.6).

#### *S2.1. Mathematical properties of the coupled model*

This supplementary section establishes basic mathematical properties of the coupled IFN–JAK/STAT–*p53*–effector–virus model used in the main text. The aim is not to re-interpret the biology of the model, which is done in the main manuscript, but to verify that the system defines a well-posed dynamical model on the biologically admissible state space.

*Notation and state space*

Let $n \in \mathbb{N}$ denote the total number of state variables of the coupled system given in Equation **??**. We collect the state variables into the vector

$$\begin{aligned}\mathbf{x}(t) = \big(&R_f, R_a, R_d, R_i,\ S1_c, S1_{cP}, D_c, C_P,\ \mathrm{soc1}, \mathrm{SOCS1}, PTP_c,\ S1_n, D_n, D_a, D_i, PTP_n, PIAS1_n,\ P_s,\\ &S_{p53}, p53, \mathrm{Mdm2}_{\mathrm{RNA}}, \mathrm{Mdm2},\ S_v, E_v, \mathrm{IFN}_{int},\ A_D, \mathrm{IRF1}, M_{\mathrm{IRF1}}, A_{\mathrm{IRF1}},\\ &P_1, P_2, P_3, P_4, P_5,\ V_g, V_p\big)^\top \in \mathbb{R}^n.\end{aligned}$$

The system can be written abstractly as

$$\dot{\mathbf{x}}(t) = F\big(\mathbf{x}(t), t\big), \qquad \mathbf{x}(0) = \mathbf{x}_0, \tag{S2.1}$$

where $F$ collects the right-hand sides of the differential equations and the auxiliary functions defined in the main model.

The biologically admissible state space is the non-negative orthant

$$\mathbb{R}^n_+ = \big\{\mathbf{x} \in \mathbb{R}^n : x_i \geq 0,\ i = 1, \ldots, n\big\}.$$

All variables represent molecular concentrations, activities, memory variables, or viral loads, and therefore negative values are biologically inadmissible.

*Local existence and uniqueness*

**Theorem 1** (Local existence and uniqueness)**.** *For every initial condition $\mathbf{x}_0 \in \mathbb{R}^n_+$, the initial-value problem* (S2.1) *has a unique maximal solution*

$$\mathbf{x} \in C^1\big([0, T_{\max}), \mathbb{R}^n\big)$$

*for some $T_{\max} \in (0, +\infty]$.*

*Proof.* The vector field $F$ is locally Lipschitz on $\mathbb{R}^n_+$. Indeed, each component of $F$ is a finite combination of polynomial terms, linear decay terms, Hill-type activation functions, and Michaelis–Menten-type saturating terms.

All Hill-type terms used in the model have the form

$$H(\xi; K, m) = \frac{\xi^m}{\xi^m + K^m}, \qquad \xi \geq 0, \quad K > 0, \quad m \in \mathbb{N}.$$

Since the denominator satisfies $\xi^m + K^m \geq K^m > 0$, such functions are smooth on $\mathbb{R}_+$. Similarly, Michaelis–Menten terms such as

$$\frac{p53}{p53 + K_u}$$

are smooth on $\mathbb{R}_+$ because $K_u > 0$. Hence every component of $F$ is locally Lipschitz on $\mathbb{R}^n_+$. The result follows from the standard Picard–Lindelöf theorem for ordinary differential equations. □

| Name | Meaning | Units | Value |
|---|---|---|---|
| *p53 core dynamics* | | | |
| $\beta_0$ | Basal *p53* synthesis rate. | $a.u.\,h^{-1}$ | 0.0005 |
| $\beta_{p53}$ | Stress-induced *p53* synthesis rate. | $a.u.\,h^{-1}$ | 60.0 |
| $K_S$ | Stress threshold for *p53* induction. | $a.u.$ | 6.0 |
| $n_\beta$ | Hill coefficient for stress-induced *p53* synthesis. | – | 4.0 |
| $\delta_p$ | Basal *p53* degradation rate. | $h^{-1}$ | 0.5 |
| $k_u$ | Maximal *Mdm2*-dependent *p53* degradation rate. | $h^{-1}$ | 30.0 |
| $K_u$ | Michaelis constant for *p53* degradation. | $a.u.$ | 0.15 |
| $K_0$ | Stress threshold controlling *Mdm2*-dependent *p53* degradation. | $a.u.$ | 12.0 |
| $n_p$ | Hill coefficient in the stress-dependent modulation of *Mdm2*-dependent *p53* degradation. | – | 6.0 |
| *Mdm2 feedback loop* | | | |
| $k_{tr}$ | Effective *Mdm2* mRNA transcription rate. | $a.u.\,h^{-1}$ | 0.00964 |
| $\delta_m$ | *Mdm2* mRNA degradation rate. | $h^{-1}$ | 0.8 |
| $k_{tl}$ | *Mdm2* protein translation rate. | $h^{-1}$ | 0.6 |
| $\delta_M$ | *Mdm2* protein degradation rate. | $h^{-1}$ | 0.5 |
| *Viral-burden-to-p53 stress conversion* | | | |
| $\chi_{p53}$ | Viral-burden-to-*p53* stress induction gain. | – | 0.20 / 0.80 / 1.20 |
| $S_{\max,p53}$ | Maximal viral-induced checkpoint-stress input. | $a.u.$ | 25.0 |
| $K_{V,p53}$ | Viral-load threshold for viral-induced checkpoint stress. | $a.u.$ | 0.50 |
| $n_{V,p53}$ | Hill coefficient for viral-burden-to-stress conversion. | – | 2.0 |
| $k_{\text{relax},S_{p53}}$ | Relaxation rate of the dynamic checkpoint-stress signal $S_{p53}$. | $h^{-1}$ | 0.25 |
| *p53 memory and JAK/STAT coupling* | | | |
| $k_{\text{mem}}$ | Accumulation rate of the *p53* memory variable $P_s$. | $h^{-1}$ | 0.10 |
| $\delta_{\text{mem}}$ | Relaxation rate of *p53* memory. | $h^{-1}$ | 0.04 |
| $K_{\text{mem}}$ | Threshold in $\Phi_{\text{mem}}$. | $a.u.$ | 8.0 |
| $K_{\text{sync}}$ | STAT1 threshold in $\Phi_{\text{sync}}$. | $a.u.$ | 0.08 |
| $K_{\text{psyn}}$ | *p53* threshold in $\Phi_{\text{sync}}$. | $a.u.$ | 8.0 |
| $K_{\text{apop}}$ | *p53* threshold in $\Phi_{\text{apop}}$. | $a.u.$ | 10.0 |
| $K_{p53}$ | *p53* threshold in $\Phi_{p53}$. | $a.u.$ | 14.0 |
| $\rho_{B_m}$ | Memory-dependent enhancement of $k_B^{\text{eff}}$. | – | 0.8 |
| $\rho_{B_s}$ | Synchronisation-dependent enhancement of $k_B^{\text{eff}}$. | – | 0.5 |
| $\eta_U$ | Memory-dependent reduction of $k_U^{\text{eff}}$. | – | 0.4 |
| $\eta_{\text{deph}}$ | Memory-dependent reduction of $k_{\text{deph}_n}^{\text{eff}}$. | – | 0.5 |
| $\rho_{D_c}$ | Synchronisation-dependent enhancement of $\gamma_{D_c}^{\text{eff}}$. | – | 0.6 |
| *External and endogenous IFN protocol* | | | |
| $IFN_0$ | External IFN pulse amplitude. | $a.u.$ | 0.0 |
| $IFN_{\text{start}}$ | Starting time of the external IFN pulse. | $h$ | 0.0 |
| $T_{\text{IFN}}$ | Duration of the external IFN pulse when an external IFN source is used. | h | 120.0 |
| $\tau_{\text{dec}}$ | Decay time constant after IFN removal. | $h$ | 8.0 |
| $\eta_{\text{ext}}$ | Weight of the external IFN contribution. | – | 0.0 |
| $\eta_{\text{int}}$ | Weight of the endogenous IFN contribution. | – | 1.0 |
| $IFN_{\text{sat}}$ | Saturation level for total IFN. | $a.u.$ | 60.0 |
| *Endogenous IFN sensing* | | | |
| $\chi_{\text{IFN}}$ | Viral-burden-to-IFN induction gain. | – | 0.022927 / 0.045369 / 0.10032 |
| $k_{\text{sens}}$ | Production rate of the sensing variable $S_v$. | $h^{-1}$ | 0.45 |
| $\delta_{\text{sens}}$ | Decay rate of $S_v$. | $h^{-1}$ | 0.10 |
| $K_{V,\text{sens}}$ | Viral-load threshold for endogenous IFN sensing. | $a.u.$ | 0.12 |
| $\alpha_S$ | Positive-feedback strength in $S_v$ amplification. | – | 0.35 |
| $K_{\text{exh}}$ | Exhaustion threshold limiting sensing amplification. | $a.u.$ | 1.0 |
| $k_E$ | Accumulation rate of exhaustion variable $E_v$. | $h^{-1}$ | 0.020 |
| $\delta_E$ | Decay rate of $E_v$. | $h^{-1}$ | 0.015 |
| $k_{\text{IFN}}$ | Endogenous IFN production rate from $S_v$. | $a.u.\,h^{-1}$ | 7.2 |
| $\delta_{\text{IFN}}$ | Decay rate of endogenous IFN. | $h^{-1}$ | 0.12 |
| $K_{S,\text{IFN}}$ | Threshold for IFN production from $S_v$. | $a.u.$ | 0.20 |

Table S1: Parameters of the *p53–Mdm2* module, the viral-burden-to-*p53* stress conversion, the *p53*-dependent JAK/STAT coupling layer, and the IFN sensing module. For scenario-dependent *p53* stress induction, values of $\chi_{p53}$ are reported for the low, intermediate, and high viral-to-*p53* stress classes, respectively. For scenario-dependent IFN induction, values of $\chi_{\text{IFN}}$ are reported for the low, intermediate, and high virus-induced IFN classes, respectively, using the calibration retained in the latest virus-induced IFN/*p53* simulations.

| Name | Meaning | Units | Value |
|---|---|---|---|
| *STAT1–IRF1 memory layer* | | | |
| $k_{A_D}$ | Accumulation rate of STAT1 DNA-binding memory $A_D$. | $h^{-1}$ | 0.35 |
| $\delta_{A_D}$ | Decay rate of $A_D$. | $h^{-1}$ | 0.05 |
| $\beta_I$ | Maximal IRF1 induction rate. | $a.u.\, h^{-1}$ | 1.0 |
| $K_I$ | STAT1 activity threshold for IRF1 induction. | $a.u.$ | 0.05 |
| $\delta_I$ | IRF1 degradation rate. | $h^{-1}$ | 0.12 |
| $k_{M_I}$ | Accumulation rate of IRF1 memory. | $h^{-1}$ | 0.20 |
| $\delta_{M_I}$ | Decay rate of IRF1 memory. | $h^{-1}$ | 0.05 |
| $k_{A_4}$ | Accumulation rate of delayed IRF1-associated apoptotic activation. | $h^{-1}$ | 0.05 |
| $\delta_{A_4}$ | Decay rate of $A_{\mathrm{IRF1}}$. | $h^{-1}$ | 0.01 |
| *Antiviral effectors $P_1$–$P_3$* | | | |
| $\beta_{P_1}$ | Maximal production rate of $P_1$. | $a.u.\, h^{-1}$ | 1.20 |
| $\beta_{P_2}$ | Maximal production rate of $P_2$. | $a.u.\, h^{-1}$ | 1.00 |
| $\beta_{P_3}$ | Maximal production rate of $P_3$. | $a.u.\, h^{-1}$ | 0.80 |
| $K_{P1}$ | Activation threshold for $P_1$. | $a.u.$ | 0.05 |
| $K_{P2}$ | Activation threshold for $P_2$. | $a.u.$ | 2.0 |
| $K_{P3}$ | Activation threshold for $P_3$. | $a.u.$ | 2.0 |
| $\delta_{P1}$ | Degradation rate of $P_1$. | $h^{-1}$ | 0.18 |
| $\delta_{P2}$ | Degradation rate of $P_2$. | $h^{-1}$ | 0.10 |
| $\delta_{P3}$ | Degradation rate of $P_3$. | $h^{-1}$ | 0.07 |
| $\rho_{P_1}$ | *p53*-dependent amplification strength of $P_1$. | – | 0.10 |
| $\rho_{P_2}$ | *p53*-dependent amplification strength of $P_2$. | – | 0.30 |
| $\rho_{P_3}$ | *p53*-dependent amplification strength of $P_3$. | – | 0.60 |
| $\mu_{P_1}$ | *p53*-memory-dependent amplification strength of $P_1$. | – | 0.01 |
| $\mu_{P_2}$ | *p53*-memory-dependent amplification strength of $P_2$. | – | 0.13 |
| $\mu_{P_3}$ | *p53*-memory-dependent amplification strength of $P_3$. | – | 0.15 |
| *Apoptotic effectors $P_4$ and $P_5$* | | | |
| $\beta_{P_4}$ | Maximal production rate of JAK/STAT–IRF1 apoptotic effector $P_4$. | $a.u.\, h^{-1}$ | 14.6 |
| $\delta_{P4}$ | Degradation rate of $P_4$. | $h^{-1}$ | 0.08 |
| $K_{I4}$ | IRF1-memory threshold for $P_4$ activation. | $a.u.$ | 3.50 |
| $K_{A4}$ | Delayed IRF1-activation threshold for $P_4$. | $a.u.$ | 5.50 |
| $\lambda_{J4}$ | JAK/STAT–IRF1 basal contribution to $P_4$. | – | 0.35 |
| $\lambda_{C4}$ | *p53*-dependent contribution to $P_4$ through $\Phi_{\mathrm{apop}}$. | – | 0.65 |
| $\lambda_{V4}$ | Viral-burden gate weight in $P_4$ activation. | – | 1.00 |
| $K_{V4}$ | Viral-burden threshold in the $P_4$ gate $\Phi_{V4}$. | $a.u.$ | 0.25 |
| $n_{V4}$ | Hill coefficient of the viral-burden gate $\Phi_{V4}$. | – | 2.00 |
| $\beta_{P_5}$ | Maximal production rate of *p53*-autonomous effector $P_5$. | $a.u.\, h^{-1}$ | 4.0 |
| $\delta_{P5}$ | Degradation rate of $P_5$. | $h^{-1}$ | 0.08 |
| $K_{P5}$ | *p53* activation threshold for $P_5$ induction. | $a.u.$ | 5.50 |
| $n_{P5}$ | Hill coefficient for *p53*-dependent induction of $P_5$. | – | 4.0 |
| *Viral replication and protein dynamics* | | | |
| $r_g$ | Intrinsic viral genome replication rate. | $h^{-1}$ | 0.10 |
| $K_g$ | Viral genome carrying capacity. | $a.u.$ | 1.0 |
| $\delta_g^V$ | Viral genome degradation or clearance rate. | $h^{-1}$ | 0.03 |
| $k_p^V$ | Viral protein production rate from viral genomes. | $h^{-1}$ | 0.50 |
| $\delta_p^V$ | Basal viral protein loss rate. | $h^{-1}$ | 0.35 |
| *Antiviral sensitivity parameters* | | | |
| $\lambda_g$ | Strength of $P_1$-mediated inhibition of viral genome replication. | – | 8.0 |
| $\lambda_m$ | Strength of $P_2$-mediated inhibition of viral protein production. | – | 5.0 |
| $\lambda_a$ | Strength of $P_3$-mediated enhancement of viral protein loss. | – | 3.0 |
| $K_{P3v}$ | Half-saturation threshold for the $P_3$ effect on viral protein loss. | $a.u.$ | 15.0 |
| $\sigma_g$ | Viral sensitivity to genome-replication inhibition. | – | 8.5 / 0.05 / 0.0002 |
| $\sigma_m$ | Viral sensitivity to viral-protein-production inhibition. | – | 1.20 / 0.00208 / 0.0004 |
| $\sigma_a$ | Viral sensitivity to enhanced viral protein loss. | – | 1.70 / 0.00417 / 0.0003 |

Table S2: Parameters of the STAT1–IRF1 memory layer, antiviral effectors, apoptotic effectors, and viral dynamics. For viral sensitivity parameters, values are reported for the sensitive, intermediate, and highly resistant viral classes, respectively, using the calibration retained in the latest virus-induced IFN/*p53* simulations. The apoptotic calibration includes the viral-burden gate in $P_4$, through $\lambda_{V4}$, $K_{V4}$, and $n_{V4}$.

*Positivity of solutions*

**Theorem 2** (Positivity)**.** *If* $\mathbf{x}_0 \in \mathbb{R}^n_+$*, then the maximal solution of* (S2.1) *satisfies*

$$\mathbf{x}(t) \in \mathbb{R}^n_+ \qquad \textit{for all } t \in [0, T_{\max}).$$

*Proof.* We use the standard quasi-positivity criterion. It is sufficient to verify that, for every component $x_i$, the corresponding right-hand side satisfies

$$F_i(\mathbf{x}, t) \geq 0 \qquad \text{whenever } x_i = 0 \text{ and } \mathbf{x} \in \mathbb{R}^n_+.$$

This condition ensures that the vector field does not point outside the non-negative orthant on its boundary.

For the receptor variables, setting any receptor state to zero leaves only non-negative incoming fluxes. For example,

$$\dot{R}_f\big|_{R_f=0} = k_{off} R_a + k_{reac} R_d + k_{rec} R_i \geq 0.$$

The same cancellation structure applies to $R_a$, $R_d$, and $R_i$.

For the cytosolic STAT1 variables, setting $S1_c = 0$ gives

$$\dot{S}1_c = \beta_{S1} + \gamma_{S1n} S1_n + k_{deph_c} S1_{cP} + k_{dph} C_P \geq 0.$$

Similarly,

$$\dot{S}1_{cP}\big|_{S1_{cP}=0} = v_{phos} R_a S1_c + k_{ubd} C_P \geq 0,$$

and the equations for $D_c$ and $C_P$ contain only non-negative production terms when the corresponding variable is zero.

For the nuclear STAT1 variables, all production terms are non-negative at the boundary. For instance,

$$\dot{D}_a\big|_{D_a=0} = k_B^{eff} D_n \geq 0, \qquad \dot{D}_i\big|_{D_i=0} = k_{su} PIAS1_n D_n \geq 0.$$

The same argument applies to $S1_n$, $D_n$, $PTP_n$, $PIAS1_n$, and $P_s$.

For the *p53* module,

$$\dot{S}_{p53}\big|_{S_{p53}=0} = k_{S_{p53}} \chi_{p53} \frac{V_g^2}{V_g^2 + K_{V,p53}^2} \geq 0,$$

and

$$\dot{p53}\big|_{p53=0} = \beta_0 + \beta_{p53} \frac{S_{p53}^{n_\beta}}{K_S^{n_\beta} + S_{p53}^{n_\beta}} \geq \beta_0 > 0.$$

Moreover,

$$\dot{\text{Mdm2}}_{\text{RNA}}\big|_{\text{Mdm2}_{RNA}=0} = k_{tr} p53^{\,2} \geq 0, \qquad \dot{\text{Mdm2}}\big|_{\text{Mdm2}=0} = k_{tl} \text{Mdm2}_{RNA} \geq 0.$$

For the sensing, memory, effector, and viral protein variables, the equations all have the generic structure

$$\dot{z} = \alpha(\mathbf{x}, t) - \delta_z z, \qquad \alpha(\mathbf{x}, t) \geq 0.$$

Thus, at $z = 0$, one has $\dot{z} = \alpha(\mathbf{x}, t) \geq 0$. For the viral genome variable,

$$\dot{V}_g\big|_{V_g=0} = 0,$$

which is also compatible with quasi-positivity.

Therefore the vector field is quasi-positive on $\mathbb{R}^n_+$, and the non-negative orthant is forward invariant. □

*Boundedness and global existence*

**Lemma 1** (Receptor conservation)**.** *The total receptor pool is conserved:*

$$R_f(t) + R_a(t) + R_d(t) + R_i(t) = R_{\text{tot}},$$

*where*

$$R_{\text{tot}} = R_f(0) + R_a(0) + R_d(0) + R_i(0).$$

*Proof.* Summing the four receptor equations gives

$$\frac{d}{dt}(R_f + R_a + R_d + R_i) = 0,$$

because all receptor transitions are internal transfers between the four receptor states. Hence the total receptor pool is conserved. □

**Theorem 3** (Boundedness and global existence)**.** *Every solution of* (S2.1) *with initial condition* $\mathbf{x}_0 \in \mathbb{R}^n_+$ *is bounded on finite and infinite time intervals. Consequently,* $T_{\max} = +\infty$.

*Proof.* By Lemma 1 and Theorem 2,

$$0 \le R_f, R_a, R_d, R_i \le R_{\text{tot}}.$$

The viral genome equation contains logistic growth and non-negative antiviral inhibition. Since $0 < I_g \le 1$, one has

$$\dot{V}_g \le r_g V_g \left(1 - \frac{V_g}{K_g}\right),$$

up to the additional non-positive clearance term. Hence

$$V_g(t) \le \max\{V_g(0), K_g\}.$$

It follows that the viral-stress input satisfies

$$\dot{S}_{p53} \le k_{S_{p53}} \chi_{p53} - \delta_{S_{p53}} S_{p53},$$

and therefore $S_{p53}$ is bounded by comparison.

Since the stress-induced *p53* synthesis term is bounded by $\beta_{p53}$ and the *Mdm2*-dependent degradation term is non-negative,

$$\dot{p53} \le \beta_0 + \beta_{p53} - \delta_p p53.$$

Thus,

$$p53(t) \le \max\left\{p53(0), \frac{\beta_0 + \beta_{p53}}{\delta_p}\right\} := \overline{p53}.$$

The equations for $\text{Mdm2}_{RNA}$ and Mdm2 are then bounded by linear comparison:

$$\dot{\text{Mdm2}}_{RNA} \le k_{tr} \overline{p53}^{\,2} - \delta_m \text{Mdm2}_{RNA},$$

and

$$\dot{\text{Mdm2}} \le k_{tl} \overline{\text{Mdm2}}_{RNA} - \delta_M \text{Mdm2}.$$

Hence both variables are bounded. The same comparison argument gives a bound for $P_s$:

$$\dot{P}_s \le k_{mem} \overline{p53} - \delta_{mem} P_s.$$

The gates $\Phi_{mem}$, $\Phi_{sync}$, $\Phi_{apop}$, and $\Phi_{p53}$ are bounded between 0 and 1. Therefore all effective rates defined from these gates are bounded above and remain non-negative, provided the parameter conditions stated in Proposition 1 hold. In particular,

$$k_B^{eff} \le k_B(1 + \rho_{B_m})(1 + \rho_{B_s}), \qquad \gamma_{D_c}^{eff} \le \gamma_{D_c,IFN}(1 + \rho_{D_c}).$$

Using these bounds, together with the bounded receptor variables and bounded input functions, the STAT1, SOCS1, phosphatase, and PIAS1 equations are bounded by systems with at most linear production and strictly negative first-order loss terms. Standard comparison arguments therefore give upper bounds for

$$S1_c,\ S1_{cP},\ D_c,\ C_P,\ \text{soc1},\ \text{SOCS1},\ PTP_c, \quad S1_n,\ D_n,\ D_a,\ D_i,\ PTP_n,\ PIAS1_n.$$

The sensing variables $S_v$, $E_v$, and $\text{IFN}_{int}$ are also bounded because their production terms are saturating and their loss terms are first-order. The memory and effector variables

$$A_D,\ \text{IRF1},\ M_{\text{IRF1}},\ A_{\text{IRF1}}, \quad P_1, \ldots, P_5$$

have bounded production terms and first-order degradation. Hence each is bounded by a scalar comparison equation of the form

$$\dot{z} \le \beta_z - \delta_z z.$$

Finally,

$$\dot{V}_p \le k_p^V \overline{V_g} - \delta_p^V V_p,$$

because $I_p \le 1$ and $I_{loss} \ge 1$. Thus $V_p$ is bounded.

All components of $\mathbf{x}(t)$ are therefore bounded on $[0, T_{\max})$. Since finite-time blow-up is the only obstruction to extending a solution of a locally Lipschitz ODE, the maximal existence time is $T_{\max} = +\infty$. □

*Forward-invariant admissible set*

**Corollary 1** (Global well-posedness on a bounded admissible set)**.** *There exists a compact set*

$$\Omega = \left\{ \mathbf{x} \in \mathbb{R}_+^n : 0 \le x_i \le \bar{x}_i,\ i = 1, \ldots, n \right\},$$

*where each $\bar{x}_i$ is an upper bound obtained in Theorem 3, such that every solution starting in $\Omega$ remains in $\Omega$ for all $t \ge 0$.*

*Proof.* The lower boundary is invariant by Theorem 2. The upper bounds $\bar{x}_i$ are chosen from comparison inequalities of the form

$$\dot{x}_i \le \beta_i - \delta_i x_i,$$

or from conservation/logistic bounds. Hence the vector field points inward at the corresponding upper boundaries. Therefore $\Omega$ is positively invariant. Compactness follows directly from its definition as a closed and bounded subset of $\mathbb{R}^n$. □

*Mathematical consistency of the p53-dependent modulation*

**Proposition 1** (Preservation of well-posedness under *p53* coupling)**.** *The p53-dependent modulations introduced through $\Phi_{mem}$ and $\Phi_{sync}$ preserve local existence, uniqueness, positivity, boundedness, and global existence, provided*

$$0 \le \eta_U < 1, \qquad 0 \le \eta_{deph} < 1.$$

*Proof.* The gates $\Phi_{mem}$ and $\Phi_{sync}$ are Hill-type functions:

$$\Phi_{mem} = \frac{P_s^2}{P_s^2 + K_{mem}^2},$$

and

$$\Phi_{sync} = \frac{D_c^2}{D_c^2 + K_{sync}^2} \cdot \frac{p53^3}{p53^3 + K_{psyn}^3}.$$

Both are smooth on $\mathbb{R}_+^n$ and satisfy

$$0 \le \Phi_{mem} < 1, \qquad 0 \le \Phi_{sync} < 1.$$

Consequently, the effective rates satisfy

$$k_B^{eff} = k_B(1 + \rho_{B_m}\Phi_{mem})(1 + \rho_{B_s}\Phi_{sync})$$

and therefore

$$k_B \le k_B^{eff} \le k_B(1 + \rho_{B_m})(1 + \rho_{B_s}).$$

Similarly,

$$k_U^{eff} = k_U(1 - \eta_U \Phi_{mem})$$

satisfies

$$k_U(1 - \eta_U) \le k_U^{eff} \le k_U.$$

The assumption $\eta_U < 1$ ensures that $k_U^{eff}$ remains strictly positive.

For the nuclear dephosphorylation rate,

$$k_{deph_n}^{eff} = \left( k_{deph_n 0} + k_{P_n} \frac{PTP_n}{PTP_n + K_{PTP_n}} \right)(1 - \eta_{deph}\Phi_{mem}),$$

and since

$$0 \leq \frac{PTP_n}{PTP_n + K_{PTP_n}} \leq 1,$$

one obtains a bounded non-negative rate. The condition $\eta_{deph} < 1$ prevents complete suppression of nuclear dephosphorylation. Finally,

$$\gamma_{D_c}^{eff} = \gamma_{D_c,IFN}(1 + \rho_{D_c}\Phi_{sync})$$

is bounded between

$$\gamma_{D_c,IFN} \quad \text{and} \quad \gamma_{D_c,IFN}(1 + \rho_{D_c}).$$

Thus, the *p53*-dependent coupling only introduces bounded, state-dependent multiplicative modulations of existing kinetic rates. It does not introduce singularities, negative rates, or unbounded source terms. Therefore the local Lipschitz, quasi-positivity, and boundedness arguments used above remain valid. □

**Remark 1.** *The conditions $\eta_U < 1$ and $\eta_{deph} < 1$ have a simple mathematical and biological interpretation. They ensure that DNA-bound STAT1 retains a finite unbinding rate and that nuclear STAT1 dephosphorylation is reduced but not abolished. In the simulations used in the main text, these conditions are satisfied by the parameter values reported in the supplementary parameter tables.*

*Apoptotic commitment as a well-defined event*

The main text treats apoptotic commitment as a route-inclusive decision: commitment can be triggered by the JAK/STAT–IRF1-associated effector $P_4$ or by the *p53*-autonomous effector $P_5$. In the supplementary formulation, we also impose a persistence condition so that a transient threshold crossing is not sufficient.

For each route $i \in \{4, 5\}$, let $\theta_{P_i} > 0$ denote the corresponding threshold and let $\tau_i > 0$ denote the minimum time during which the effector must remain above threshold. The apoptotic commitment indicator is defined by

$$A_{apop}(t) = 1 \quad \Longleftrightarrow \quad \exists i \in \{4, 5\} \text{ such that } P_i(s) > \theta_{P_i} \quad \forall s \in [t - \tau_i, t].$$

Equivalently, the first commitment time is

$$t^* = \inf \{t \geq 0 : \exists i \in \{4, 5\} \text{ such that } P_i(s) > \theta_{P_i} \quad \forall s \in [t - \tau_i, t]\},$$

with the convention $t^* = +\infty$ if the condition is never satisfied.

**Proposition 2** (Well-defined apoptotic commitment rule)**.** *The duration-dependent apoptotic commitment rule is well defined along every solution of the coupled system.*

*Proof.* By Theorem 3, the solution exists globally and all components of $\mathbf{x}(t)$ are continuous functions of time. In particular, $P_4(t)$ and $P_5(t)$ are continuous on $[0, +\infty)$.

For each $i \in \{4, 5\}$, the set

$$\mathcal{A}_i = \{t \geq 0 : P_i(t) > \theta_{P_i}\}$$

is open because $P_i$ is continuous. The condition

$$P_i(s) > \theta_{P_i} \quad \forall s \in [t - \tau_i, t]$$

is therefore a well-defined persistence condition on a compact time interval. Thus, the indicator $A_{apop}(t)$ is well defined for all $t \geq 0$, and the first commitment time $t^*$ is a well-defined extended real number. □

**Remark 2.** *The use of route-specific durations $\tau_4$ and $\tau_5$ avoids imposing the same commitment time on the two apoptotic routes. This is consistent with the structure of the model: $P_4$ represents a JAK/STAT–IRF1-associated apoptotic route, whereas $P_5$ represents a p53-autonomous route. The two timescales may be set equal in a specific calibration, but the mathematical definition does not require this.*

### S3. Definition of response metrics

To compare dynamical responses across stimulation protocols, we use a set of scalar metrics that summarise complementary features of a time-dependent model output. These metrics are not restricted to a single molecular species. Instead, they can be applied to any state variable or derived readout analysed in the manuscript, including active STAT1 species, memory variables, transcriptional effectors, apoptotic outputs, or viral variables.

For clarity, we introduce the definitions for a generic observable

$$Y(t) = Y\big(t; \mathcal{P}, p\big), \qquad t \in [0, T_{\mathrm{obs}}],$$

where $\mathcal{P}$ denotes the stimulation protocol, $p$ denotes the model parameter set, and $T_{\mathrm{obs}}$ is the observation time. In the main text, the choice of $Y(t)$ depends on the biological question addressed. For instance, when analysing JAK/STAT1 activation, we may take

$$Y(t) = D_a(t),$$

where $D_a(t)$ denotes the transcriptionally active DNA-bound STAT1 dimer. Alternatively, when the objective is to quantify the total pool of activated nuclear STAT1, one may use

$$Y(t) = D_n(t) + D_a(t).$$

The same definitions can also be applied to downstream quantities such as $A_D(t)$, $M_{\mathrm{IRF1}}(t)$, $P_i(t)$, or viral readouts such as $V_g(t)$ and $V_p(t)$, provided that the biological interpretation of the metric is adapted to the chosen observable.

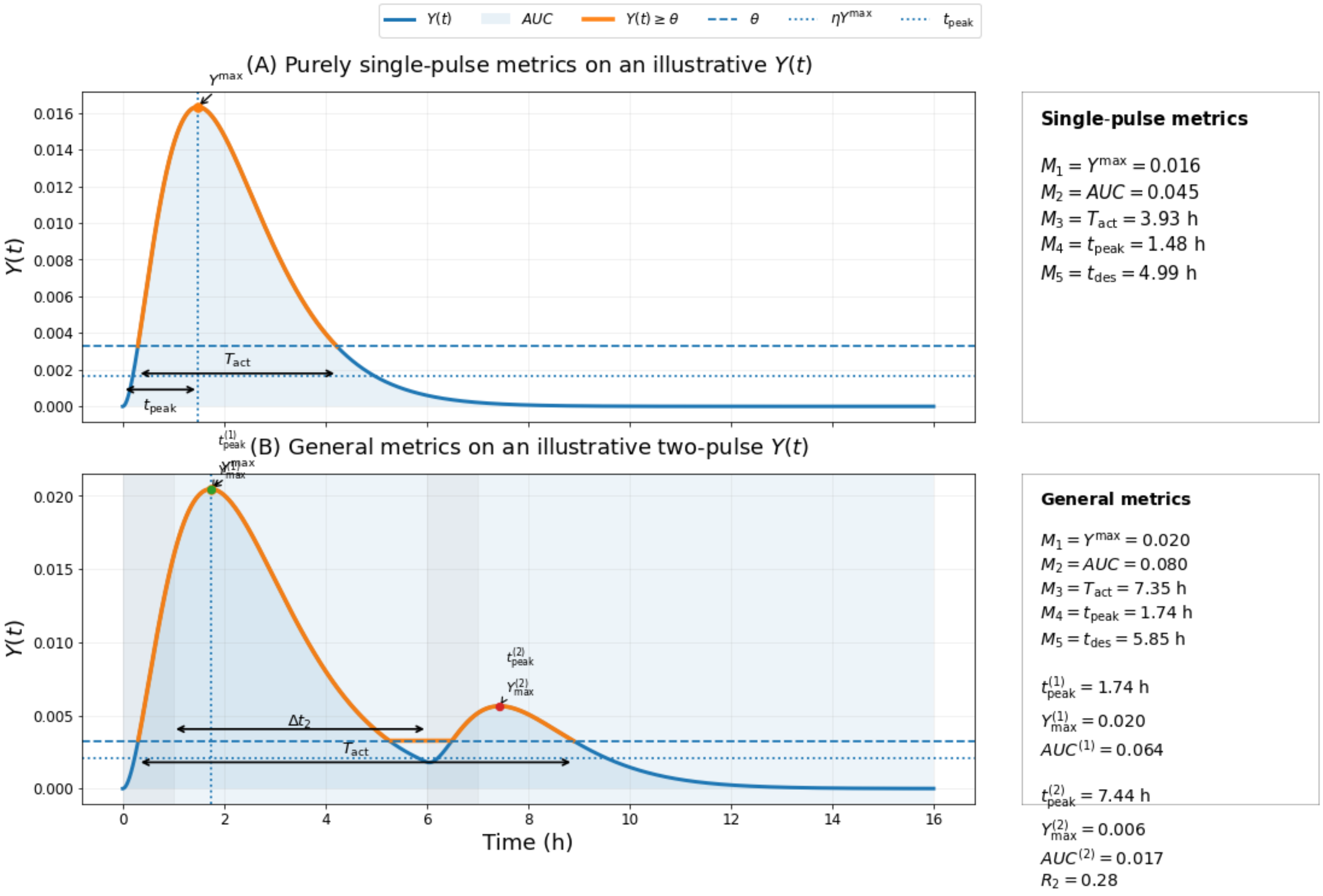


Figure S1: Illustrative STAT1 activity metrics defined on a synthetic nuclear signal $Y(t)$. Panel (A) shows the basic single-pulse metrics $M_1$–$M_5$: peak amplitude, integrated activity, time above threshold, time-to-peak, and post-peak desensitisation time. Panel (B) shows their extension to a two-pulse protocol, together with pulse-resolved quantities such as $t^{(j)}_{\mathrm{peak}}$, $D^{(j)}_{n,\max}$, $AUC^{(j)}$, the interpulse interval $\Delta t_2$, and the recovery ratio $R_2$. The displayed trajectory is purely illustrative.

*S3.1. Global protocol-level metrics*

We first define metrics over the full observation window $[0, T_{\text{obs}}]$. These quantities are valid for arbitrary stimulation protocols, including single-pulse, delayed-pulse, repeated-pulse, and combined-input protocols.

*Peak response.* The maximal value of the observable over the observation window is defined as

$$M_1(\mathcal{P}) := Y_{\max}(\mathcal{P}) = \max_{0 \le t \le T_{\text{obs}}} Y(t). \tag{S3.1}$$

This metric captures the strongest instantaneous response induced by the protocol. When $Y(t) = D_a(t)$, $M_1$ represents the maximal level of transcriptionally active STAT1. For downstream effectors, it represents the maximal effector level reached during the simulation, whereas for viral variables it measures the maximal viral burden.

*Integrated response.* The cumulative response is quantified by the area under the curve,

$$M_2(\mathcal{P}) := \text{AUC}(\mathcal{P}) = \int_0^{T_{\text{obs}}} Y(t)\, dt. \tag{S3.2}$$

This metric combines both amplitude and duration. It is therefore useful when two protocols generate different peak amplitudes but comparable cumulative exposure. In the case of $D_a(t)$, $M_2$ measures the integrated transcriptionally active STAT1 signal. For effector variables, it measures cumulative effector exposure, whereas for viral variables it measures cumulative viral burden.

*Time above a fixed threshold.* To quantify response persistence, we define the total time spent above a prescribed threshold $\theta$:

$$M_3(\mathcal{P}; \theta) := T_{\text{act}}(\mathcal{P}; \theta) = \text{meas}\,\{t \in [0, T_{\text{obs}}] : Y(t) \ge \theta\}, \tag{S3.3}$$

where meas denotes the length of the corresponding time set. The threshold $\theta$ is chosen from a fixed reference protocol rather than rescaled independently for each condition. A convenient choice is

$$\theta = \alpha\, Y_{\max}(\mathcal{P}_*), \qquad 0 < \alpha < 1, \tag{S3.4}$$

where $\mathcal{P}_*$ is a prescribed reference stimulation protocol and $\alpha$ is kept fixed throughout the analysis. This choice ensures that persistence is compared on a common scale across protocols.

*Time-to-peak.* The response latency is measured by the time at which the relevant peak is reached:

$$M_4(\mathcal{P}) := t_{\text{peak}}(\mathcal{P}). \tag{S3.5}$$

For a single-pulse protocol, we define

$$t_{\text{peak}}(\mathcal{P}) = \min \arg \max_{0 \le t \le T_{\text{obs}}} Y(t), \tag{S3.6}$$

that is, the first time at which the global maximum is attained. For repeated-pulse protocols, $t_{\text{peak}}$ may instead be defined pulse by pulse, depending on the objective of the analysis. In this case, $t_{\text{peak}}^{(j)}$ denotes the time of the peak associated with the $j$-th stimulation pulse.

*Post-peak persistence.* To quantify how long the response persists after its peak, we define the first post-peak threshold-crossing time

$$t_{\text{des}}(\mathcal{P}; \theta) = \inf\,\{t \ge t_{\text{peak}}(\mathcal{P}) : Y(t) \le \theta\}. \tag{S3.7}$$

The post-peak persistence is then given by

$$M_5(\mathcal{P}; \theta) := T_{\text{post}}(\mathcal{P}; \theta) = t_{\text{des}}(\mathcal{P}; \theta) - t_{\text{peak}}(\mathcal{P}). \tag{S3.8}$$

When applied to $D_a(t)$, this metric measures how long the transcriptionally active STAT1 signal remains above the fixed threshold after its maximum. More generally, it provides a termination-oriented measure of persistence for any model output.

Taken together, the metric vector

$$\mathcal{M}(\mathcal{P}) = (M_1, M_2, M_3, M_4, M_5) \tag{S3.9}$$

provides a compact summary of the trajectory $Y(t)$. The first component measures peak intensity, the second cumulative exposure, the third threshold-dependent persistence, the fourth response latency, and the fifth post-peak persistence.

*S3.2. Illustrative example using STAT1 activity*

Although the definitions above are generic, STAT1 activity provides a useful didactic example because it is directly linked to the early signalling response. In this case, we typically use

$$Y(t) = D_a(t),$$

so that the metrics quantify the amplitude, integration, timing, and persistence of the transcriptionally active STAT1 signal. This choice is particularly useful for comparing IFN protocols and *p53* regimes, because two conditions may produce similar peak values $M_1$ while differing substantially in their integrated response $M_2$ or post-peak persistence $M_5$.

For example, in the prediction analysis, $M_1$ is used to compare protocols that reach comparable STAT1 peak activity, $M_2$ is used to determine whether distinct temporal profiles produce similar cumulative activity, and $M_5$ is used to quantify whether *p53*-dependent memory prolongs the active STAT1 state after the peak. Thus, STAT1 activity is used as an illustrative readout, but the same metric definitions apply to other variables analysed throughout the manuscript.

*S3.3. Pulse-resolved metrics and recovery ratios*

For repeated stimulation protocols, global metrics can be complemented by pulse-resolved quantities. Let $Y_{\max}^{(j)}$ denote the peak value associated with the $j$-th pulse, and let $AUC^{(j)}$ denote the area under $Y(t)$ over the corresponding pulse-response window. The recovery of the response to a later pulse can then be quantified by the peak recovery ratio

$$R_j^{\text{peak}} = \frac{Y_{\max}^{(j)}}{Y_{\max}^{(1)}}, \qquad j \geq 2, \tag{S3.10}$$

or by the integrated recovery ratio

$$R_j^{\text{AUC}} = \frac{AUC^{(j)}}{AUC^{(1)}}, \qquad j \geq 2. \tag{S3.11}$$

Values close to one indicate near-complete recovery of the response, whereas values below one indicate partial refractoriness. These pulse-resolved metrics are particularly useful when analysing repeated IFN stimulation, because the first response may alter receptor availability, phosphatase activity, SOCS1 feedback, or *p53*-dependent memory, thereby changing the amplitude or persistence of subsequent responses.

*S3.4. Interpretation across model variables*

The biological interpretation of each metric depends on the chosen observable $Y(t)$. For active STAT1 variables such as $D_n(t)$ or $D_a(t)$, the metrics describe signalling amplitude, transcriptional engagement, and signal termination. For memory variables such as $A_D(t)$ or $M_{\text{IRF1}}(t)$, they quantify the accumulation and persistence of pathway history. For effector variables $P_i(t)$, they measure the strength and duration of downstream antiviral or apoptotic programmes. For viral variables such as $V_g(t)$ and $V_p(t)$, the same mathematical quantities describe viral burden rather than signalling activation; in this case, lower integrated values generally correspond to stronger antiviral control.

Thus, the metrics are used as a common mathematical framework for comparing protocols across different layers of the model, while their biological interpretation is always tied to the specific variable under consideration.

### S4. Sensitivity analysis

To assess how the implemented *p53*-dependent coupling layer modifies the control structure of the system, we compared the core JAK/STAT1 module with the coupled JAK/STAT1–*p53* model. Because memory and persistence are encoded in the *p53*-dependent coupling architecture, the aim here is not to prove their existence independently. Rather, this analysis asks how the coupling layer redistributes parameter sensitivity across distinct response features: amplitude, temporal integration, activation timing, threshold-duration commitment, and post-peak persistence.

We report first-order Sobol indices $S_1$ and total-order Sobol indices $S_T$ for each response metric. First-order indices quantify the contribution of a parameter acting alone, whereas total-order indices quantify the full contribution of that parameter, including its interactions with all other parameters. Therefore, a large difference between $S_T$ and $S_1$ indicates that the corresponding

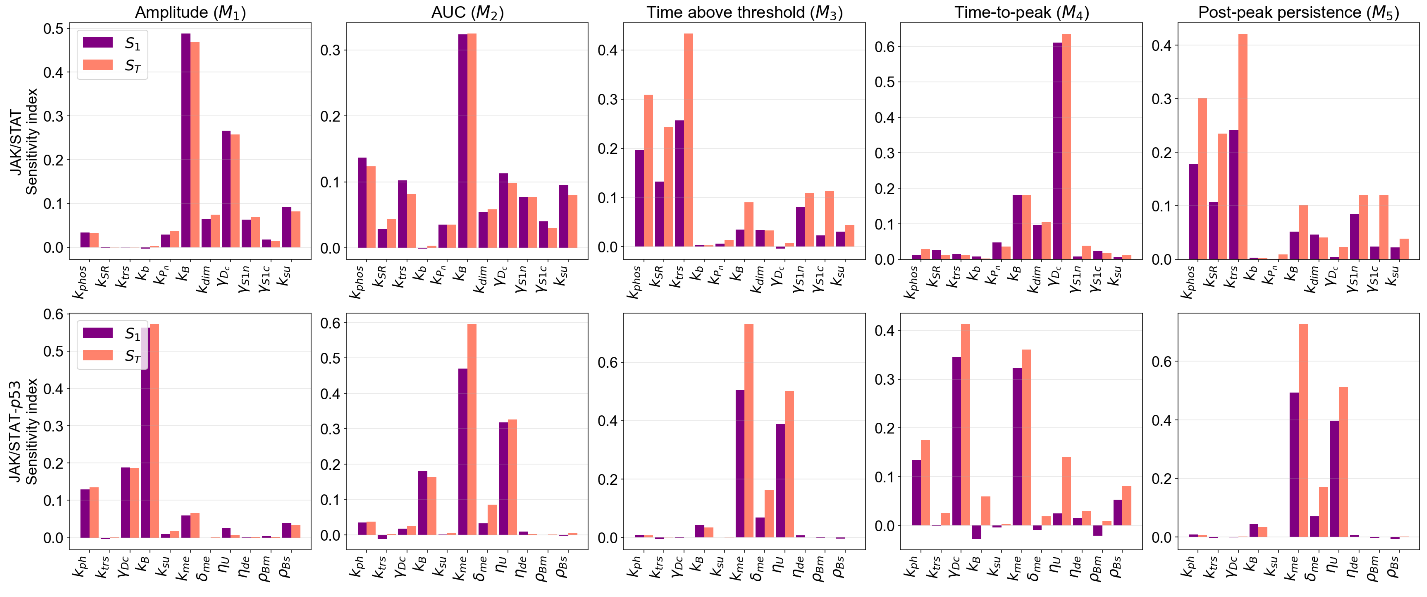


Figure S2: **Reorganisation of parameter sensitivities under *p53*-dependent coupling.** Comparison of first-order $S_1$ and total-order $S_T$ Sobol indices for the core JAK/STAT1 model (top row) and the coupled JAK/STAT1–*p53* model (bottom row). Columns correspond to the response metrics: amplitude $M_1$, integrated response $M_2$, time above threshold $M_3$, time-to-peak $M_4$, and post-peak persistence $M_5$. Purple bars show $S_1$, whereas salmon bars show $S_T$. Slightly negative first-order estimates, when present, are interpreted as numerical estimation artefacts and treated as values close to zero. Evidence for interaction-driven control is assessed from the gap between $S_T$ and $S_1$, not from the sign of $S_1$.

parameter contributes substantially through interaction terms rather than through an isolated main effect.

Figure S2 shows that the core JAK/STAT1 model and the coupled JAK/STAT1–*p53* model do not have the same sensitivity organisation. In the core model, the response metrics are mainly controlled by parameters associated with STAT1 activation, dimerisation, nuclear import, DNA-binding dynamics, and signal propagation. This is consistent with a system in which the main source of variability lies in how efficiently the IFN-induced signal is transmitted through the JAK/STAT1 module.

After inclusion of the *p53*-dependent coupling layer, the dominant sensitivities shift towards parameters associated with nuclear persistence, memory accumulation, and memory-dependent modulation. This shift is most visible for the integration and persistence-related metrics, particularly the AUC $M_2$, the time above threshold $M_3$, and the post-peak persistence $M_5$. These metrics depend less on a single activation step and more on the combined effect of DNA-bound STAT1 persistence, memory accumulation, and *p53*-dependent modulation of nuclear STAT1 dynamics.

The comparison between $S_1$ and $S_T$ further indicates that the coupled model is more interaction-driven than the core model. In several coupled-model metrics, total-order indices exceed first-order indices, showing that parameter effects are mediated by combinations of processes rather than by isolated parameters alone. This is expected in the coupled architecture, where *p53*-dependent gates modify effective DNA binding, unbinding, dephosphorylation, nuclear import, and downstream memory accumulation simultaneously.

Slightly negative first-order Sobol estimates were not interpreted as evidence for biological or dynamical antagonism. Such values can arise from numerical estimation error in variance-based sensitivity analysis and should be considered as estimates close to zero. The evidence for interaction-dominated behaviour therefore comes from the relative dominance of total-order effects and from the difference $S_T - S_1$, rather than from the negativity of individual first-order estimates.

Taken together, the sensitivity analysis supports the interpretation that the implemented *p53*-dependent coupling layer changes the parametric control of the JAK/STAT1 response. The core model is organised primarily around activation and signal-propagation parameters, whereas the coupled model becomes more sensitive to parameters controlling persistence, memory, and higher-order interactions. This result should be read as a control-structure analysis of the model architecture: once *p53*-dependent gates are introduced, the system becomes less dominated by isolated activation parameters and more governed by the coordinated regulation of nuclear persistence and transcriptional memory.

### S5. Supplementary analysis of transcriptional recovery and return to baseline

The main text uses the nuclear STAT1 dimer pool $D_n(t)$ to quantify the ability of the JAK/STAT1 pathway to restart after repeated IFN stimulation. Here, we provide complementary analyses based on the DNA-bound STAT1 state $D_a(t)$. Whereas $D_n(t)$ reports upstream pathway re-induction, $D_a(t)$ captures the recovery of the transcriptionally active nuclear STAT1 state, which depends on DNA binding, residence time, and nuclear persistence.

For a generic readout $Y(t)$, with

$$Y(t) \in \{D_n(t), D_a(t)\},$$

the incremental recovery fraction after the $k$-th pulse was defined as

$$\mathcal{R}_k^Y(g) = \frac{\max\limits_{t\in W_k(g)} \left[Y(t) - Y(t_k^-)\right]_+}{\max\limits_{t\in W_1} \left[Y(t) - Y(t_1^-)\right]_+}, \qquad [z]_+ = \max(z, 0).$$

Here, $g$ is the interpulse gap, $W_1$ is the response window following the first IFN pulse, $W_k(g)$ is the response window following the $k$-th pulse, and $Y(t_k^-)$ is the value of the readout immediately before the $k$-th stimulation. This incremental definition measures the newly induced response generated by each pulse, rather than residual activity remaining from previous stimulation.

The $D_a(t)$-based recovery analysis shows how rapidly the transcriptionally active STAT1 state becomes available for a new response. In the full model, stronger *p53* regimes are associated with slower recovery of the DNA-bound STAT1 response, especially in the plateau-like regime. The decomposition of the $S = 24$ condition shows that this delay is mainly associated with the memory component of the *p53* coupling. Thus, the same memory-dependent mechanism that prolongs DNA-bound STAT1 activity also increases the time required for the transcriptional layer to reset.

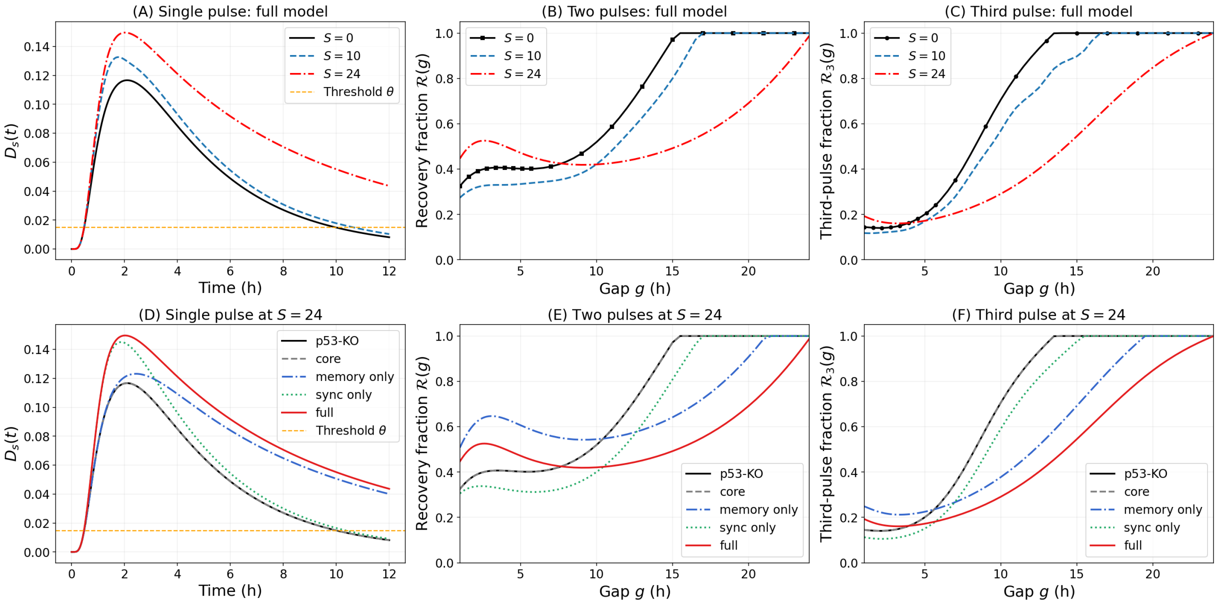


Figure S3: Recovery analysis based on the DNA-bound STAT1 readout $D_a(t)$. Panels (A)–(C) show, for the full model, the single-pulse $D_a(t)$ response, the recovery fraction after a second IFN pulse, and the recovery fraction after a third IFN pulse, respectively, under three *p53* regimes: $S = 0$, $S = 10$, and $S = 24$. Panels (D)–(F) show the corresponding analysis at $S = 24$ for the *p53*-KO, core, memory-only, synchronisation-only, and full-coupling scenarios. Recovery fractions are plotted as functions of the interpulse gap $g$ and are normalised to the single-pulse response. The orange dashed line in panels (A) and (D) indicates the threshold $\theta$ used to define the response window.

To complement the recovery fractions, we also quantified the time required for $D_a(t)$ to return below the baseline threshold after a single IFN pulse. This metric directly measures the persistence of the transcriptionally active STAT1 state. The return-to-baseline analysis confirms that plateau-like *p53* activity markedly prolongs the DNA-bound STAT1 response. In the $S = 24$ decomposition, the memory-only scenario gives the longest return-to-baseline time, whereas the synchronisation-only scenario remains close to the core response. This supports the conclusion that the slow *p53*-memory component is primarily responsible for prolonging the DNA-bound STAT1 state.

Together, these supplementary analyses support the interpretation developed in the main text. The DNA-bound STAT1 readout $D_a(t)$ shows that *p53* memory prolongs the transcriptionally active

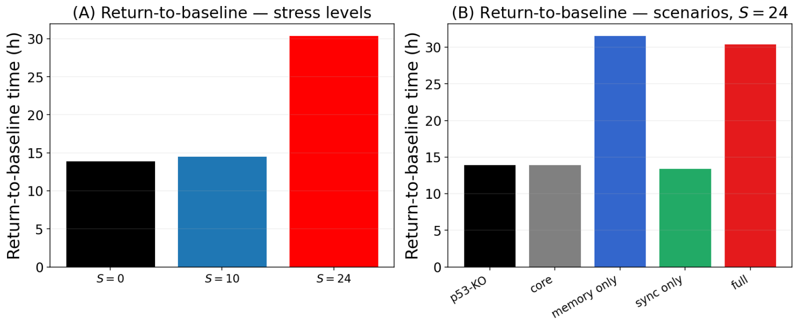


Figure S4: Return-to-baseline time of the DNA-bound STAT1 response $D_a(t)$ after a single IFN pulse. Panel (A) compares the return-to-baseline time across the three *p53* regimes $S = 0$, $S = 10$, and $S = 24$ in the full model. Panel (B) shows the corresponding decomposition of the $S = 24$ condition into *p53*-KO, core, memory-only, synchronisation-only, and full-coupling scenarios. The return-to-baseline time is reported in hours and corresponds to the time required for $D_a(t)$ to fall back below the chosen baseline threshold after the single-pulse response.

nuclear state, whereas the main-text $D_n(t)$ recovery analysis shows that this persistent state is associated with delayed pathway re-induction. Therefore, *p53* memory does not merely amplify STAT1 activity; it also reshapes the balance between signal persistence and recovery after repeated IFN stimulation.

### S6. Passive viral classes and viral-response traits

To analyse how host antiviral activation translates into functional viral control, we defined viral classes by fixed intrinsic traits rather than by explicit adaptive strategies. In the model, viruses do not actively modify host-cell parameters and do not deploy explicit counter-regulatory mechanisms. Instead, each viral class is described by two types of passive properties. The first type defines how sensitive the viral life cycle is to the antiviral effector programme. The second type defines how efficiently viral burden is converted into upstream host inputs, namely endogenous IFN production and stress-dependent *p53* activation. This separation allows us to distinguish host-side antiviral activation from realised viral control.

#### *S6.1. Effector-sensitivity classes*

The effector-sensitivity layer controls how efficiently the antiviral effectors $P_1$, $P_2$, and $P_3$ reduce viral burden. In the model, the three sensitivity parameters $\sigma_g$, $\sigma_m$, and $\sigma_a$ modulate the effect of the antiviral programme on viral genome replication, viral particle production, and viral loss, respectively. Larger values correspond to stronger viral sensitivity to the corresponding effector axis, whereas smaller values represent stronger resistance.

Table S3: Effector-sensitivity classes used to define viral susceptibility to the antiviral programme. The parameters $\sigma_g$, $\sigma_m$, and $\sigma_a$ control the sensitivity of viral genome replication, viral particle production, and viral loss to the effector layer. The intermediate class represents a transition regime in which antiviral activation can lead either to viral control or to viral persistence, depending on IFN induction and virus-induced *p53* stress.

| Viral class | $\sigma_g$ | $\sigma_m$ | $\sigma_a$ |
|---|---|---|---|
| Sensitive | 8.5 | 1.20 | 1.70 |
| Intermediate | 0.05 | 0.00208 | 0.00417 |
| Highly resistant | 0.0002 | 0.0004 | 0.0003 |

A sensitive viral class is efficiently restricted once the antiviral effectors are activated. An intermediate viral class is only partially restricted, so that antiviral activation can reduce but not necessarily clear viral burden. A highly resistant viral class remains weakly affected by the effector programme, even when the host activates several antiviral axes. Thus, the same value of $L_{\mathrm{AV}}$ can lead to different viral outcomes depending on the intrinsic effector sensitivity of the viral class.

#### *S6.2. Viral induction traits*

The viral induction layer describes how strongly viral burden generates upstream host inputs. We separated this layer into two intrinsic conversion traits: viral-to-IFN induction and viral-to-*p53* stress induction.

For endogenous IFN induction, the parameter $\chi_{\mathrm{IFN}}$ controls the efficiency with which viral burden is converted into an IFN-inducing sensing signal. The values of $\chi_{\mathrm{IFN}}$ used in the integrated simulations were calibrated to produce low, intermediate, and high realised levels of virus-induced IFN.

Table S4: Endogenous IFN induction classes used in the integrated simulations. The parameter $\chi_{\mathrm{IFN}}$ controls the viral-burden-to-IFN induction gain. The calibrated values generate low, intermediate, and high realised maxima of virus-induced $IFN_{\mathrm{int}}$.

| IFN induction class | $\chi_{\mathrm{IFN}}$ | Target maximal $IFN_{\mathrm{int}}$ | Realised maximal $IFN_{\mathrm{int}}$ |
|---|---|---|---|
| Low IFN induction | 0.022927 | 10 | $\approx 10.0$ |
| Intermediate IFN induction | 0.045369 | 25 | $\approx 25.0$ |
| High IFN induction | 0.10032 | 45 | $\approx 45.0$ |

For virus-induced *p53* activation, the parameter $\chi_{p53}$ controls how efficiently viral burden is converted into a checkpoint-stress input for the *p53* module.

Table S5: Virus-induced *p53* activation classes. The parameter $\chi_{p53}$ controls the viral-burden-to-stress induction gain, that is, the strength with which viral burden drives stress-dependent *p53* activation.

| *p53* induction class | $\chi_{p53}$ |
|---|---|
| Low *p53* induction | 0.20 |
| Intermediate *p53* induction | 0.80 |
| High *p53* induction | 1.20 |

These induction traits are independent of the effector-sensitivity traits. Therefore, a viral class can be strongly sensed but resistant to the antiviral effectors, or weakly sensed but sensitive once the effector programme is activated. This formulation separates viral immunogenicity, checkpoint-stress induction, and antiviral susceptibility without introducing explicit viral counter-regulatory mechanisms.

*S6.3. Late viral-control classification*

Functional viral control was evaluated from the late viral burden over the window $[96, 120]$ h. The late mean viral genome and particle levels were compared with the untreated reference scales $V_g^{\mathrm{ref}}$ and $V_p^{\mathrm{ref}}$, giving the genome-level and particle-level suppression scores $\varphi_g$ and $\varphi_p$. The global viral-control score was defined conservatively as

$$\varphi = \min(\varphi_g, \varphi_p).$$

Thus, strong viral control is assigned only when both viral genomes and viral particles are suppressed.

Table S6: Classification of the functional viral outcome from the global suppression score $\varphi$.

| Outcome class | Criterion |
|---|---|
| Cleared | $\varphi \geq 0.97$, or late $V_g$ below the viral floor |
| Controlled | $0.70 \leq \varphi < 0.97$ |
| Restricted | $0.30 \leq \varphi < 0.70$ |
| Evaded | $\varphi < 0.30$ |

This classification separates the host antiviral state from the realised viral outcome. The host-side antiviral state is measured by $L_{\mathrm{AV}}$, whereas the functional outcome is measured by $\varphi$. A strong antiviral state can therefore still lead to viral persistence if the viral class is poorly sensitive to the induced effectors or if suppression is discordant between $V_g$ and $V_p$.

*S6.4. Viral-burden conversion modules*

To connect viral replication with host-cell signalling, we introduced two conversion modules that translate viral burden into upstream cellular inputs. First, viral burden is converted into an IFN-inducing signal, representing the activation of viral sensing and the subsequent production of virus-induced IFN. Second, viral burden is converted into a checkpoint-activating stress signal, representing the cellular stress generated by viral replication and used to drive the *p53* module. These two conversions allow the same viral load to generate two distinct host responses: an antiviral JAK/STAT1 response through IFN, and a stress-checkpoint response through *p53*.

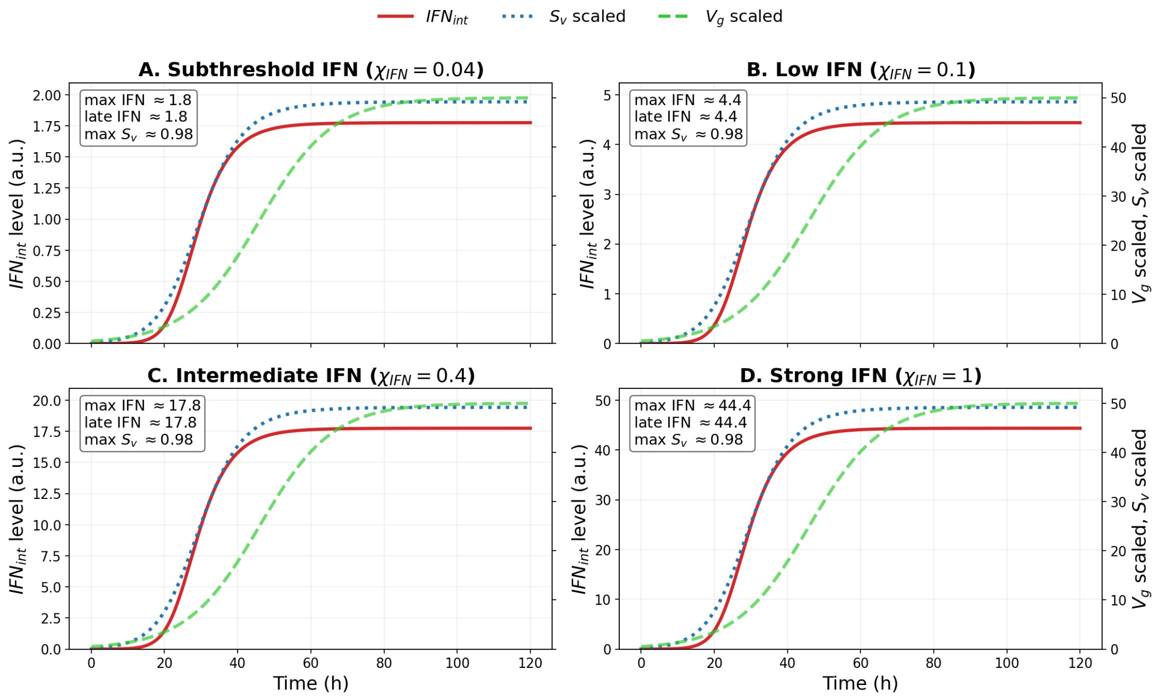


Figure S5: Calibration of the viral-burden-to-IFN conversion. Each panel represents a viral-to-IFN induction class defined by the conversion gain $\chi_{\mathrm{IFN}}$. Viral genome burden $V_g(t)$ is used as the upstream input and drives the sensing variable $S_v(t)$, which then induces virus-derived $IFN_{\mathrm{int}}(t)$. Increasing $\chi_{\mathrm{IFN}}$ progressively raises the maximal and late IFN levels while preserving the same viral-burden input structure. This calibration defines the IFN-induction benchmarks used to distinguish weak, intermediate, and strong virus-induced IFN responses.

This distinction is important because viral classes may differ not only in their sensitivity to antiviral effectors, but also in the strength with which viral burden is converted into IFN production or checkpoint activation. A viral class may strongly induce IFN while weakly activating *p53*, favouring a JAK/STAT-associated antiviral or apoptotic route. Conversely, another viral class may generate stronger checkpoint stress and robust *p53* activation, favouring a *p53*-autonomous apoptotic route. The conversion modules therefore provide a controlled way to analyse how viral burden is interpreted by the cell and how this interpretation shapes antiviral control and apoptotic commitment.

*Viral-burden-to-IFN conversion.* We calibrated the conversion from viral burden to virus-induced IFN. In this diagnostic setting, viral genome burden $V_g(t)$ is used as the upstream input, which drives the sensing variable $S_v(t)$. The sensing signal then induces $IFN_{\mathrm{int}}(t)$. The conversion gain $\chi_{\mathrm{IFN}}$ controls how efficiently viral burden is translated into endogenous IFN production. Increasing $\chi_{\mathrm{IFN}}$ therefore does not change the qualitative structure of the conversion, but expands the amplitude of the induced IFN response.

This conversion should be read as a calibration step rather than as a full antiviral-response simulation. The aim is to define how efficiently a given viral burden is translated into endogenous IFN. Low values of $\chi_{\mathrm{IFN}}$ represent infection scenarios in which viral sensing produces only a weak IFN response, whereas high values of $\chi_{\mathrm{IFN}}$ represent efficient viral sensing and strong IFN induction. These IFN-induction classes are then combined with viral-to-stress classes in the integrated simulations.

*Viral-burden-to-stress conversion.* Viral burden was converted into a checkpoint-activating stress input through a saturating viral-to-stress function. The conversion gain $\chi_{p53}$ was calibrated against the standalone *p53* module so that increasing viral-to-stress sensitivity progressively expanded the set of reachable *p53* regimes, from subthreshold activation to sustained oscillations, damped oscillations, and plateau-like activation (Figure S6).

#### *S6.5. Robustness of apoptotic-route classification to commitment thresholds*

The apoptotic-route classifications reported in the main simulations depend on a duration-dependent threshold rule. To assess whether the qualitative routing regimes are artefacts of a single threshold choice, we repeated the route classification over a grid of apoptotic thresholds. The nominal values were

$$\theta_{P4} = 30, \qquad \theta_{P5} = 40, \qquad \tau = 8\,\mathrm{h}.$$

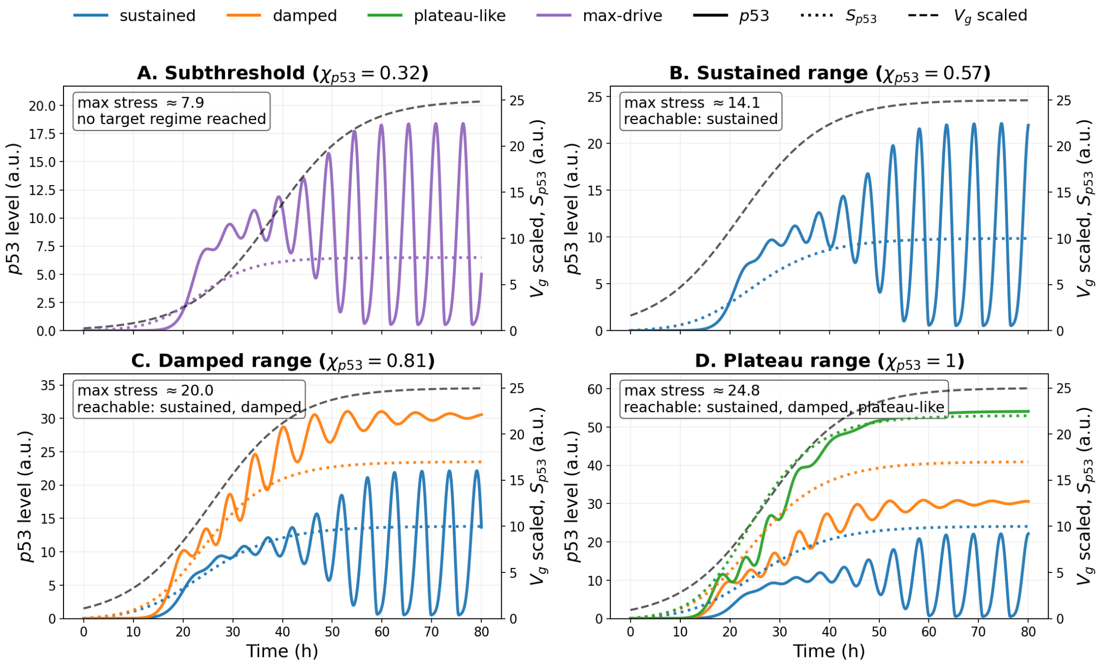


Figure S6: Calibration of the viral-burden-to-*p53*-stress conversion. Each panel corresponds to a viral-to-stress gain class $\chi_{p53}$. For each class, only the *p53* regimes reachable by the induced stress signal are shown.

We varied both apoptotic thresholds by $\pm 40\%$,

$$\theta_{P4} \in 30 \times \{0.6, 0.8, 1.0, 1.2, 1.4\}, \qquad \theta_{P5} \in 40 \times \{0.6, 0.8, 1.0, 1.2, 1.4\},$$

and repeated the classification for $\tau \in \{4, 8, 12\}$ h. The heatmaps in Figs. S7–S9 show the classifications obtained for $\tau = 8$ h, while the percentage reported inside each panel also gives the fraction of the full threshold–duration grid that preserves the nominal route. The black cross marks the nominal threshold pair. The four possible classifications are: no sustained apoptotic commitment (NONE), JAK/STAT–IRF1-associated apoptosis (APO-J), *p53*-autonomous apoptosis (APO-P), and dual-route apoptosis (APO-D).

Overall, the threshold-robustness analysis supports the qualitative routing conclusions. The sensitive viral class defines a robust non-apoptotic endpoint: for all IFN levels and all $\chi_{p53}$ values, the classification remains NONE over the full threshold–duration grid. This supports the interpretation that efficient viral restriction prevents sustained apoptotic commitment by reducing the viral-burden-dependent danger signal before the apoptotic routes can remain above threshold.

The highly resistant class defines the opposite robust endpoint. At low $\chi_{p53}$, persistent viral burden favours a JAK/STAT–IRF1-associated apoptotic route. When $\chi_{p53}$ is increased, the same persistent infection also sustains the *p53*-autonomous route, leading to dual-route commitment. The APO-D classification remains stable for most of the threshold grid, although the right-hand region of the heatmaps shows that increasing $\theta_{P5}$ can remove the *p53*-autonomous component and return the classification to APO-J. Thus, the dual-route conclusion is robust, but remains logically dependent on the threshold required for sustained *p53*-autonomous commitment.

The intermediate viral class is the least robust and should be interpreted as a transition regime. Its nominal classification can switch between APO-J and APO-D, and in some low-stress settings between NONE and APO-J, depending on the apoptotic thresholds. This threshold sensitivity is not a weakness of the analysis; rather, it indicates that intermediate viral resistance places the system close to the boundary between non-apoptotic restriction, JAK/STAT–IRF1-associated apoptosis, and dual-route apoptosis. Therefore, the sensitive and highly resistant classes provide robust endpoint behaviours, whereas the intermediate class marks the transition zone in which apoptotic routing is most dependent on the commitment criterion.

### S7. Different scenarios

In this section, we provide additional simulations supporting the main-text analysis. These scenarios test whether the memory–effector and apoptotic-routing patterns observed in the benchmark

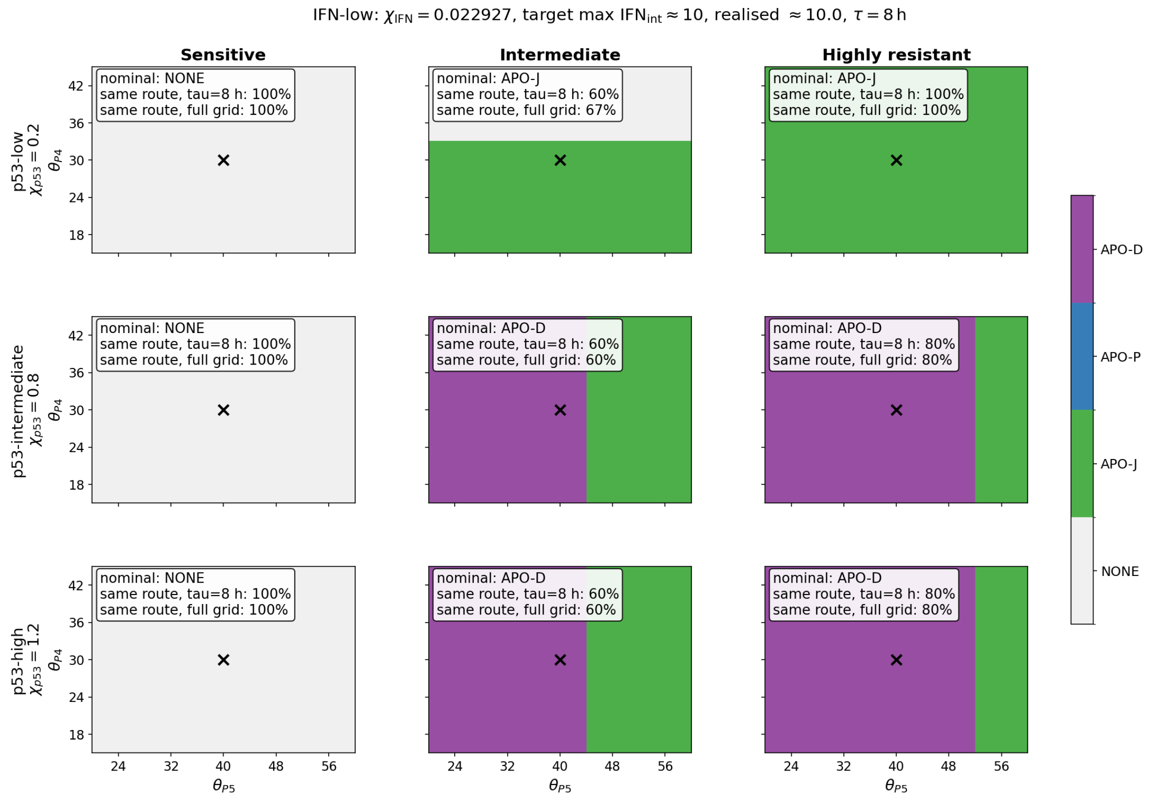


Figure S7: Robustness of apoptotic-route classification at low virus-induced IFN. The figure shows the route obtained when $\theta_{P4}$ and $\theta_{P5}$ are varied around their nominal values, with $\tau = 8\,\mathrm{h}$. Rows correspond to increasing viral-to-*p53* stress gain $\chi_{p53}$, and columns correspond to the sensitive, intermediate, and highly resistant viral classes. The sensitive class remains non-apoptotic across the threshold grid. The highly resistant class is classified as APO-J at low $\chi_{p53}$, and as APO-D at higher $\chi_{p53}$, with only partial sensitivity to the $\theta_{P5}$ threshold. The intermediate class lies closer to the classification boundary.

conditions are preserved when IFN source, viral resistance, viral-to-IFN induction and viral-to-*p53* stress conversion are varied.

#### *S7.1. Effectors in the prescribed p53 setting under dual IFN-source control*

These trajectories show that prescribed *p53* acts mainly as a gain-control layer on the antiviral programme. The plateau-like regime produces the strongest amplification of $P_1$, $P_2$, and $P_3$, whereas the oscillatory regime remains closer to the JAK/STAT1 core condition. Thus, transient *p53* activity can reinforce the antiviral response, but a sustained *p53* state is required to impose a stronger long-term effector gain.

The three effectors are not affected identically. $P_1$ behaves as a more rapid and transient antiviral component, closely following the acute IFN/JAK–STAT1 input. In contrast, $P_2$ and especially $P_3$ show more sustained accumulation, consistent with their dependence on integrated STAT1 or IRF1 memory. This indicates that *p53* does not uniformly amplify all antiviral outputs, but preferentially reinforces the more persistent arm of the effector programme.

The qualitative hierarchy is similar under virus-induced and combined IFN conditions. Once virus-induced IFN is sufficiently strong, adding an external IFN component does not fundamentally change the ordering of the effector responses. In this regime, the prescribed *p53* state mainly modulates effector amplitude, whereas IFN-source composition affects how the upstream signal is maintained.

#### *S7.2. Trajectory-level view of the dual-source IFN simulations*

Figure S11 shows the time-course counterpart of the response map in Figure 11 (main text), for an intermediate and a highly resistant viral class under prescribed *p53* regimes. The trajectories confirm the class-dependent reading summarised in the main text: the intermediate class engages the antiviral programme and improves $\varphi$, whereas the highly resistant class sustains a high $L_{\mathrm{AV}}$ with low $\varphi$, and the virus-coupled IFN component keeps the IFN–IRF1 danger signal active so that $P_4$ accumulates with delay (APO-J under oscillatory *p53*, dual-route APO-D under plateau-like *p53*). Full simulation and threshold settings are given in the figure caption.

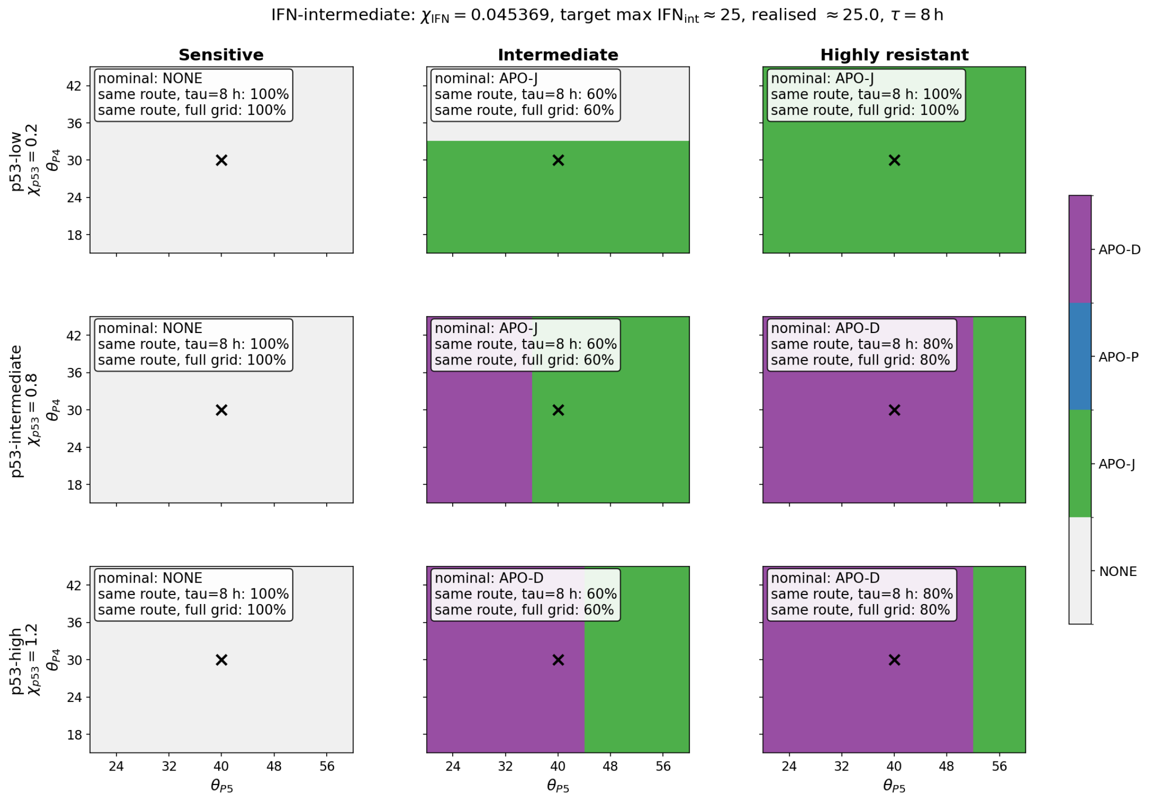


Figure S8: Robustness of apoptotic-route classification at intermediate virus-induced IFN. The same threshold scan is shown for the intermediate IFN condition. The qualitative organisation is preserved: the sensitive class remains non-apoptotic, the highly resistant class shows robust JAK/STAT–IRF1-associated commitment at low $\chi_{p53}$, and dual-route commitment at higher $\chi_{p53}$. The intermediate viral class remains the most threshold-sensitive regime, switching between APO-J and APO-D depending mainly on the $\theta_{P5}$ value.

#### *S7.3. Viral-to-stress and viral-to-IFN contributions to JAK/STAT1 antiviral states and antiviral responses*

To assess whether the memory–effector dynamics observed in the main-text benchmark depend on the infection context, we considered two additional configurations. These scenarios separate the contribution of viral resistance from the capacity of viral burden to induce IFN.

In the intermediate, strongly IFN-inducing configuration, early memory and effector responses remain similar across *p53* regimes. Separation between regimes appears mainly at later times, when persistent viral-to-stress signalling progressively reveals the modulatory effect of *p53*. This late reinforcement remains moderate because partial viral control limits the duration of the infection-coupled IFN and checkpoint-stress inputs.

In the highly resistant but weakly IFN-inducing configuration, memory and effector responses develop more gradually. Viral persistence maintains the checkpoint-stress input and allows damped and plateau-like *p53* regimes to reinforce the late response, but weak viral-to-IFN conversion limits the early JAK/STAT1-driven phase. Thus, persistent viral burden can sustain late *p53*-dependent amplification, but it cannot fully compensate for weak IFN induction.

Together, Figures S12 and S13 show that memory–effector dynamics are controlled jointly by viral resistance and viral-to-IFN conversion. Viral resistance determines how long infection-coupled inputs persist, whereas IFN-induction capacity determines how efficiently viral burden is converted into an early JAK/STAT1 response.

#### *S7.4. Dual IFN-induced sources and virus-induced p53*

We next asked whether antiviral control depends only on the presence of IFN, or also on the source from which IFN is generated. In this setting, *p53* is virus-induced, whereas IFN can arise from two sources: an externally imposed IFN input and a virus-induced IFN input generated by viral burden. This separates two questions: whether IFN-source composition alters activation of the host antiviral programme, and whether virus-induced *p53* modifies the functional outcome of this programme.

The two IFN sources activate the same JAK/STAT1 module, but they do not carry the same temporal information. External IFN imposes an antiviral drive independently of the infection state,

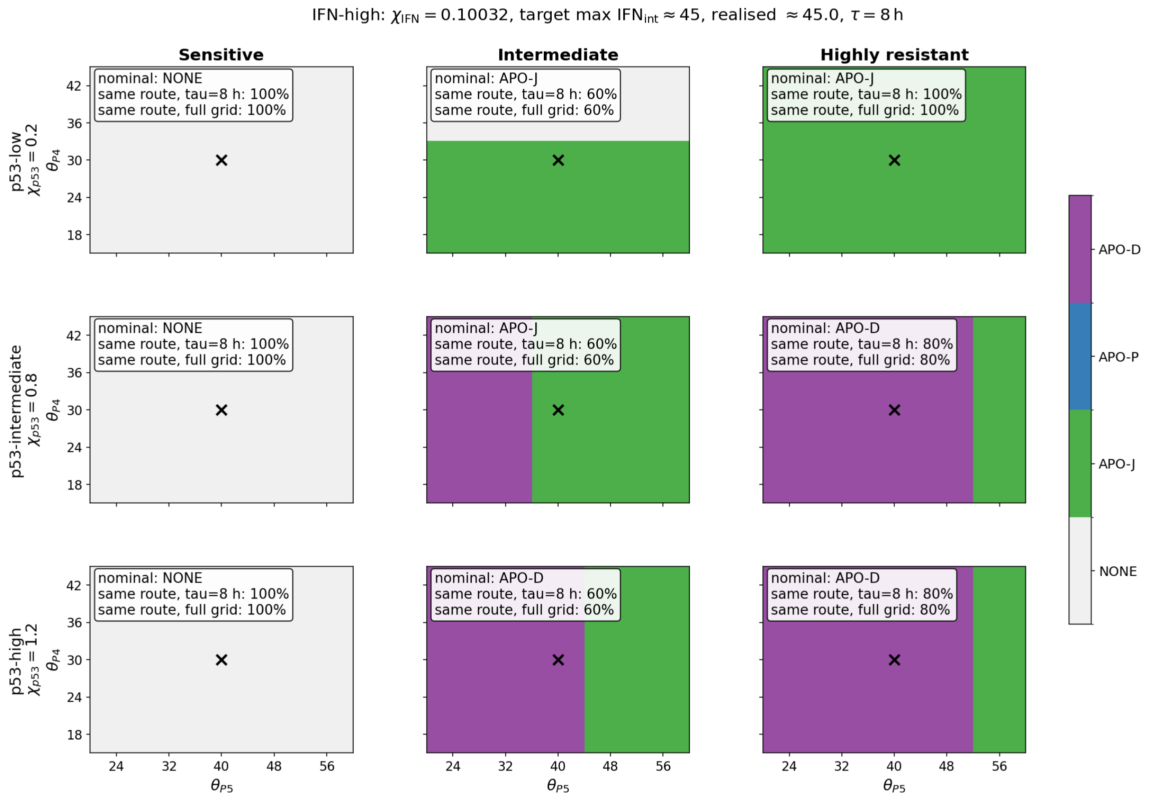


Figure S9: Robustness of apoptotic-route classification at high virus-induced IFN. Increasing the realised endogenous IFN level does not qualitatively change the threshold-robustness structure. Sensitive viral classes remain non-apoptotic over the full scan, whereas highly resistant viral classes maintain APO-J at low $\chi_{p53}$ and APO-D at intermediate and high $\chi_{p53}$. The persistence of the same routing structure across IFN levels indicates that apoptotic routing is governed primarily by viral persistence and virus-induced *p53* stress, rather than by IFN amplitude alone.

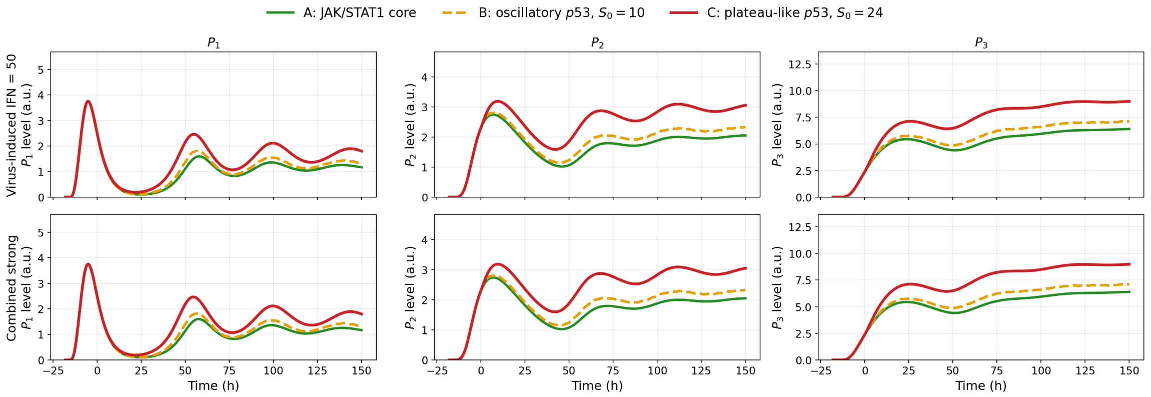


Figure S10: Antiviral effector responses under virus-induced and combined IFN inputs in the prescribed-*p53* setting. Columns show the three antiviral effectors $P_1$, $P_2$, and $P_3$. The upper row corresponds to the virus-induced IFN condition with $IFN_{max} = 50$, whereas the lower row corresponds to the combined strong IFN condition. Curves compare the JAK/STAT1 core condition, oscillatory *p53* ($S_0 = 10$), and plateau-like *p53* ($S_0 = 24$). Time is shown in hours, and effector levels are reported in arbitrary units.

whereas virus-induced IFN remains coupled to the evolving viral burden. Thus, in a dual-source setting, IFN is not only a dose variable; it also determines how strongly the host response remains linked to infection dynamics. We therefore analyse both the antiviral state, defined by sustained activation of $P_1$, $P_2$, and $P_3$, and the functional antiviral response, defined by late suppression of $V_g$ and $V_p$.

The large effector-level simulations show that virus-induced *p53* preserves the qualitative hierarchy observed in the prescribed-*p53* setting: increasing checkpoint activation strengthens the downstream antiviral programme and modulates the relative deployment of $P_1$, $P_2$, and $P_3$. The main difference appears at the apoptotic layer. When *p53* is prescribed, $P_5$ can be imposed independently of the viral trajectory. When *p53* is virus-induced, $P_5$ requires sustained viral-to-stress conversion and

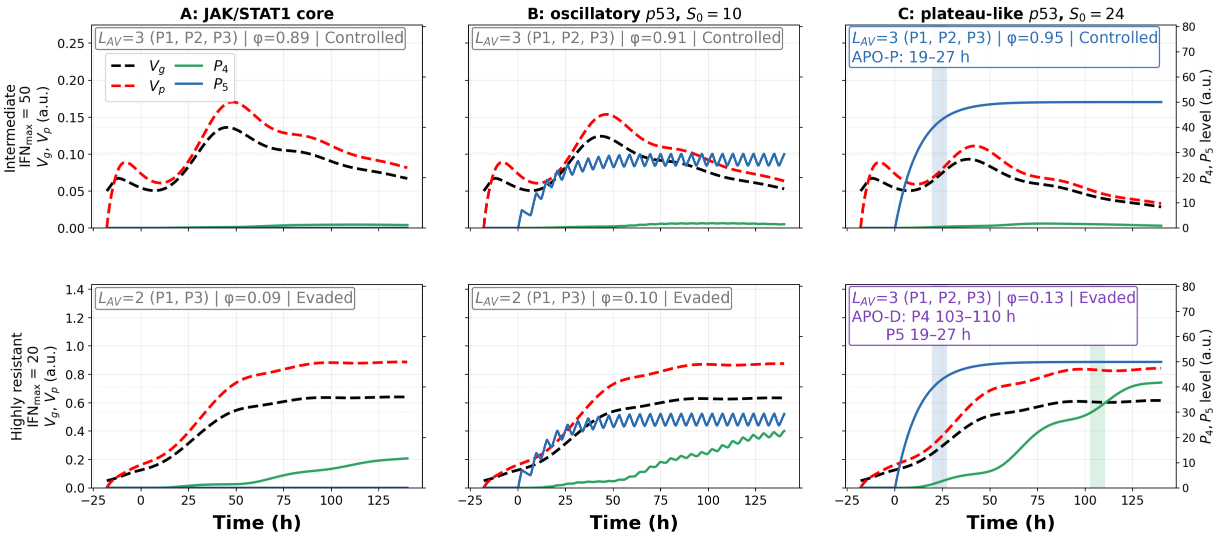


Figure S11: Dual-source IFN simulations under prescribed *p53* regimes. The total IFN input contains an externally supplied component and a virus-induced component. Rows correspond to two viral effector-sensitivity classes: an intermediate class with virus-induced IFN capacity $\mathrm{IFN}_{\max} = 50$, and a highly resistant class with virus-induced IFN capacity $\mathrm{IFN}_{\max} = 20$. Columns compare the JAK/STAT1 core condition, oscillatory *p53* ($S_0 = 10$), and plateau-like *p53* ($S_0 = 24$). Black and red dashed curves show viral genomes $V_g$ and viral particles $V_p$, respectively, on the left axis. Green and blue curves show the apoptotic outputs $P_4$ and $P_5$, respectively, on the right axis. For this scenario, apoptotic commitment was assigned using a scenario-specific $P_4$ threshold, $\theta_{P_4} = 30$, together with $\theta_{P_5} = 40$ and $\tau = 8\,\mathrm{h}$. Vertical shaded regions mark the sustained-threshold intervals used to assign apoptotic commitment. Decision boxes report the antiviral state, the viral outcome $\varphi$, and the selected apoptotic route when a commitment criterion is satisfied.

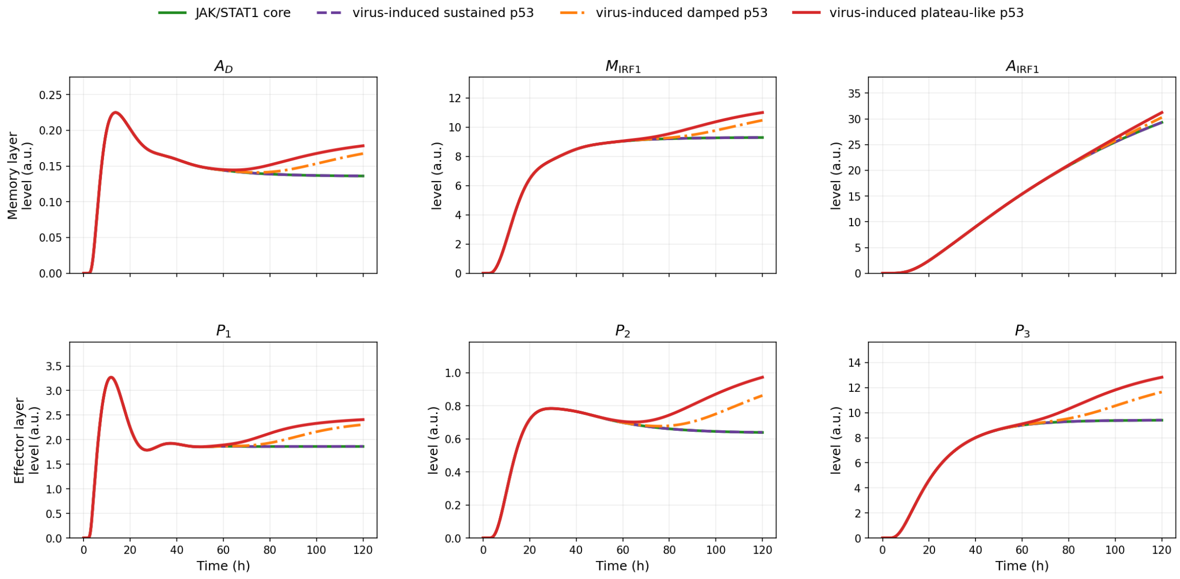


Figure S12: Memory and effector deployment under virus-induced IFN and virus-induced *p53* for an intermediate virus with high IFN-induction capacity. Curves compare the JAK/STAT1 core condition with virus-induced sustained, damped, and plateau-like *p53* regimes. The upper row shows the memory variables $A_D$, $M_{\mathrm{IRF1}}$, and $A_{\mathrm{IRF1}}$, whereas the lower row shows the antiviral effectors $P_1$, $P_2$, and $P_3$.

therefore appears mainly when viral burden generates a sufficiently strong and persistent checkpoint signal.

This shift makes apoptotic-route selection more dependent on infection dynamics. Virus-induced *p53* favours JAK/STAT–IRF1-associated $P_4$ commitment in many conditions because $P_4$ remains tied to persistent viral burden through $\Phi_{V4}(V_g)$. By contrast, $P_5$-driven commitment requires stronger and more sustained viral-to-*p53* stress induction. Thus, virus-induced *p53* does not simply amplify the prescribed-input response; it links apoptotic routing to how viral burden is partitioned between IFN induction and checkpoint stress.

We also examined how the relative contribution of external and virus-induced IFN affects late viral control under virus-induced *p53* (Figure S14). The scan shows that the most efficient antiviral

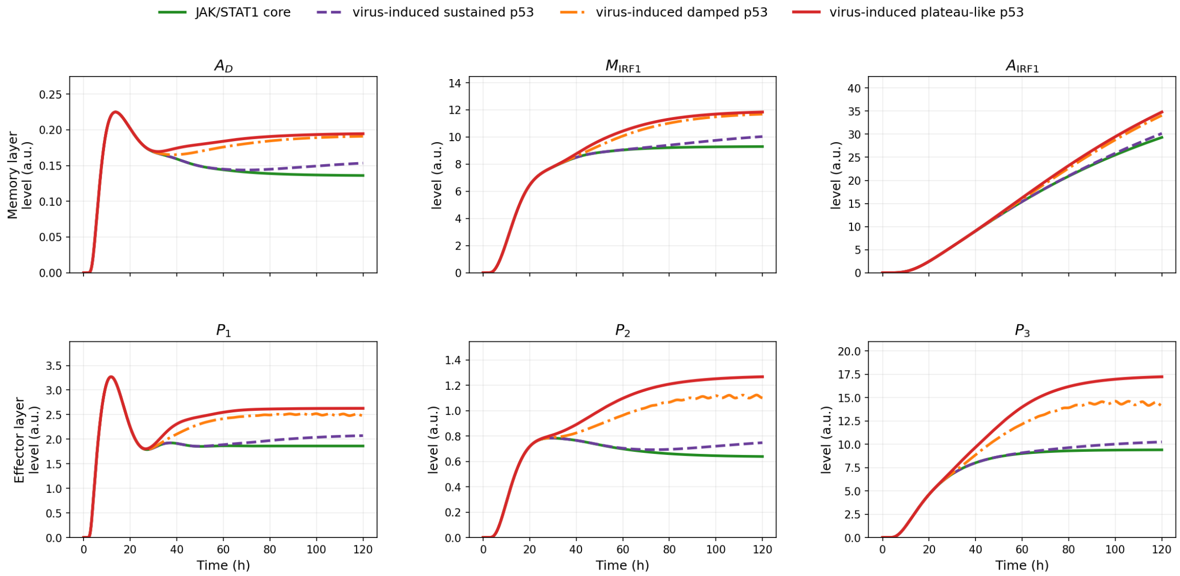


Figure S13: Memory and effector deployment under virus-induced IFN and virus-induced *p53* for a highly resistant virus with low IFN-induction capacity. Curves compare the JAK/STAT1 core condition with virus-induced sustained, damped, and plateau-like *p53* regimes. The upper row shows the memory variables $A_D$, $M_{\mathrm{IRF1}}$, and $A_{\mathrm{IRF1}}$, whereas the lower row shows the antiviral effectors $P_1$, $P_2$, and $P_3$.

control does not necessarily occur when the response is dominated by external IFN. In the tested configuration, increasing the external IFN weight reduces the virus-coupled endogenous component and can weaken late control. This indicates that virus-induced IFN carries information about infection persistence, whereas external IFN provides an imposed stimulus. The balance between the two sources therefore affects not only IFN amplitude, but also the coupling between viral persistence, checkpoint activation and apoptotic routing.

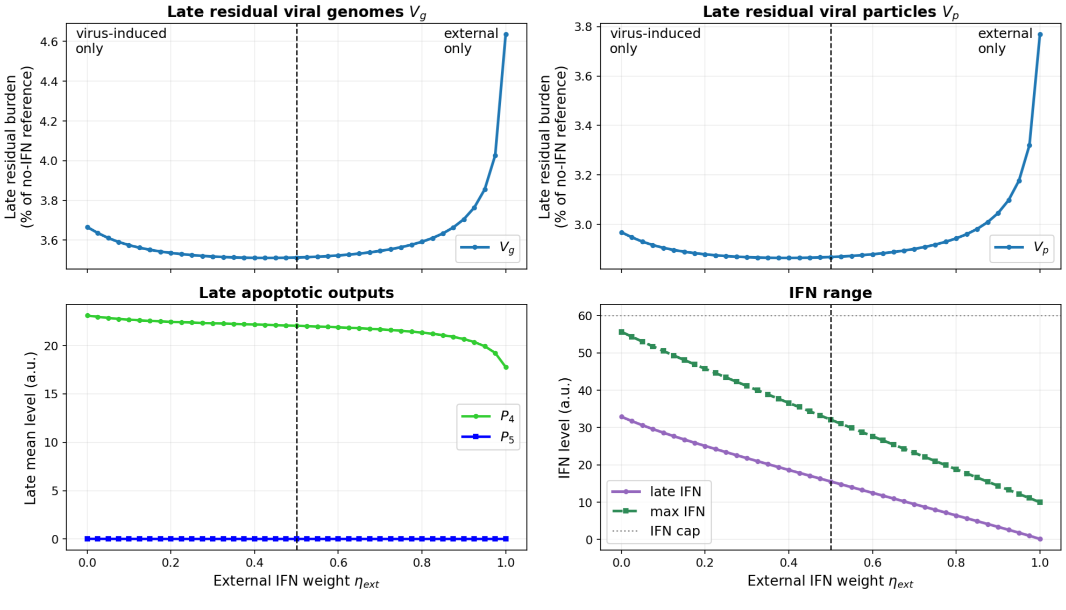


Figure S14: Effect of IFN-source composition on late viral control and apoptotic outputs under virus-induced *p53*. The external IFN weight $\eta_{\mathrm{ext}}$ is varied, with $\eta_{\mathrm{int}} = 1 - \eta_{\mathrm{ext}}$. Late residual viral genomes and particles are reported relative to the no-IFN reference, together with late $P_4$, $P_5$, and IFN levels. The scan shows how the balance between imposed and virus-coupled IFN inputs modifies late viral control and the apoptotic-output landscape.

#### *S7.5. Response-map view of the fully virus-coupled IFN–p53 simulations*

Figure S15 shows the response-map counterpart of the time-course simulations in Figure 14 (main text), generalising them across the continuous viral-to-IFN gain $\chi_{\mathrm{IFN}}$ and the viral-to-*p53* stress gain $\chi_{p53}$. It confirms the class-dependent routing described in the main text: the sensitive class stays non-apoptotic across the scanned range, the highly resistant class keeps the $V_g$-gated $P_4$ route

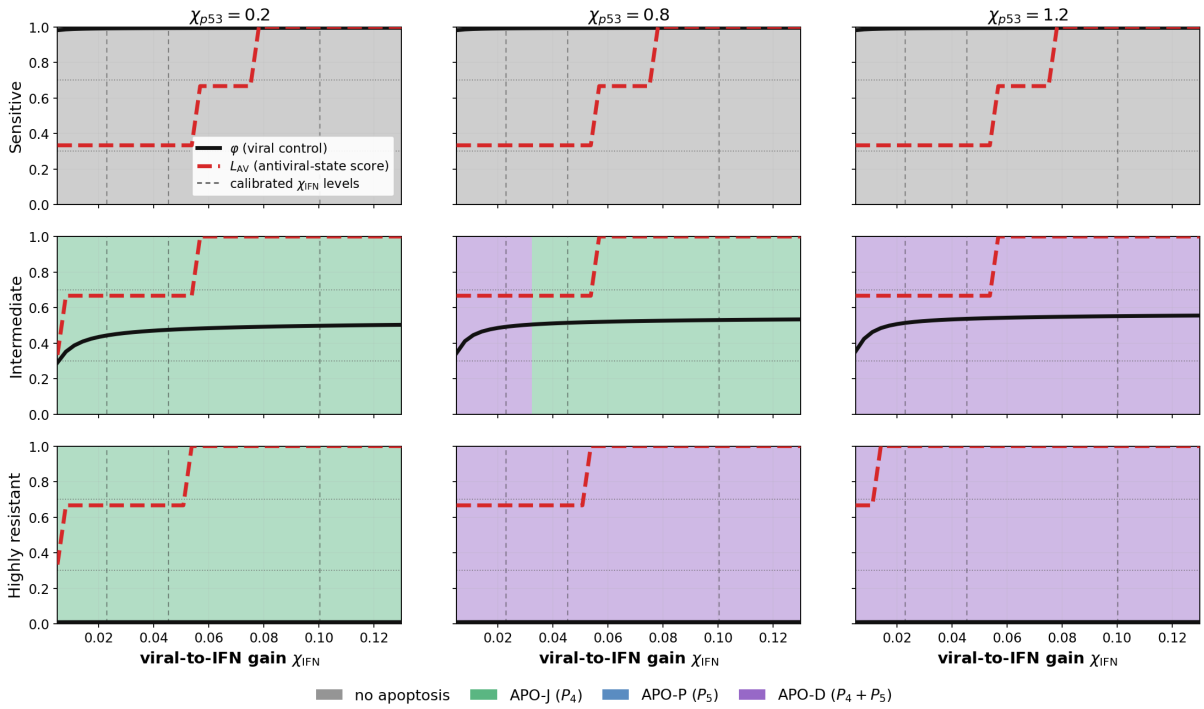


Figure S15: Response-map summary for the fully virus-coupled IFN–*p53* setting. Rows correspond to viral classes: sensitive, intermediate, and highly resistant. Columns correspond to increasing virus-induced *p53* stress, parameterised by $\chi_{p53}$. In each panel, the horizontal axis gives the viral-to-IFN sensing gain $\chi_{\mathrm{IFN}}$. Background colours indicate the apoptotic route selected at each value of $\chi_{\mathrm{IFN}}$: no apoptosis, APO-J for $P_4$-associated commitment, APO-P for $P_5$-associated commitment, and APO-D for dual $P_4 + P_5$ commitment. The same apoptotic-route colour code is used as in the decision boxes of the time-course figures. The solid black curve shows the viral-control score $\varphi$, whereas the dashed red curve shows the antiviral-state score $L_{\mathrm{AV}}$, with discrete levels 0, 1/3, 2/3, and 1. Horizontal dotted lines mark reference levels used to interpret the normalised readouts, and vertical dashed lines indicate the calibrated non-saturating $\chi_{\mathrm{IFN}}$ values used in the corresponding time-course simulations.

active over a broad range of $\chi_{\mathrm{IFN}}$ and shifts towards APO-D as $\chi_{p53}$ increases, and the intermediate class occupies the transition zone in which delayed $P_4$-commitment combines with $P_5$ recruitment. Full readout definitions and calibrated $\chi_{\mathrm{IFN}}$ values are given in the figure caption.

### S8. Gate-parameter robustness analysis

The *p53*-dependent coupling gates are phenomenological coarse-grained functions, and their individual parameters are not identifiable separately from currently available quantitative data. Robustness to gate-parameter variation is therefore not an additional check performed after calibration, but the central validity criterion for the qualitative conclusions of the model. A conclusion is retained only if it persists across broad, biologically admissible variation of the memory, synchronisation, effector-modulation, apoptotic-competence, and viral-burden-gating parameters.

This analysis was not designed to show that the gate parameters have no effect. On the contrary, these parameters are expected to control the magnitude and timing of the coupling. The purpose was instead to determine whether the main conclusions depended on a narrow nominal calibration of the *p53*-dependent gates.

We varied parameters associated with the memory gate $\Phi_{\mathrm{mem}}$, the synchronisation gate $\Phi_{\mathrm{sync}}$, the effector-modulation gate $\Phi_{p53}$, the apoptotic-competence gate $\Phi_{\mathrm{apop}}$, and the viral-burden gate $\Phi_{V4}(V_g)$. Strictly positive scale parameters were sampled around their nominal values, whereas fractional or bounded parameters were sampled within admissible intervals that preserved positive effective rates and biologically interpretable gate strengths. The viral-burden dependence of the $P_4$ route was kept explicit in all samples. Thus, the analysis varied the sensitivity and shape of the $P_4$ viral-burden gate, but did not remove the dependence of $P_4$ on $V_g$.

For each sampled parameter set, we recomputed the relevant signalling, effector, viral-control, and apoptotic-routing readouts. We focused on five qualitative conclusions: the redistribution of STAT1 signalling towards DNA-bound persistence and transcriptional memory, delayed JAK/STAT1 recovery under stronger *p53* regimes, buffering of *p53* preactivation at the effector layer, dissociation between antiviral-state engagement and viral control, and persistence-dependent apoptotic-route classification.

#### S8.1. Robustness of signalling-level redistribution

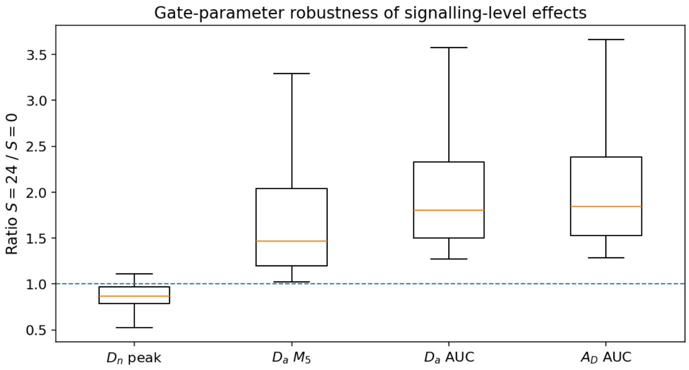


Figure S16: Gate-parameter robustness of signalling-level effects. For each sampled parameter set, response metrics were computed under the plateau-like *p53* regime ($S = 24$) and normalised to the no-*p53* condition ($S = 0$). Boxplots show the resulting ratios for the peak nuclear phosphorylated STAT1 dimer pool $D_n$, the post-peak persistence metric $M_5$ of DNA-bound STAT1 $D_a$, the AUC of $D_a$, and the AUC of the transcriptional memory variable $A_D$. The dashed horizontal line marks a ratio of one, corresponding to no change relative to the $S = 0$ condition.

Figure S16 tests whether the signalling-level interpretation is preserved when the *p53*-dependent gate parameters are varied. The peak of the nuclear phosphorylated STAT1 dimer pool $D_n$ remains close to, or below, the no-*p53* reference across the sampled parameter sets. By contrast, DNA-bound STAT1 persistence ($D_a M_5$), integrated DNA-bound STAT1 activity ($D_a$ AUC), and transcriptional memory ($A_D$ AUC) remain consistently above one.

This confirms that the qualitative signalling-level conclusion is not restricted to the nominal gate calibration. Plateau-like *p53* does not primarily increase the upstream nuclear STAT1 dimer pool. Instead, under broad gate-parameter variation, the response remains redistributed towards DNA-bound STAT1 persistence, integrated DNA-bound activity, and transcriptional memory. The variability of the boxplots shows that the gate parameters do affect the magnitude of this redistribution, but not its qualitative direction.

#### S8.2. Robustness of delayed JAK/STAT1 recovery

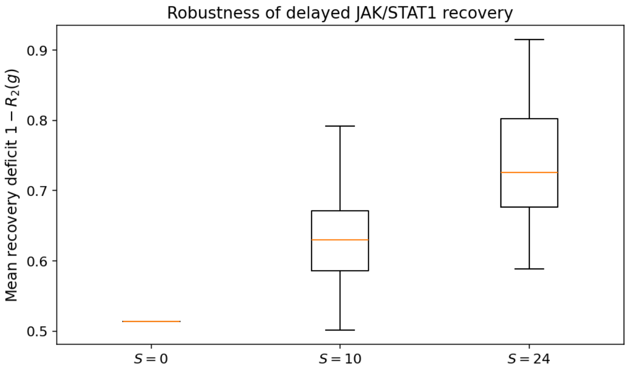


Figure S17: Robustness of delayed JAK/STAT1 recovery under gate-parameter variation. For each sampled parameter set and each *p53* regime, the mean recovery deficit $1 - R_2(g)$ was computed from the second-pulse recovery fraction $R_2(g)$. Larger values indicate poorer recovery and stronger refractoriness after repeated IFN stimulation. Boxplots compare the no-*p53* condition ($S = 0$), the oscillatory *p53* regime ($S = 10$), and the plateau-like *p53* regime ($S = 24$).

Figure S17 shows that the recovery hierarchy is preserved under gate-parameter uncertainty. The recovery deficit is lowest in the no-*p53* condition, intermediate in the oscillatory *p53* regime, and highest in the plateau-like *p53* regime. Thus, stronger *p53*-dependent memory is associated with delayed recovery after repeated IFN stimulation across the sampled parameter sets.

This supports the interpretation that *p53* introduces a persistence–recovery trade-off. The same coupling architecture that prolongs DNA-bound STAT1 activity and transcriptional memory also makes the pathway less readily re-inducible at short interpulse gaps. This behaviour is not a simple statement that one gate increases one variable; it arises from the interaction between memory accumulation, pathway reset, and repeated IFN stimulation.

#### *S8.3. Robustness of p53-preactivation buffering*

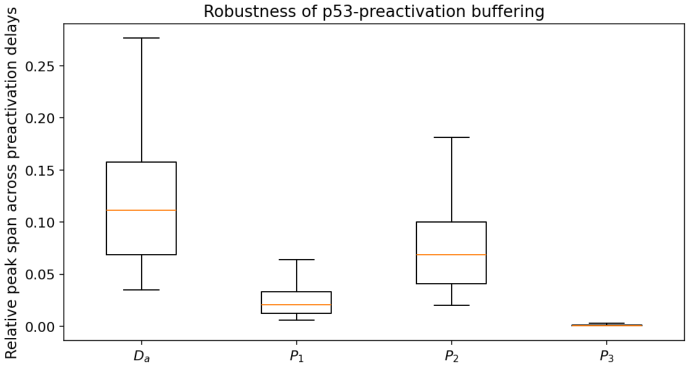


Figure S18: Robustness of *p53*-preactivation buffering under gate-parameter variation. For each sampled parameter set, the relative peak span across *p53*-preactivation delays was computed for DNA-bound STAT1 $D_a$ and for the antiviral effectors $P_1$, $P_2$, and $P_3$. Larger values indicate stronger sensitivity to the duration of *p53* preactivation, whereas smaller values indicate stronger buffering across preactivation delays.

Figure S18 tests whether the effect of *p53* preactivation is transmitted proportionally from the signalling layer to the effector layer. Across the sampled gate-parameter sets, $D_a$ shows the largest relative peak span across preactivation delays. In contrast, the antiviral effectors show a smaller spread, especially $P_1$ and $P_3$, with $P_2$ retaining an intermediate sensitivity.

Thus, the buffering conclusion is preserved under parameter uncertainty. *p53* preactivation can modify the upstream DNA-bound STAT1 response, but this effect is not transmitted linearly to all downstream antiviral effectors. The memory–effector layer filters the upstream timing differences through IRF1-associated memory, effector production, and effector turnover.

#### *S8.4. Robustness of the dissociation between antiviral-state engagement and viral control*

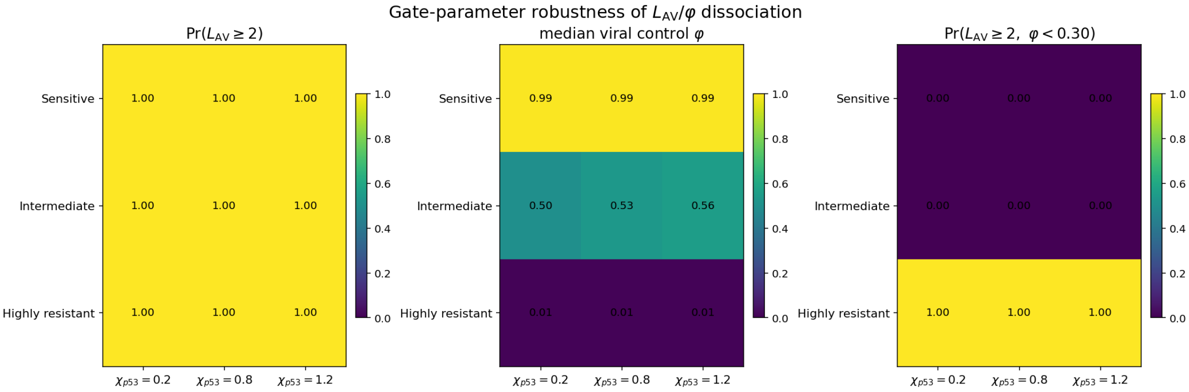


Figure S19: Robustness of the dissociation between antiviral-state engagement and functional viral control under gate-parameter variation. The left panel shows the proportion of sampled parameter sets for which the antiviral score satisfies $L_{\mathrm{AV}} \geq \frac{2}{3}$. The middle panel shows the median viral-control score $\varphi$. The right panel shows the proportion of samples for which a moderate or strong antiviral state coexists with poor viral control, defined as $L_{\mathrm{AV}} \geq \frac{2}{3}$ and $\varphi < 0.30$. Rows correspond to viral effector-sensitivity classes and columns to the viral-to-*p53* stress gain $\chi_{p53}$. Values indicate proportions across sampled gate-parameter sets, except for the middle panel, which reports the median value of $\varphi$.

Figure S19 directly tests whether antiviral-state engagement and functional viral control remain separable under gate-parameter uncertainty. Across all viral classes and all values of $\chi_{p53}$, the probability of reaching at least a moderate antiviral state satisfies $\Pr(L_{\mathrm{AV}} \geq \frac{2}{3}) = 1$. Thus, in these simulations, broad variation of the gate parameters does not abolish engagement of the antiviral programme.

However, viral control remains strongly class-dependent. Sensitive viral classes show near-complete control, with median $\varphi \simeq 0.99$. Intermediate viral classes show partial restriction, with median $\varphi$ around 0.50–0.56. Highly resistant viral classes remain poorly controlled, with median $\varphi \simeq 0.01$, despite robust antiviral-state engagement. Consequently, the dissociated outcome

$$L_{\mathrm{AV}} \geq \frac{2}{3}, \qquad \varphi < 0.30,$$

occurs systematically in the highly resistant class, but not in the sensitive or intermediate classes.

This confirms that the separation between $L_{\mathrm{AV}}$ and $\varphi$ is robust to gate-parameter variation. The failure of viral control in the highly resistant class is not due to failure to engage the antiviral programme, but to the poor sensitivity of that viral class to the induced antiviral effectors. Thus, $L_{\mathrm{AV}}$ and $\varphi$ quantify distinct levels of the response: host antiviral-state engagement and functional viral control.
